# Gene-Embedded Multi-Modal Networks for Population-Scale Multi-Omics Discovery

**DOI:** 10.1101/2025.01.22.634403

**Authors:** Vaha Akbary Moghaddam, Sandeep Acharya, Michaela Schwaiger-Haber, Shu Liao, Wooseok J. Jung, Bharat Thyagarajan, Leah P. Shriver, E. Warwick Daw, Nancy L. Saccone, Ping An, Michael R. Brent, Gary J. Patti, Michael A. Province

**Affiliations:** Department of Genetics, School of Medicine, Washington University in St. Louis, MO, USA; Division of Computational & Data Sciences, McKelvey School of Engineering, Washington University in St. Louis, MO, USA; Department of Chemistry, School of Arts & Sciences, Washington University in St. Louis, MO, USA; Center for Mass Spectrometry and Metabolic Tracing, School of Medicine, Washington University in St. Louis, MO, USA; Department of Computer Science & Engineering, McKelvey School of Engineering, Washington University in St. Louis, MO, USA; Department of Laboratory Medicine & Pathology, School of Medicine, University of Minnesota, MN, USA

## Abstract

We present Gene-Embedded Multi-modal Networks (GEM-Net), a semi-supervised framework for constructing multi-modal networks centered on genes. GEM-Net uses gene-level modules and selectively incorporates heterogeneous omics profiles using a correlated meta-analysis strategy that accounts for scale imbalance, missingness, and intra-modular correlation. Prior to network inference, we developed a harmonized data processing protocol that adjusts each omic layer independently through a shared mathematical workflow involving transformation, dimensionality reduction, and regression-based covariate adjustment. GEM-Net modules were inferred and benchmarked against unsupervised methods using transcriptomic, metabolomic, and lipidomic data from the Long Life Family Study (LLFS), a unique cohort enriched for exceptional familial longevity and health. GEM-Net modules were more diverse and biologically interpretable, with stronger support from protein– protein interactions, transcriptional regulation, and metabolic annotations. Applying GEM-Net to metabolic health in LLFS revealed an axis between the microbiome-derived metabolite N-acetylglycine and immune genes (*FCER1A, HDC, CPA3, MS4A2*) associated with improved insulin sensitivity and reduced inflammation in healthy older individuals. GEM-Nets offer a reusable reference from a long-lived population and a generalizable framework for multi-omics discovery. https://doi.org/10.5281/zenodo.15003731.

## 1. Introduction

Metabolism underlies nearly every aspect of human physiology, driving the biochemical reactions that sustain life. It is orchestrated by a vast, interconnected network of genes, proteins, and small molecules (SMs), encompassing thousands of interactions that maintain homeostasis^1^. Metabolic reactions also play a central role in determining how organisms adapt to internal and external signals throughout the lifespan. Dysregulation in core metabolic pathways has been linked to chronic inflammation, impaired insulin signaling, and increased obesity—defining features of aging-related decline^2,3^

Given the complexity of metabolism driven by the interactions across diverse molecular features, individuals can exhibit substantial variability in their metabolic phenotypes. For instance, the NIA’s Long Life Family Study (LLFS), a multi-generational and multi-national cohort enriched for exceptional longevity, consists primarily of older adults with a mean age of 70^4^. Despite this advanced age and an average body mass index (BMI) in the obese range, LLFS participants exhibit lower insulin resistance than the general non-diabetic U.S. population^5^. They also show significanttly reduced rates of diabetes and peripheral artery disease compared to other aging cohorts^6^. These patterns imply diverse, context-dependent regulatory mechanisms that defy general expectations about metabolic decline in aging.

In recent years, untargeted metabolomics has expanded our view of SMs, revealing their diverse roles in metabolism, gene expression regulation, and homeostasis^7^. Standard population-level approaches— such as metabolome-wide association studies (MWAS)—identify individual omic–trait associations but offer limited insight into broader regulatory mechanisms^8^. Functional interpretation is often pursued through metabolic enrichment analysis, grouping omics features into biologically meaningful sets^9^. While this adds a systems-level perspective, enrichment analysis is fundamentally limited to well-annotated pathways. However, many chemically characterized human SMs still lack functional annotations, restricting discovery beyond known biology^10^.

To move beyond association testing and support pathway discovery, multi-omics integration and network inference offer a powerful alternative. Network-based approaches capture coordinated patterns across omics layers by grouping related features—such as genes, SMs, and epigenetic features—into modules^11, 12^. Yet, integrating diverse omics profiles remains technically challenging due to differences in data modality, missingness, and scale imbalance^13,4^ For example, RNA-seq typically captures over 10,000 transcripts. Meanwhile liquid chromatography / mass spectrometry (LC/MS) detects only ∼1,000 SMs, with distinct distributions and noise characteristics compared to RNA-seq^15^. These disparities can lead to model overfitting, spurious associations, and often lack of meaningful integration.

Here, we present Gene-Embedded Multi-modal Networks (GEM-Net): A generalizable semi-supervised framework for constructing multi-modal networks from heterogeneous omics data. The method begins with a gene-level network—either data-driven or knowledge-guided—which is clustered into modules.

These gene-level modules serve as the input units for selectively integrating additional omics layers using a correlated meta-analysis (CMA) strategy to construct the final multi-modal networks. The CMA strategy accounts for co-expression structure, scale imbalance, and missingness across multiple omics profiles. It also corrects for inflation of type-I error due to non-independences in the data when the omics profiles are generated on the same population. We apply this framework to RNA-seq and LC/MS lipidomics and metabolomics profiles to construct Transcriptome–Small Molecule Networks.

We apply the framework to the LLFS, one of the largest human multi-omics cohorts to date^4^, and benchmark its performance against well-established unsupervised module identification methods. The resulting multi-modal modules show improved biological coherence and stronger support from known molecular interactions and associations sets. While demonstrated on RNA-seq and LC/MS data, the framework is broadly applicable to other omics layers such as proteomics or DNA methylation profiles. The inferred GEM-Nets are publicly available as a generalizable resource for multi-omics investigations across diverse biological contexts.

## 2. Results

### 2.1 Overview of GEM-Net Framework

To bridge the gap between untargeted metabolomics/lipidomics and transcriptomics in complex trait studies, we developed a scalable, semi-supervised framework for multi-modal network construction and interpretation (Figure 1). The framework proceeds in three stages:

**Figure 1.**
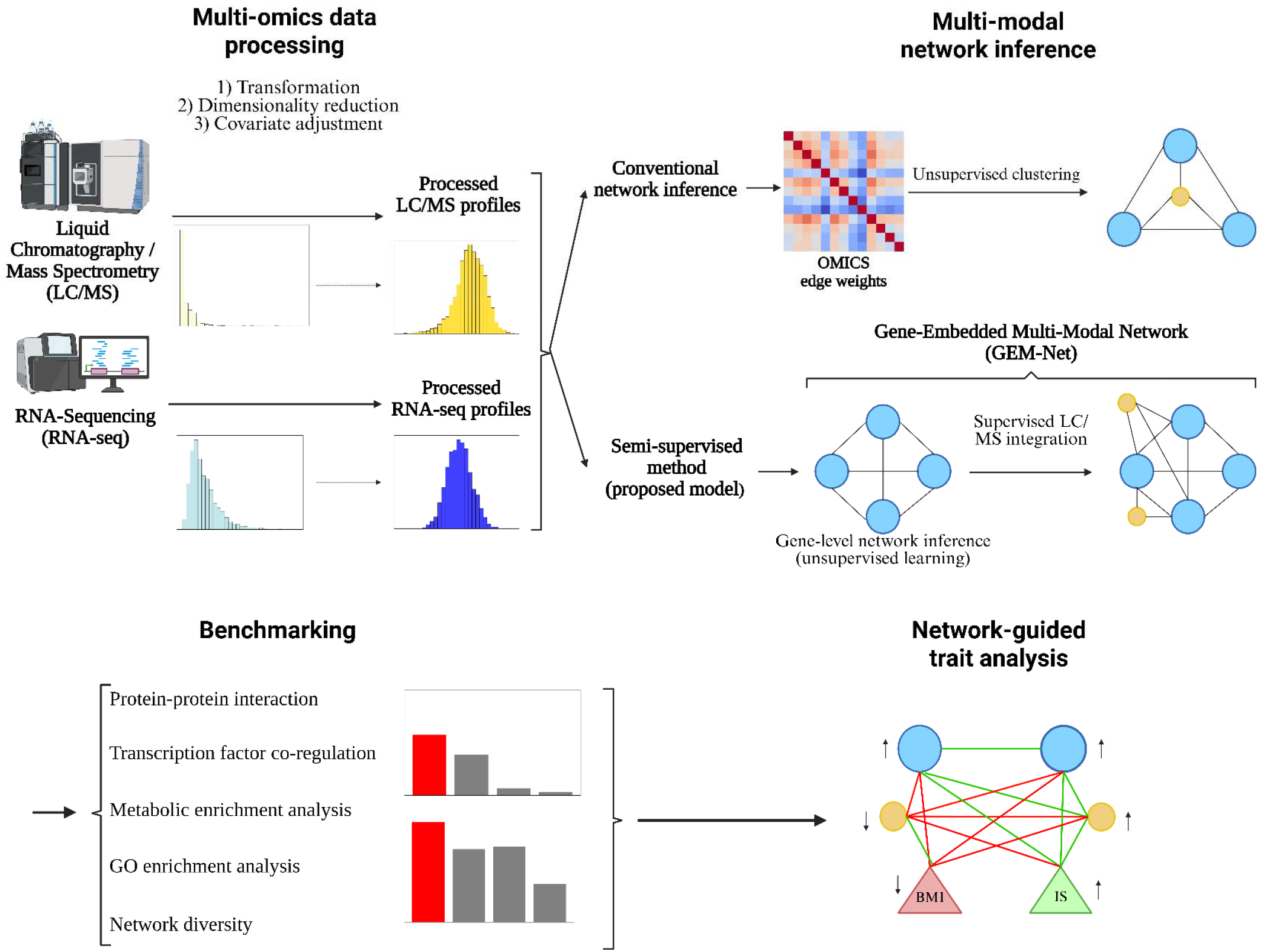
General workflow of the study. In the first section of the study, a ML-based protocol is proposed, composed of data transformation and regression-based adjustments to prepare the LC/MS and RNA-seq profiles for integration. In the second part, conventional unsupervised network inference approaches as well as a newly proposed semi-supervised method—GEM-Net—were used to construct multi-modal network. The GEM Net framework includes unsupervised construction of gene-level networks from the RNA-seq profiles with the conventional approaches, followed by supervise integration of additional omics profiles into the baseline network. GEM-NET is benchmarked against the conventional approaches using multiple evaluation metrics. Upon benchmarking the resulting GEM-NETs, they were used to study metabolic traits, such as IS or BMI, in a network-guided manner.

#### 1. Multi-omics data harmonization

We first developed a machine learning (ML)–based protocol to harmonize untargeted LC/MS and RNA-seq profiles. While data from each modality is processed independently using algorithms suited to its characteristics, the protocol follows a shared mathematical workflow comprising data transformation, dimensionality reduction, and regression-based covariate adjustment. This data harmonization structure improves the compatibility of multi-omics profiles for downstream integration and effectively accounts for technical, demographical, and biological confounders.

#### 2. Multi-modal network inference and module identification

Using the processed data, we benchmarked unsupervised module identification approaches against our proposed GEM-Net framework. Our method first infers gene-level modules from RNA-seq data and then selectively integrates additional omics data (metabolomics and lipidomics) into these modules using a CMA strategy to build GEM-Net. This two-step design addresses key integration challenges for multi-modal networks. Dividing the process into unsupervised module detection followed by supervised omics integration mitigates scale imbalance between different omics layers. In addition, CMA combines gene–SM associations within modules in a manner that accounts for not only missing data, but also intra-module correlation of gene expression—unlike traditional meta-analysis or association tests.

#### 3. Trait-focused application of GEM-Net

To demonstrate the framework’s utility, we applied the inferred GEM-Net to study metabolic health indicators in the LLFS cohort, including insulin sensitivity (IS), BMI, triglycerides (TG), and IL6. Network-guided analysis and module association tests uncovered a novel regulatory axis linking fiber-degrading gut metabolite and immune genes with improved metabolic outcomes.

### 2.2 Enhanced multi-omics integration with LC/MS data processing protocol

#### 2.2.1 Data distribution

The proposed LC/MS protocol is designed to account for technical, demographic, and biological confounders in multiple steps (Section 5.2), resulting in marked improvement in data quality and integration with the RNA-seq profiles and complex trait analysis. Raw LC/MS peak areas initially exhibited highly skewed, exponential distributions with high kurtosis and standard deviation (SD)(Figure 2a). Through batch correction, principal component analysis, and regression-based adjustments, the SM peak residuals approximated normality, with reduced kurtosis and SD across most features (Figure 2b). Importantly, the processed SM distributions aligned closely with the distribution profiles of RNA-seq data and complex traits, despite independent data processing (Figures 2c & 2d). This alignment is critical to prevent spurious signals during data integration and network inference^16^. Similar improvements were observed for SMs in the lipid category (Supplementary Figure 1).While batch effect correction reduced the SD of raw peak areas, it was insufficient for achieving full harmonization with RNA-seq data (Supplementary Figure 1). Accounting for demographic and biological confounders played a pivotal role in achieving high-quality integrable data.

**Figure 2.**
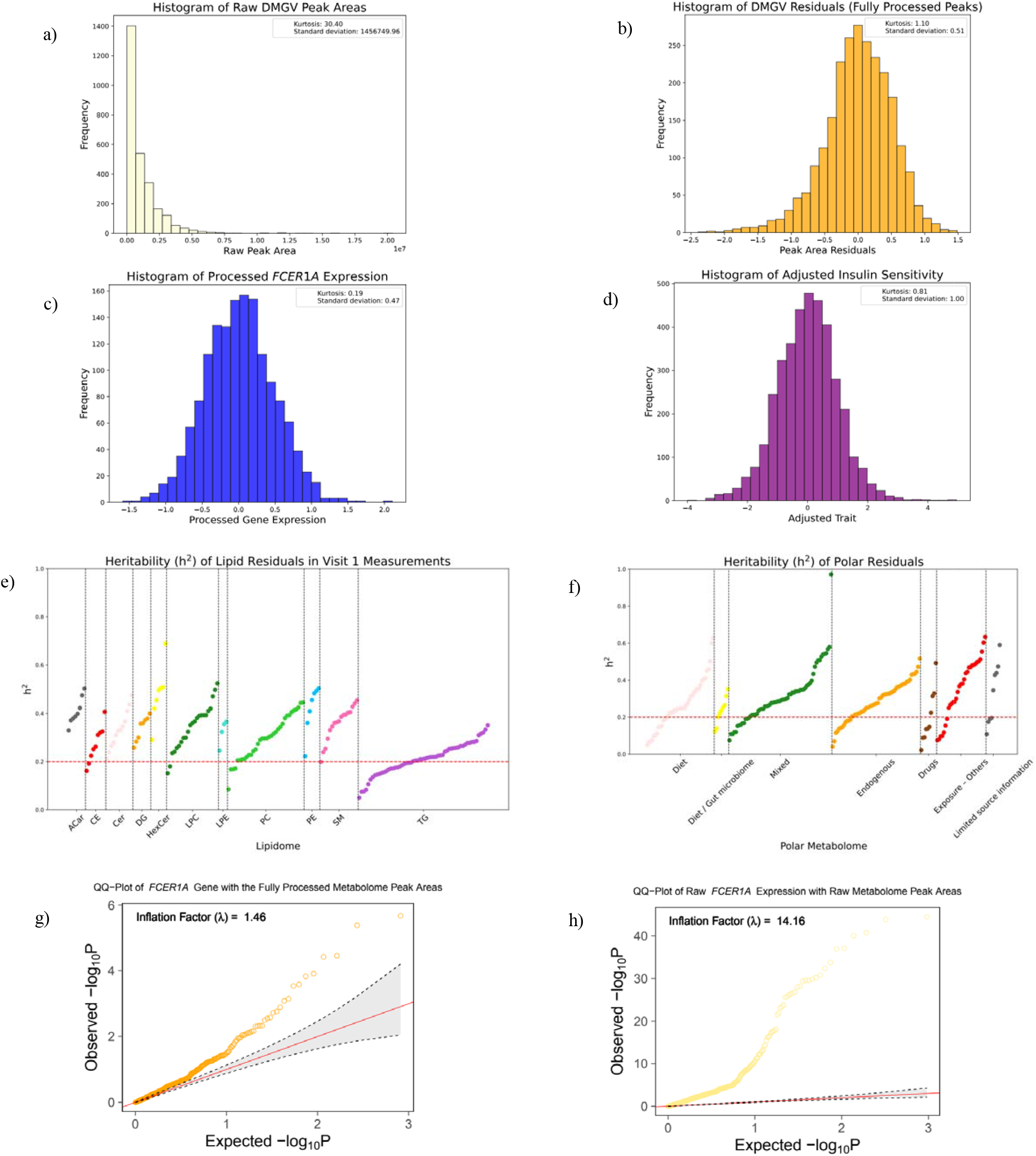
LC/MS data processing results. **a)** Distribution of raw peak areas for dimethtylguanidino valeric acid (DMGV). **b)** Distribution of DMGV upon adjustments for technical, demographical, and biological covariates. **c)** Distribution of *FCER1A* gene upon data processing. **d)** Distribution of insulin sensitivity upon covariate adjustment. **e)** Heritibality (*h*^2^) of lipids across different lipid categories. **f)** *h*^2^ of polar SMs grouped by their primary source. **g)** QQ-plot of raw *FCER1A* gene expression with raw metabolome peak areas. **h)** QQ-plot of the adjusted *FCER1A* gene expression with the adjusted metabolome peak areas. Lipid categories include: “Acar”: acetylcarnitines, “Cer”: ceramides, “CE”: cholesterol esters, “DG”: diglycerides, “HexCer”: hexosylceramides, “LPC”: lysophosphatidylcholines, “LPE”: lysophosphatidylethanolamines, “PC”: phosphatidylcholines, “PE”: phosphatidylethanolamines, “SM”: sphingomyelins. The primary source of polars include: “Diet”: from dietary sources, “Diet / Gut microbiome”: from gut microbiome metabolism, “Mixed”: from dietary and endogenous sources, “Endogenous”: synthesized internally within the body, “Drugs”, “Exposures – Others”, and “Limited source information”.

#### 2.2.2 Heritability analysis

To further validate the biological robustness of the processed LC/MS profiles, we assessed heritability (*h*^2^) using SOLAR^17^. Across both lipid and polar SM categories, strong *h*^2^ was observed (mean *h*^2^ = 0.30 for lipids and 0.294 for polars; Figures 2e & 2f), which is consistent with findings from other populations^18^. Most lipid categories showed *h*^2^, while triglycerides displayed relatively lower *h*^2^ (0.21), likely reflecting stronger environmental influence. Supporting this pattern, a strong correlation was also observed between BMI and TG levels in the LLFS (r = 0.32; Supplementary Figure 2c). Notably, heritability patterns were consistent across independent LLFS clinical visits (Supplementary Figure 3), further reinforcing the reliability of the processed data. Among polar SMs, compounds derived primarily from diet, gut microbiome, or endogenous sources exhibited strong *h*^2^, whereas drug metabolites displayed lower *h*^2^ (average *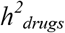* = 0.198). Interestingly, most chemical exposures showed intermediate heritability, suggesting contributions from both shared environments and genetically influenced metabolism. Together, these results confirm that the processing protocol preserves genuine biological signal while controlling confounders.

#### 2.2.3 Stable OMICS integration following data processing

Following LC/MS data processing, gene-SMs association tests were performed to evaluate p-value distribution patterns and inflation factors (λ) across the association scans. As shown in Figure 2h, gene-SMs associations based on the processed data exhibited p-value distributions closely matching uniform expectations, with most scans centering around λ = 1 (Supplementary Figure 4). This marks a substantial improvement over raw gene-SMs associations, which showed large deviations from uniformity, suggesting biased association signals (Figure 2g). For trait-omics associations, larger inflations were observed. To correct this, we applied BACON^19^, an empirical Bayesian method for controlling inflation in association scans. This adjustment resulted in p-value distributions that better adhered to the expected uniform pattern under the null hypothesis (Supplementary Figure 5).

### 2.3 GEM-Net framework enhances multi-modal module identification beyond conventional approaches

We refer to conventional network inference methods as those that construct multi-modal modules by integrating gene-gene, gene-SM, and SM-SM edges into a single edge-weight matrix, followed by unsupervised clustering to identify multi-modal modules. In this study, we applied WGCNA^20^ and GENIE3^21^ to infer edge weights across omics layers. For module identification, we used kernel-based (KB) clustering and modularity optimization (MO), the winners of the 2019 DREAM challenge for gene-level module identification^11^ (Section 5.4.1). Combinations of GENIE3 and WGCNA with MO and KB were implemented to infer multi-modal modules.

In contrast, the proposed semi-supervised GEM-Net framework first constructs gene-level modules exclusively from gene-gene edges using conventional unsupervised clustering methods. Subsequently, additional omics profiles are selectively integrated into these gene-level modules using a CMA strategy^22^. In the present study, CMA combines gene-SM associations for genes within a module while accounting for the correlation structure among the genes (Section 5.4.2). This two-step process aims to simultaneously preserve the integrity of the gene co-expression network and enable the integration of lipids or metabolites strongly correlated with genes within each module.

The evaluation strategies used to benchmark the proposed model against the conventional approaches focused on multi-modal modules and can be categorized as:

1. Enrichment analysis: Gene Ontology (GO) enrichment and metabolic set enrichment analyses (MSEA) on genes and SMs of each module.
2. Gene-level biological support: For genes within each module, biological support is assessed using PPI networks and transcription factor (TF) co-regulation. Support is defined as the proportion of genes in a module that either interact with at least one other gene based on STRING PPI^23^ or share a common regulatory TF with at least one other gene based on blood gene regulatory networks (GRNs)^24^.
3. Diversity metrics: Based on the number of inferred multi-modal modules, and the representation of genes and SMs across these modules.

#### 2.3.1 GEM-Net improves biological coherence of multi-modal modules

MSEA confirmed that the GEM-Net framework better recapitulates known metabolic relationships, with 57.47% of modules enriched for metabolic terms versus the second best-performing method, WGCNA-MO, with 39.46% enrichment (Figure 3a). We next assessed if integrating SM edges into the network for unsupervised clustering can disrupt the gene coexpression structure in the resulting multi-modal modules. The conventional methods demonstrated lower gene-level biological support compared to the GEM-Net. The GEM-Net framework exhibited the highest PPI support, with 59.33% of genes within modules on average supported by STRINGdb (Figure 3b). Similarly, TF co-regulation analysis revealed that on average, 62.85% of genes in the GEM-Net modules share a common TF with at least one other gene within the same module (Figure 3c). However, for GO enrichment analysis, WGCNA-

MO emerged as the best-performing approach, with 48.71% of modules showing enrichment for at least one GO term. GEM-Net followed as the second-best method with 24% enrichment for GO terms (Figure 3d).

**Figure 3.**
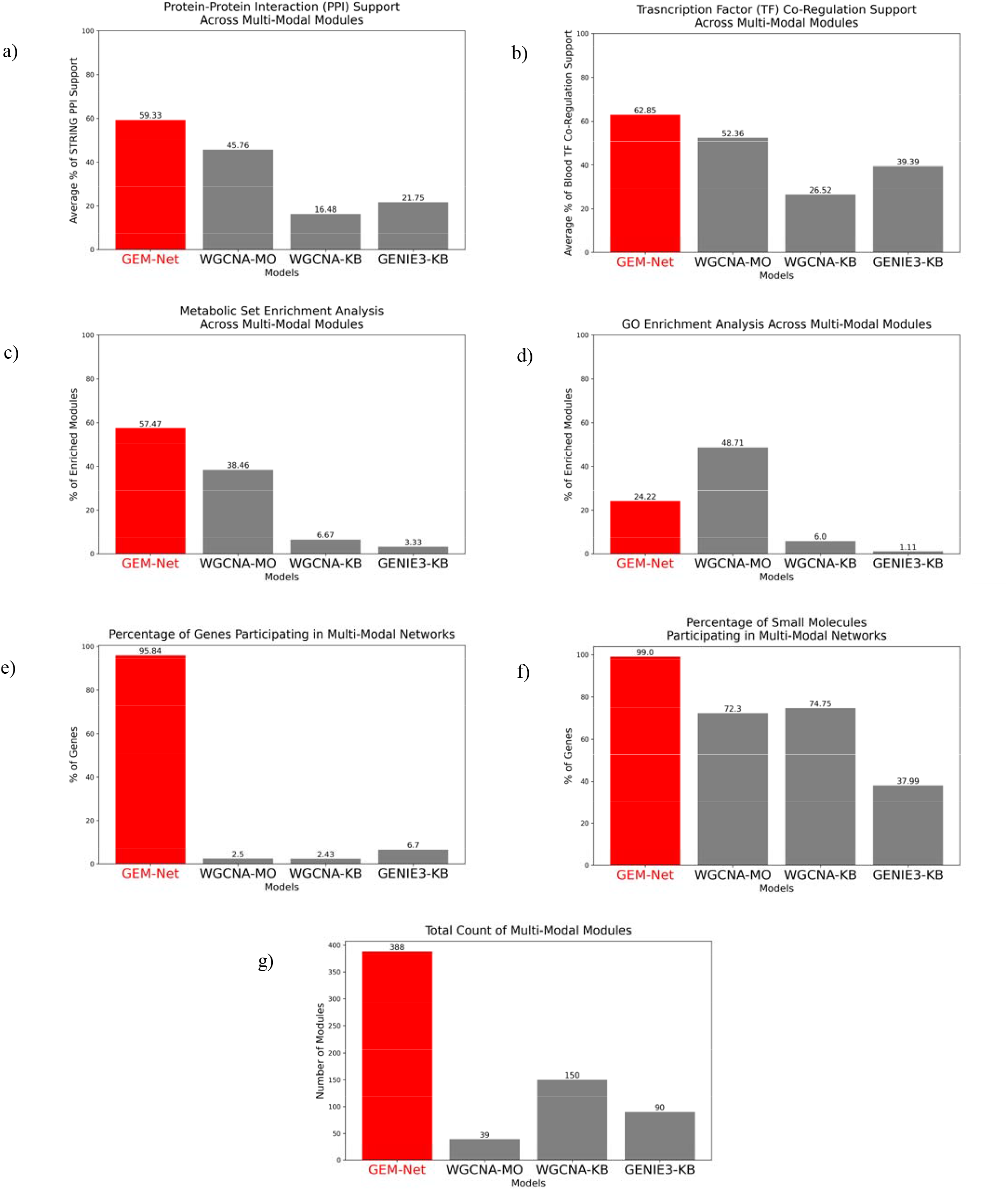
Benchmarking network inference approaches for multi-modal networks. a) Represents the proportion of multi-modal modules with significant hits from MSEA. b) Represents the average proportion of gene-gene interactions supported by STRINGdb PPI networks. For each module, proportion of support was calculated, which was averaged across all modules. c) Represents the average proportion of genes co-regulated by same TFs based on the blood-specific GRNs. d) Represents the proportion of multi-modal modules with significant GO terms. e) Representation of genes profiled in the LLFS across the multi-modal networks inferred by each method. f) Representation of SMs profiled in the LLFS across multi-modal networks inferred by each method. g) Total number of multi-modal modules inferred by each method.

#### 2.3.2 GEM-Net increases network diversity without compromising biological support

A primary goal in the present study is to increase the information content of omics data by incorporating gene, SMs, and their interactions. Thus, we assessed how well different approaches preserved the diversity of genes and SMs within the multi-modal networks. The GEM-Net framework retained 99% of SMs and 96% of genes in LLFS modules. GEM-Net represents a nearly 38-fold increase in gene retention compared to the second-best method from the independent biological support, WGCNA-MO, which retained only 2.5% of genes (Figure 3e). Similarly, SM representation based on the conventional models ranged from 38% (GENIE3-KB) to 74% (WGCNA-KB) (Figure 3f). Lastly, The GEM-Net framework yielded a total of 388 multi-modal modules across the entire network—an almost 10-fold increase compared to WGCNA-MO (Figure 3g). Importantly, GEM-Net simultaneously improved independent biological support for genes and SMs (Section 2.3.1) and preserved the integrity of gene coexpression patterns while markedly increasing network diversity.

#### 2.3.3. GEM-Net properties

Having benchmarked the performance of the GEM-Net framework against conventional approaches, we next extended the framework to construct knowledge-guided GEM-Nets. These networks were built by integrating SMs into pre-existing gene-level modules derived from STRING PPI, InWeb PPI, and GEO coexpression networks^11^.

In both LLFS-derived and knowledge-guided GEM-Nets, SMs consistently integrated into gene-level modules, constituting ∼21% and 16% of module features, respectively (Figure 4a). In contrast, modules inferred by the conventional methods exhibit a higher proportion of SMs (Supplementary Figure 6). This is due to the reduced network diversity (Figure 3e & 3g), which limits their capacity to distribute SMs across diverse modules. Furthermore, even though the framework incorporates SMs into the gene-level modules based on gene-SM associations, SMs exhibited strong intra-module. Module-wise PCA revealed that PC1 on average explains 44% of the variance among SMs in modules (Figure 4b). Knowledge-guided GEM-Nets exhibited a PCA pattern consistent with that of the LLFS GEM-Nets (Supplementary Figure 8). Lastly, although small molecules are integrated based on gene–SM associations, SM–SM association tests revealed that 91% of modules contain small molecules with significant within-module associations (average p-value < 0.05), which further highlights a coherent multi-omics structure. (Figure 4c).

**Figure 4.**
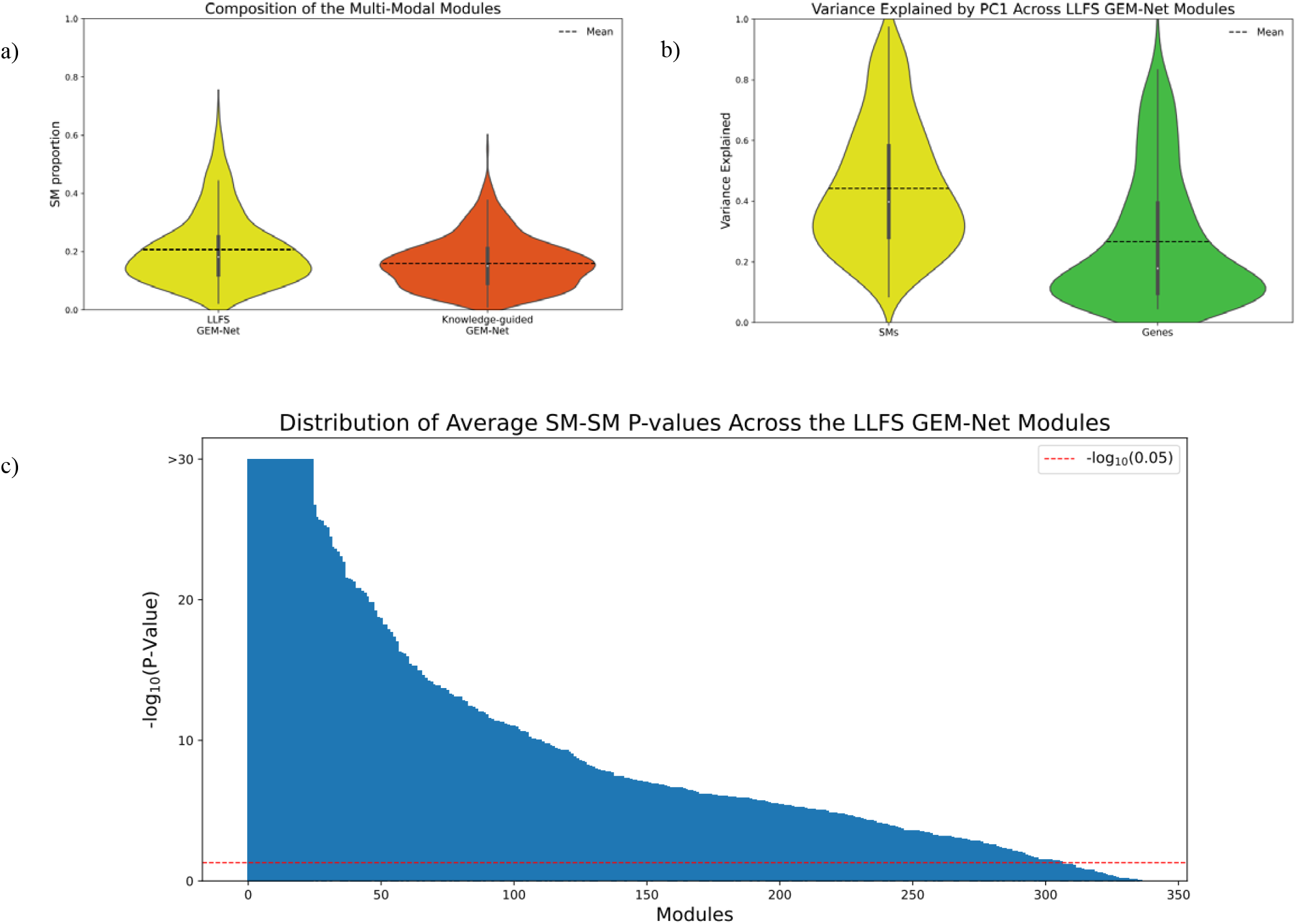
GEM-Net properties. **a)** Violin plots representing the composition of the GEM-Net modules based on the proportion of SMs across modules in the LLFS and the knowledge-guided GEM-Nets. **b)** Violin plots of the variance explained by PC1 across the LLFS GEM-Net modules. For each module, separate PCA were performed on genes and SMs participating in the module. **c)** Histogram of the average –log_10_(p-value) of SM-SM associations in the GEM-Net modules. In each module, pairwise SM-SM associations were assessed and subsequently averaged for the number of pairs. The maximum range of the plot was set to –log_10_(p-value) ≤ 30 for readability.

### 2.4 GEM-Net Modules Link Microbial Metabolites to Immune Regulation in Aging Metabolism

Traditional omic-wide association studies (OWAS), such as transcriptome-wide association studies (TWAS) or MWAS, are limited to identifying individual marker–trait associations without capturing broader regulatory context. In contrast, GEM-Net modules can serve as interpretable units that integrates multiple omics modalities to investigate systems-level molecular patterns underlying complex traits.

Module association tests were conducted using Pascal^25^ to identify GEM-Net modules significantly associated with IS. This analysis revealed seven significant knowledge-guided modules (p < 8.88E-5; Supplementary Table 1) and twenty significant LLFS modules (p < 1.29E-4; Supplementary Table 2). Most GEM-Net modules included one or more nodes that were also significant in OWAS analyses (TWAS or MWAS—Supplementary datasheets). However, they also contained a larger number of genes and SMs with suggestive associations (p < 0.05) that are overlooked in single-omic analyses.Module-based association testing can recover these signals when they participate in the same module, allowing individually weak associations to aggregate into a stronger, detectable signal.

The top-ranked module (p = 4.10E-10) in the knowledge-guided GEM-Net reveals a protective cross-talk between the gut microbiome metabolism and blood immune system for metabolic health. This module contains four transcriptome-wide significant genes (Supplementary Table 3)—*FCER1A, HDC, MS4A2*, and *CPA3*—and two metabolome-wide significant SMs: N-acetylglycine (NAG) and dimethylguanidino valeric acid (DMGV). All genes are positively connected to NAG and negatively connected to DMGV, revealing opposing SM–gene interaction patterns (Figure 5a).

The genes in this module are well-known in mast cell–mediated inflammatory responses^26, 27^. However, in the LLFS cohort, they show protective associations with IS and are also associated with lower BMI, TG, and IL6 (Figure 5b). These asociations were also replicated in the FHS cohort with consistent effect directions (Supplementary Table 3). Prior population-based studies have identified the associations of *HDC, MS4A2*, and *CPA3* with lower glucose levels in non-diabetic individuals^28^.Additionally, *FCER1A* has been linked to multiple anti-inflammatory functions across white blood cell lineages^29, 30^.

NAG, a microbiome-derived metabolite linked to fiber metabolism^31, 32^, is associated with higher IS and lower BMI, and TG in the LLFS. These associations are supported by prior population studies linking NAG to improved metabolic health^33-35^. In addition, NAG supplementation improves weight loss in diet-induced obese mice^36^. In contrast, DMGV—an oxidative stress–related metabolite^37^— exhibits associations with higher BMI, TG, and IL6 levels, and lower IS. These findings align with prior reports linking DMGV to increased risk of metabolic and cardiovascular complications^38, 39^.Notably, DMGV and NAG exhibit a strong inverse association (p = 4.65E-34) in the LLFS, reinforcing their antagonistic effects.

The structure of the GEM-Net module reflects these functional patterns: NAG is positively connected to *FCER1A, HDC, MS4A2*, and *CPA3*, aligning with the protective associations of NAG and the genes for the metabolic traits. In contrast, DMGV is negatively connected to these genes and is associated with adverse metabolic and inflammatory outcomes (Figure 5). These patterns are consistent with the established role of fiber intake and fiber-degrading microbiome activity in reducing inflammation across various conditions^40^. Supporting this, *rs2251746*, an intronic variant in FCER1A, has also been associated with Ruminococcaceae abundance^41^, a key fiber-degrading bacterial family in gut^42^.

Combining the results together, the analysis of GEM-Net reveals the novel cross-talk between the gut microbiome metabolism and the immune system, driven by the interaction of NAG with *FCER1A*, HDC, MS4A2, and CPA3. This axis is strongly associated with improved IS and reduced obesity and markers of inflammation, including DMGV and IL6. While prior studies have individually linked some of these genes and SMs to metabolic health, their joint involvement in a shared regulatory module and their opposing roles within this module was not previously recognized.

## 3. Discussion

This study presents a generalizable framework for constructing and interpreting multi-modal networks from the high-throughput omics data. We demonstrate the utility of this framework using transcriptomics, metabolomics, and lipidomics profiles from the LLFS to build multi-modal networks and identify novel regulatory patterns associated with metabolic health (Figure 1).

The foundational step for multi-modal network inference is adjusting omics profiles for confounding effects and harmonizing them across data modalities. RNA-seq and LC/MS differ in data structure and noise, and each requires its own pre-processing strategy. We demonstrate that applying a similar mathematical workflow comprising data transformation, dimensionality reduction, and covariate adjustment to each omic profile independently produces harmonized distributions while controlling for technical and biological confounders. This process yields residuals that approximate a normal distribution (Figure 2a, Supplementary Figure 1), with reduced technical variability (Figure 2b–c) and preservation of biological signals. Rather than recommending specific algorithms, we emphasize selecting methods suited to study design and data characteristics. Diagnostic checks, such as distribution plots, heritability estimates, and QQ-plots can guide these choices. In the LLFS, for instance, we used a random forest–based method for LC/MS batch effect correction, though simpler models may suffice in smaller cohorts. This protocol makes the foundation for GEM-Net construction. But beyond network inference, it is broadly applicable to population-scale multi-omics studies.

GEM-Nets are constructed using a semi-supervised strategy that builds on gene-level modules and selectively integrates other omics profiles using a meta-analysis strategy. In contrast, unsupervised module identification methods use all pairwise interactions with limited structural constraints. Our approach preserves transcriptomic structure, handles missing data across omics modalities, and avoids overfitting to sparse or imbalanced multi-omic data, leading to more diverse and biologically supported modules (Figure 3). To evaluate GEM-Nets, we applied a multi-layered validation framework that extends beyond solely pathway enrichment analyses. While enrichment analyses remain important, they are limited by incomplete annotations of many SMs and genes^43^. Our evaluation incorporates additional high-throughput evidences, such as PPIs and TF co-regulation, to assess module integrity (Figure 3). Notably, although SMs are integrated based on gene-SM associations, we observed strong pairwise associations among SMs within modules (Figure 4e), suggesting that biologically meaningful multi-omic structure can emerge without explicit modeling of SM–SM links.

Analysis and biological interpretation of GEM-Nets highlight their advantage over single-omic association studies. Traditional OWAS identify trait-associated markers, but they do not reveal broader regulatory context or interactions between omics layers^44^. GEM-Net enables the detection of trait-relevant modules containing both significant and suggestive omics features, grouped by shared statistical patterns. The top-ranked module for IS illustrates this: it links NAG, a microbiome-derived metabolite^31, 32^, with four immune-related genes—*FCER1A, HDC, MS4A2*, and *CPA3*—all showing protective associations with metabolic traits in the LLFS and replicated in FHS (Figure 5). In contrast, DMGV, a pro-inflammatory metabolite^38^, was negatively connected to the same genes and associated with adverse metabolic outcomes. While these genes and SMs have been previously identified for some metabolic phenotypes in isolation^28, 33^-^35, 38, 39^, their joint regulatory behavior was only uncovered through network-based analysis. Moreover, both NAG and DMGV remain functionally understudied, highlighting the value of resources like GEM-Net that can place uncharacterized features within biological context.

**Figure 5.**
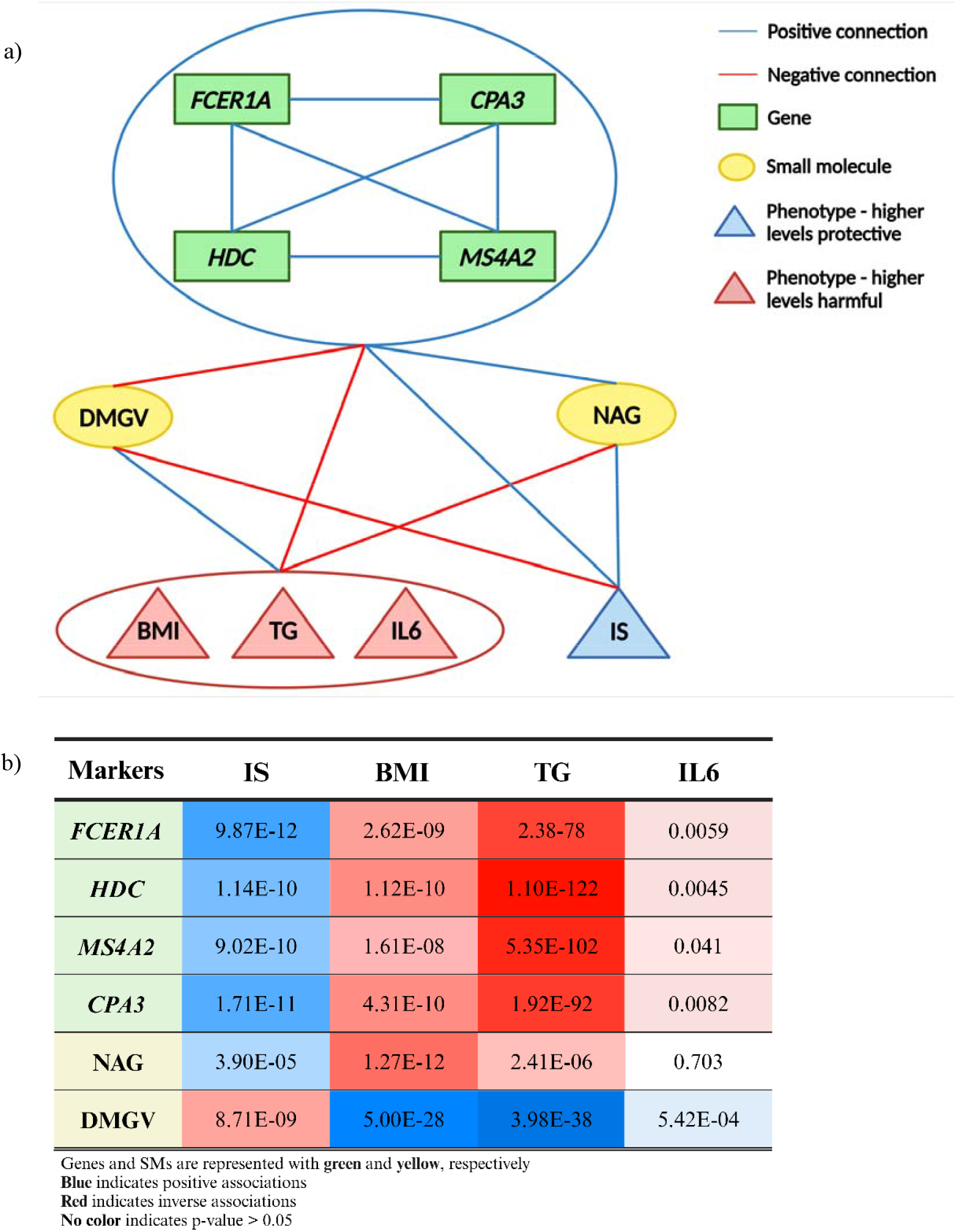
Demonstration of the significant gene-SM interactions in the top-ranked GEM-Net module for IS. **a)** NAG and DMGV are connected to all 4 genes illustrated in the sub-module. NAG is also connected to BMI and TG (not IL6), DMGV is connected to BMI, TG, and IL6, and the 4 genes are connected to BMI and TG. However, only *FCER1A, HDC*, and *CPA3* are connected to IL6. The large blue and red ellipses are used for clear illustration. **b)** Association summary of the significant nodes of the top-ranked module for the metabolic traits.

The GEM-Net framework offers a flexible paradigm for multi-omics discovery. While it captures biologically meaningful cross-omics interactions, it is based on statistical associations and does not imply causality. Functional validation will be essential to confirm the mechanistic roles of the identified interactions. In this study, we demonstrated the framework’s applicability through the integration of transcriptomic, metabolomic, and lipidomic data. However, GEM-Net is readily extensible to other gene-based omics layers, such as proteomics, and can incorporate additional profiles including DNA methylation and genetic variation profiles. Furthermore, GEM-Net modules provide a resource for downstream machine learning applications, such as graph neural networks, which may enhance module detection or improve trait prediction.

## 4. Conclusions

This study presents a framework for multi-omics integration, network inference, and biological interpretation, addressing key challenges in LC/MS data processing, multi-modal module identification, and network-guided trait analysis. We demonstrate the ability of GEM-Nets to uncover biologically meaningful multi-modal modules and cross-omic interactions. The integration of SMs from diverse sources further highlights the framework’s potential to advance SM pathway discovery, particularly for uncharacterized or understudied compounds. Importantly, the inferred GEM-Nets offer a resource for multi-omics investigations in human populations, providing reference networks derived from the healthy individuals from the LLFS. More broadly, GEM-Net framework establishes a scalable foundation for network-based multi-omics discovery across complex traits and disease contexts.

## 5. Materials & Methods

### 5.1 Participants

Procedures and criteria for eligibility and recruitment of the LLFS participants are described in detail by *Wojczynski et al^4^*. For this study, data from the first clinical exam containing 4953 total participants from 539 families was used. Glucose (mg/dL), insulin (pmol/L), TG (mg/dL), IL6 (pg/mL), hemoglobin (g/dL), and glycosylated hemoglobin (HbA1c) were measured by the Advanced Research and Diagnostics Laboratory at the University of Minnesota. Participants with clinical diagnosis of type 2 diabetes mellitus (T2DM) or those with fasting glucose levels ≥ 126 mg/dL or HbA1c ≥ 6.5% were excluded. Additionally, all non-diabetic participants with fasting time < 8h were also excluded to avoid any metabolic bias. After applying these criteria, the final study sample consisted of 3,839 individuals. IS was calculated by HOMA2 software using fasting insulin and glucose measurements^45^. BMI was calculated as weight (kg) / height (m^2^). All traits were adjusted using a stepwise regression model, where age, age^2^, age^3^, sex, and clinical field centers served as base covariates, and the top 20 genetic PCs were included as additional covariates. IS, TG, and IL6 were ln-transformed prior to covariate adjustments.

### 5.2 LC/MS workflow

In the LC/MS workflow, each batch generally consisted of 92 research samples, 2 quality control (QC) samples, and 2 blank samples. QC samples were prepared by pooling a subset of the research samples. Peak lists were generated through MS feature detection, background subtraction, and adduct selection. Following peak list generation, compounds were identified by annotating mass, MS/MS fragmentation patterns, and retention times to both in-house and online libraries. Peak areas were then obtained for the identified compounds. To account for batch effects in raw peak areas, a random forest method was applied^15^, leveraging QC variability within each batch. These peak areas, adjusted for batch effects, were further processed to account for potential biological confounders. The variables include chronological age, age^2^, sex, smoking status, and medication usage. A stepwise regression model coupled with PCA were used for covariate adjustment. All steps in the workflow were conducted separately for polar SMs and lipids. Detailed protocol information is provided in the Supplementary Text S1.

### 5.3 RNA-seq protocol

RNA extraction and sequencing were performed by the McDonnell Genome Institute (MGI) at Washington University. Total RNA was obtained from PAXgene™ Blood RNA tubes through the Qiagen PreAnalytiX PAXgene Blood miRNA Kit (Qiagen, Valencia, CA). RNA-Seq data processing was carried out by version 3.3 of the nf-core/RNASeq pipeline with STAR/RSEM, applying default parameters (https://zenodo.org/records/5146005). Genes with fewer than three counts per million in over 98.5% of samples were excluded, and samples with more than 8% intergenic reads were also removed. The remaining data were transformed by the variance stabilizing transformation (VST) function in DESeq2^46^. The transformed gene expression levels were then adjusted through a stepwise regression model for age, age^2^, sex, field centers, percent of intergenic reads, and the counts of red blood cells, white blood cells, platelets, monocytes, and neutrophils as baseline covariates. Furthermore, RNA-seq batch information and the top 10 PCs of gene expression were incorporated into the model as additional covariates^47^.

### 5.4 Multi-modal network inference

#### 5.4.1 Conventional methods

Traditionally, network inference in single-omic profiles (e.g., RNA-seq) relies on unsupervised learning, where feature relationships (commonly refer to as edge weights) are first quantified using similarity metrics. An unsupervised clustering algorithm is then applied to the resulting edge weight matrix to define modules of densely connected features. We refer to this as the ^‘^conventional^’^ network inference approach.

For the conventional methods used in this study, adjusted SM profiles (comprising polar metabolites, lipids, and identified exposures) were combined with adjusted gene expression profiles from the start of the network inference workflow. Edge weights of gene-gene, gene-SM, and SM-SM pairs were computed using two established methods: WGCNA^20^ and GENIE3^21^. For WGCNA, the optimal soft-threshold (β = 2.3) was selected based on an approximate scale-free topology index of 0.9 and mean connectivity of 15. The resulting adjacency matrix was converted into a topological overlap matrix. For GENIE3, a forest-based model was applied with default parameters.

Following edge weight computation, clustering was performed using MONET software, which implements two top-performing methods from the 2019 DREAM Challenge for gene-level module detection: MO and KB clustering^11, 48^. The conventional network inference workflow thus consists of WGCNA and GENIE3 in combination with KB and MO clustering approaches. A detailed description of these methods is provided in Supplementary Text S2.

#### 5.4.2 Gene-Embedded Multi-Modal Network (GEM-Nets)

The proposed model combines conventional unsupervised network inference with a supervised strategy to selectively integrate SM and RNA-seq profiles, creating multi-modal networks. First, gene-level modules were constructed from the adjusted RNA-seq data of the LLFS using conventional approaches, as described earlier. These modules served as input entities for the supervised integration of SM profiles, guided by statistical associations and a meta-analysis approach. A two-step significance assessment was developed to connect SMs to the genes in the input modules.

Initially, associations between each SM and individual genes across co-expression modules were calculated using linear mixed models (LMMs). After multiple testing correction for SM associations with each gene in the co-expression network, SMs with significant associations were directly connected to their corresponding genes. In the second step, a CMA^22^ strategy was applied to refine SM integration based on the gene-SM associations in each module. For a SM, CMA combines gene-SM associations of genes within each module, while accounting for the inter-correlation of these genes using a covariance matrix derived from gene-SM associations. To perform CMA, a predefined p-value threshold of 0.0025 for gene-SM associations was used to only select nominally significant SMs for genes of a module. This threshold was chosen as it is approximately an order of magnitude larger than the Bonferroni significance threshold of lipidome or polar metabolome in the LLFS. SMs with significant meta-analytic p-value after multiple testing correction were then connected to their respective genes in each module. The detailed statistical and algorithmic framework of the semi-supervised approach is described in the Supplementary Text S3.

#### 5.4.3 Knowledge-Guided GEM-Nets

In addition to the semi-supervised multi-modal network inference based on data-driven co-expression networks from the LLFS, knowledge-guided GEM-Nets were constructed using external sources. Specifically, PPI networks from STRINGdb and InWeb, as well as coexpression networks derived from GEO were used as the baseline gene-level networks. Modules of these networks were selected from those generated by the winners of the 2019 DREAM challenge for unsupervised network clustering^11^.SMs were then integrated into these baseline networks using the same two-step semi-supervised approach described earlier.

### 5.5 Multi-Modal Network Inference Evaluation

Evaluation was performed at the module level based on functional enrichment, external gene level support and diversity metrics to assess the biological coherence and diversity of the multi-modal modules produced by each method. For functional enrichment, MSEA was performed using MetaboAnalyst 6.0^9^. Additionally, GO enrichment analysis was conducted using GOATOOLS^49^. Gene-gene interaction support was evaluated using STRINGdb PPI networks^23^, where support was defined as the proportion of genes within a module that interact with at least one other gene in the same module. Similarly, TF co-regulation support of genes within each module was assessed using human tissue-specific GRNs for blood cell lineages^24^. Support was defined as the proportion of genes sharing a common TF with at least one other gene in the module. Lastly, network diversity was evaluated by examining the number of modules, genes and SMs participating in the networks.

### 5.6 “Omics-wide” and network association tests

LMMs were used for all genome-wide, transcriptome-wide, metabolome-wide, and lipidome-wide association tests to account for familial relatedness based on the LLFS kinship matrix (details in Supplementary Text S3). To control for inflation factors for each “omics”-wide association study, the BACON method was applied if the inflation factor (λ) ≥1.2 ^19^. Lastly, to test the association of multi-modal modules with complex traits, Pascal was used^25^. This method computes the sum of chi-squared statistics upon ranking all omics units across the entire network based on their significance for a complex trait.

### 5.7 Framingham Heart Study

Replication of the transcriptomic analysis for each trait was conducted using data from the FHS cohort. FHS is a multi-generational, family-based study investigating genetic, molecular, and environmental factors influencing cardiovascular and related traits^50^. For this study, we utilized data from the second examination of the third-generation cohort, which includes the largest number of RNA-seq measurements available. Eligible participants were selected based on the same criteria as those used for the LLFS cohort, and all traits were adjusted in a manner consistent with the methods outlined in Section 5.1. RNA-seq measurements were processed and adjusted following the same method described in Section 5.3. After applying these criteria and adjustments, a maximum of 1,248 subjects were included in the transcriptomic analysis.

## Data and code availability

The LLFS data is available at the Exceptional Longevity Translational Resources portal (https://prod.eliteportal.synapse.org/Explore/Projects/DetailsPage?shortName=LLFS). LLFS and knowledge-guided GEM-Net modules and their respective gene-SM interactions are available at

https://doi.org/10.5281/zenodo.15003731. TWAS, MWAS, and network association summary statistics for IS are provided as supplemental information. All code implementations are available at https://github.com/vaha-am/GEM-Net.

## Acknowledgements

We extend our gratitude to the Long Life Family Study consoritum, including the participants, investigators, and administrative and clinical staff

## Author contributions

V.A.M conceptualized the project, developed the methodologies, processed the LLFS data, performed all formal analyses, wrote the manuscript, and revised the manuscript. S.A contributed to project conceptualization, RNA-seq data processing, FHS data processing, and writing the RNA-seq data processing methods. M.S.H contributed to the development of LC/MS protocol and supervised the LC/MS data processing. S.L processed the FHS data, and contributed to writing the FHS methods section. W.J.J assisted with the development of CMA algorithm for network inference. B.T supervised all of the LLFS blood assays. L.P.S supervised the development of LC/MS protocol. E.W.D and N.L.S contributed to the development of statistical methodologies. P.A contributed to phenotype adjustments and selection criteria of the participants. M.R.B supervised the RNA-seq protocol and contributed to project conceptualization. G.J.P supervised the LC/MS protocol and contributed to project conceptualization. M.A.P conceptualized the project and supervised the statistical methodologies. All authors contributed to the revision of the manuscript.

