## Supplementary Text for "Gene-Embedded Multi-Modal Networks for Population-Scale Multi-Omics Discovery"

**S1. Metabolomics protocol**

Upon sample extraction from blood, pooled samples were first created from a subset of samples to serve as quality controls (QCs) in batches. Acetonitrile/methanol and methyl tert-butyl ether/ methanol were used to extract polar and lipid metabolites respectively. For both polar metabolomic and lipidomic analyses, samples were randomized and batches were created to generally contain 92 research samples, 2 QCs, and 2 blank samples. For polar metabolites, XCMS^1^ and CAMERA^2^ were used for peak detection, background subtraction, and adduct selection to generate polar peak lists. Lipid Annotator was used to generate peak lists for lipidomic analysis. MS features with intensities less than three times higher in study samples compared to blank samples were excluded. Upon acquiring LC/MS/MS data, compound identification was done by annotating mass and MS/MS fragmentation patterns to in-house and online libraries. The process were further refined by matching with retention times. However for lipids, sum composition is provided for lipid annotations. Peak areas were then obtained by Skyline^3^ for each batch. These peak areas were further adjusted to account for technical and biological confounders.

To correct for batch effects, a random forest-based approach was applied to the raw peak areas^4^. After log2 transformation of peak areas, batch number and QC sample run order were used to model each small molecule’s (SM) intensity deviation in each QC sample from the mean intensity of that molecule across all QC samples, with one model per small molecule. The model was trained on QC samples, and applied on research samples to predict the deviation of SM peak areas in research samples. The predicted deviation was subtracted from the peak area for each SM, yielding a semi-processed data.

Next, this semi-processed data were adjusted for potential biological confounders. First, principal component analysis (PCA) was conducted separately for polar and lipid SMs to obtain the top 10 principal components (PCs) within each category for further adjustments. A stepwise regression model was applied to the semi-processed peaks of each SM, where chronological age, age², sex, and field center information were baseline covariates. Additional covariates included smoking status (binary), medication usage across five categories (hypertension drugs, general lipid-lowering drugs, statins, angina drugs, and diabetes drugs), and the top 10 PCs. The fully processed data are represented by the residuals of the stepwise regression model, calculated as the difference between the semi-normalized peak values and their predicted values. The fully processed were used for all downstream analyses. Supplementary Figure 1 depicts the raw and fully processed peak area distributions for each SM category across all samples.

**S2: Conventional network inference approaches: Technical details**

***Edge weight calculation***

Two methods that were used in this study to calculate edge weights were based on weighted correlation analysis – WGCNA^5^ – and random forest-based model, GENIE3^6^.

**WGCNA.** For WGCNA, correlation matrix was first created by calculating the Pearson’s correlation coefficient for all possible pairs of genes, gene-SMs, and SMs. The correlation matrix was then transformed to an adjacency matrix by applying a soft-threshold as follows:

$a_{ij}=r_{ij}^{\beta}$	(1)

Where *a_ij_* and *r_ij_* are the adjacency and correlation between omics features *i* and *j* respectively, and *β* is the soft-threshold. The choice of *β* was based on the scale-free topology index as well as mean connectivity. The scale-free topology index indicates how closely a graph follows a scale-free topology based on power-law distribution. On the other hand, connectivity, *k*, is defined by the sum of all adjacency values for an omics feature except for itself:

$k_{i}=\sum_{j\neq i} a_{ij}$	(2)

To determine the best *β* value, adjacency matrices were created from *β =* 1 to *β* = 20 and the scale-free topology index and mean connectivity were calculated for each matrix. The selection of *β* was based on the scale-free topology index being closer to 0.9 with connectivity remaining between 10-20, resulting a graph with certain hub nodes that have stronger connection to other nodes while most nodes are weakly connected to each other, better reflecting the biological properties*.* The best adjacency matrix was then transformed to a topological overlap matrix (TOM). The topological overlap similarities for omics features were used as the inputs for the clustering algorithms.

**GENIE3**. The python version of GENIE3 was used to calculate the edge weights among the omics features (<https://github.com/vahuynh/GENIE3/tree/master/GENIE3_python>) based on the random forest model. All default variables were used in the model. One model was trained per each omics feature and the importance of all other omics features for the target feature were calculated to construct a feature importance matrix, which was subsequently inputted to the clustering algorithms.

***Clustering***

To perform clustering on the topological overlap matrix from WGCNA and the feature importance matrix of GENIE3, The MONET package^7^ were used. MONET includes the top-performing algorithms of the 2019 DREAM challenge for unsupervised clustering of gene-level networks (e.g. co-expression networks, protein-protein interaction networks, etc…)^8^. The unsupervised clustering methods include a kernel-based (KB) clustering algorithm based on the diffusion state distance^9^, a modularity optimization (MO) algorithm based on multiple resolution screening of cluster structures within a network^10^, and a random walk algorithm. For each clustering algorithm, the fully connected network based on TOM and feature importance matrix, as well as subsets of the complete network consisting of the top 1000, 500, 250, and 100 genes for each SM were used. Input parameters of each algorithm were also iteratively adjusted. The selection of best resulting clusters was based on the number of multi-modal clusters as well as the average number of SMs across all multi-modal clusters. It is worth noting that the random walk algorithm did not yield any multi-modal cluster and were excluded for downstream analyses.

**S3: Gene-Embedded Multi-Modal Network (GEM-Net): Statistical framework**

The proposed model selectively integrates omics profiles into a baseline gene-level network. This process requires two inputs: (1) a baseline network and (2) cross-omics association data. In this study, the baseline networks are derived from the adjusted RNA-seq data, while metabolome and lipidome profiles (collectively referred to as SM profiles) are integrated into the baseline networks in a supervised manner.

Gene-level networks and their modules were constructed using conventional approaches (detailed in S2) applied to the adjusted RNA-seq measurements. The selective addition of SMs is performed in two steps, involving statistical associations and meta-analysis.

***Step 1: Individual Associations***

Given the family-based structure of the Long Life Family Study (LLFS), gene-SM associations were calculated using linear mixed models (LMMs):

$y=X\beta+Zu+\epsilon$	(3)

where *y* represents adjusted gene expression, *X* is the design matrix for fixed effects (adjusted SM peak intensities), *β* is the vector of fixed effect coefficients, *Z* is the design matrix for random effects based on familial relatedness, and *u* is the vector of random effects following a multivariate normal distribution:

$u\sim N\left( 0,\mu_{u}^{2}K \right)$	(4)

where *K* is the kinship matrix derived from the familial structure of the LLFS dataset.

Significant SM associations for each gene were determined using the Bonferroni multiple testing threshold based on the number of SMs tested. These significant SMs were initially connected to their corresponding genes within each module. Note that the choice of the initial test statistics is dependent on the design and structure of the population-level study. For instance, for independent and identically distributed variables, a simple linear regression model can be used.

***Step 2: Meta-Analysis Using CMA***

In the second step, gene-SM connections were further refined leveraging the co-expression structure of modules based on a meta-analysis strategy. The purpose of the meta-analysis strategy is to combine the SM-association information of genes to account for overlooked gene-SM connections due to the multiple testing burden. We use a modified form of the traditional Stouffer’s meta-analysis strategy called “Correlated meta-analysis” (CMA) to achieve this.

The Stouffer’s meta-analysis is based on the sum of z-scores for common “predictors” across multiple “outcomes”. Assuming a standard normal distribution, probit transformation of p-values can be applied to obtain the z-scores:

$z=\Phi^{-1}\left( 1-p \right)$	(5)

where Φ^-1^, the probit function, is the inverse cumulative distribution function of normal distribution. Following transformation of p-values to z-scores, the Stouffer’s method calculates Z_meta_ as:

$Z_{meta}=\frac{\sum_{i=1}^{k} z_{i}}{\sqrt{k}}$	(6)

where *k* is the number of outcomes. However, Stouffer’s method assumes independence among outcomes, which is violated in the context of this study as genes within a co-expression module are highly correlated. To address the issue of inter-correlation among the outcomes, which are the genes in a co-expression module in our case, we used the CMA framework introduced by *Province & Borecki^11^*.

Suppose we have association scans of M common predictors (e.g. SMs) for *k* outcomes (e.g. expression of N genes). Upon probit transformation of p-values for each outcome (*p_kx1_*) to their corresponding z-scores (*z_kx1_*), *Province & Borecki* proved that if *z_Nx1_* is a multi-variate normal random variable, then:

$z_{kx1}\sim N\left[ \mu_{kx1},\sum_{kxk} \right]$	(7)

Where *∑_kxk_* is the variance-covariance matrix of *k* outcomes. *∑_kxk_* can be derived from the tetrachoric correlation of the z-scores of the *M* common predictors among *k* outcomes. Subsequently, CMA incorporates the inter-correlation of the outcomes in meta-analysis through the estimated *∑_kxk_* . Therefore, *Z_meta_* can be represented as:

$Z_{meta}=\frac{\sum_{i=1}^{k} z_{i}}{\sqrt{\text{SUM}\left( \sum_{kxk} \right)}}$	(8)

For example, if all genes in a module are perfectly correlated (*z_1_ = z_2_ =...= z_k_*), all elements of the variance-covariance matrix will be equal to 1 and therefore, *SUM(∑_kxk_)* = *k^2^*. Therefore, *Z_meta_* will be:

$$Z_{meta}=\frac{\sum_{i=1}^{k} z_{i}}{\sqrt{\text{SUM}\left( \sum_{kxk} \right)}}=\frac{kz}{\sqrt{k^{2}}}=z$$

In other words, in this hypothetical scenario the meta-analytic z-score will be equal to the input z due to the perfect correlation of outcomes

***Setting up CMA for GEM-Nets***

Although CMA effectively accounts for the inter-correlation of genes within a co-expression module, combining gene-SM associations for all genes in a module can result in false-positive connections. This can occur when strong associations between certain genes and SMs inflate the meta-analytic outcome, even if other genes in the module exhibit weak associations. To mitigate this issue, we implement a pre-defined p-value threshold (p < 0.0025) to select sub-clusters of genes within a module as inputs for CMA. This threshold is approximately one order of magnitude less stringent than the Bonferroni significance threshold applied to polar metabolome or lipidome profiles in the LLFS.

Let *G = {g_1_*, *g_2_,..., g_k_}* be the set of *k* genes in a module and *S={s_1_, s_2_,..., s_m_}* be the set of all common SMs tested for *k* genes. Then, for each gene in the module, *g_i_*, SM p-values can be represented as:

$p_{g_{i}}=\text{\{}p_{g_{i}s_{1}},p_{g_{i}s_{2}},...,p_{g_{i}s_{m}}\text{\}}$	(9)

Based on the pre-defined threshold, the subset of genes, $G_{s_{j}}$, for a given SM (*s_j_*) can be defined as:

$\forall s_{j}\in S, G_{s_{j}}=\text{\{}g_{i}\in G:p_{g_{i},s_{j}}<0.0025\text{\}}$	(10)

If $\text{|}G_{s_{j}}\text{|} \geq2$, the genes in $G_{s_{j}}$form the input for a CMA process.

This way, all eligible unique combination of genes within a module based on the pre-defined threshold can be obtained and a CMA is conducted on each unique set to obtain the meta-analytic p-value of the SMs meeting the criteria of the p-value threshold. All SM associations for the genes in$G_{s_{j}}$are used to construct the variance-covariance matrix, maximizing the available information. However, CMA is restricted to SMs that meet the *p* < 0.0025 threshold for the genes in$G_{s_{j}}$, reducing multiple testing burden and computation time.
