## Supplementary Figures and Tables for "Gene-Embedded Multi-Modal Networks for Population-Scale Multi-Omics Discovery"

**Supplementary Table 1. Network association summary of the significant knowledge-guided TSI-Net modules**

| **Module ID** | **Module P-value** | **Available Nodes** | **Significant Nodes*** | **Suggestive Nodes**** |
| --- | --- | --- | --- | --- |
| **GEO97** | 4.10E-10 | 63 | 7 | 9 |
| **STRING193** | 5.52E-08 | 68 | 4 | 10 |
| **GEO89** | 2.50E-06 | 101 | 2 | 20 |
| **GEO149** | 8.31E-06 | 145 | 1 | 26 |
| **GEO104** | 1.87E-05 | 33 | 1 | 10 |
| **InWeb47** | 2.25E-05 | 48 | 1 | 10 |
| **GEO102** | 6.78E-05 | 61 | 1 | 15 |
| * Transcriptome significance threshold: p = 3.05E-06, polar metabolome significance threshold: p = 2.27E-04, lipidome significance threshold: p = 2.66E-04  ** Suggestive association: p < 0.05 | | | | |

**Supplementary Table 2. Network associations summary for the LLFS transcriptome-small molecule interaction network (TSI-Net).**

| **Module ID** | **Module P-value** | **Module Size** | **Significant Nodes*** | **Suggestive Nodes**** |
| --- | --- | --- | --- | --- |
| **LLFS12** | 1.30E-28 | 28 | 13 | 6 |
| **LLFS128** | 6.87E-16 | 59 | 2 | 22 |
| **LLFS198** | 4.95E-11 | 122 | 0 | 35 |
| **LLFS117** | 1.28E-10 | 14 | 0 | 8 |
| **LLFS351** | 1.56E-10 | 128 | 3 | 23 |
| **LLFS176** | 8.19E-10 | 15 | 1 | 8 |
| **LLFS137** | 3.25E-09 | 113 | 2 | 24 |
| **LLFS243** | 3.13E-08 | 130 | 2 | 28 |
| **LLFS13** | 3.33E-08 | 13 | 0 | 10 |
| **LLFS125** | 4.73E-08 | 58 | 0 | 21 |
| **LLFS113** | 7.39E-08 | 23 | 0 | 11 |
| **LLFS84** | 3.42E-07 | 132 | 1 | 26 |
| **LLFS38** | 4.25E-07 | 14 | 1 | 8 |
| **LLFS40** | 5.64E-07 | 22 | 1 | 10 |
| **LLFS298** | 2.28E-06 | 118 | 0 | 22 |
| **LLFS15** | 4.11E-06 | 7 | 1 | 5 |
| **LLFS352** | 7.97E-06 | 66 | 1 | 16 |
| **LLFS134** | 1.21E-05 | 117 | 1 | 21 |
| **LLFS3** | 4.34E-05 | 13 | 0 | 5 |
| **LLFS78** | 1.24E-04 | 27 | 2 | 4 |
| * Transcriptome significance threshold: p = 3.05E-06, polar metabolome significance threshold: p = 2.27E-04, lipidome significance threshold: p = 2.66E-04  ** Suggestive association: p < 0.05 | | | | |

**Table 1. Significant TWAS summary statistics for IS**

| **Gene Symbol** | **LLFS P-value** | **LLFS Beta** | **LLFS Standard Error** | **FHS**  **P-value** | **FHS Beta** | **FHS Standard Error** | **TWAS Atlas** |
| --- | --- | --- | --- | --- | --- | --- | --- |
| ***FCER1A*** | 9.87E-12 | 0.449 | 0.051 | 9.65E-14 | 0.375 | 0.050 | - |
| ***CPA3*** | 1.71E-11 | 0.400 | 0.046 | 6.71E-15 | 0.336 | 0.043 | Fasting Glucose |
| ***GATA2*** | 7.17E-11 | 0.236 | 0.028 | 1.41E-13 | 0.254 | 0.034 | Fasting Glucose, Fasting Insulin |
| ***HDC*** | 1.14E-10 | 0.277 | 0.033 | 1.84E-13 | 0.244 | 0.033 | Fasting Glucose |
| ***SLC45A3*** | 1.52E-10 | 0.368 | 0.045 | 3.69E-10 | 0.287 | 0.045 | - |
| ***ABCG1*** | 2.94E-10 | 0.549 | 0.068 | 1.49E-08 | 0.458 | 0.080 | - |
| ***MS4A2*** | 9.02E-10 | 0.354 | 0.045 | 9.12E-11 | 0.337 | 0.052 | Fasting Glucose |
| ***AKAP12*** | 9.73E-10 | 0.355 | 0.045 | 8.48E-14 | 0.337 | 0.045 | Fasting Glucose |
| ***ITGB8*** | 1.83E-08 | 0.353 | 0.049 | N/A | N/A | - | - |
| ***CX3CR1*** | 1.85E-07 | -0.646 | 0.099 | 8.64E-06 | -0.4444 | 0.100 | - |
| ***MBNL3*** | 3.06E-07 | 0.434 | 0.066 | 0.782 | -0.0385 | 0.139 | - |
| ***PTGER2*** | 5.32E-07 | -0.805 | 0.128 | 1.55E-08 | -0.768 | 0.135 | Fasting Glucose |
| ***LINC02458*** | 5.39E-07 | 0.418 | 0.065 | 1.51E-12 | 0.474 | 0.66 | - |
| ***CCDC71*** | 9.71E-07 | 1.087 | 0.174 | 7.15E-5 | 0.809 | 0.203 | - |
| ***KLHDC8B*** | 1.77E-06 | 0.419 | 0.069 | 5.62E-5 | 0.291 | 0.072 | - |
| ***LTF*** | 2.27E-06 | -0.175 | 0.029 | 0.0925 | -0.0434 | 0.026 | - |
| ***GCSAML*** | 2.52E-06 | 0.446 | 0.074 | 9.70E-12 | 0.503 | 0.073 | - |
| ***ENPP3*** | 2.62E-06 | 0.413 | 0.069 | 2.61E-11 | 0.502 | 0.075 | - |

**Supplementary Figure 1. Distribution patterns of small molecules (SMs) according to their category.** In each panel, plots represent the raw, semi-processed (for technical variables), and processed peak areas from left to right. Only one representative of each category is illustrated.

**a) Acylcarnitines:**

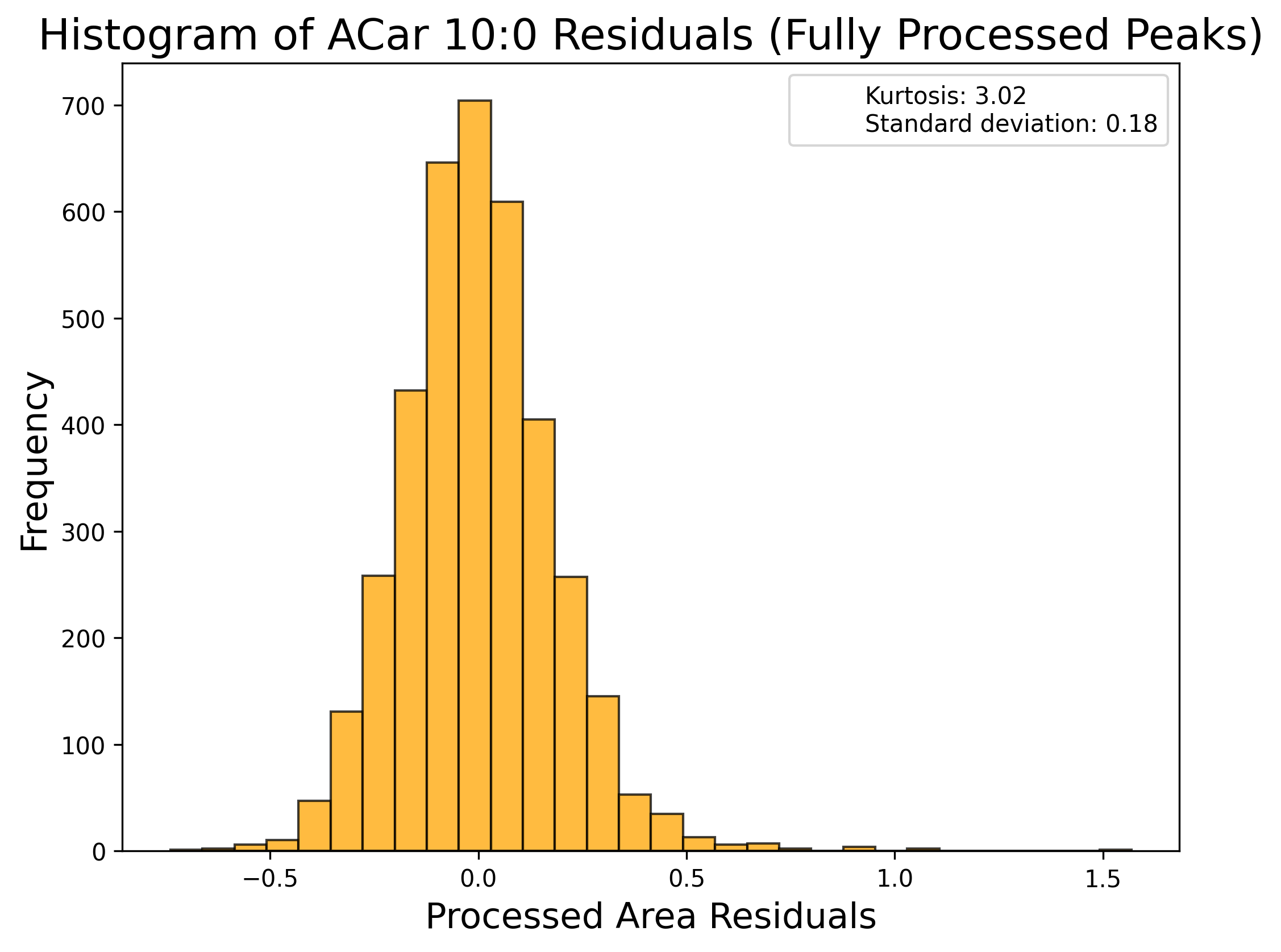

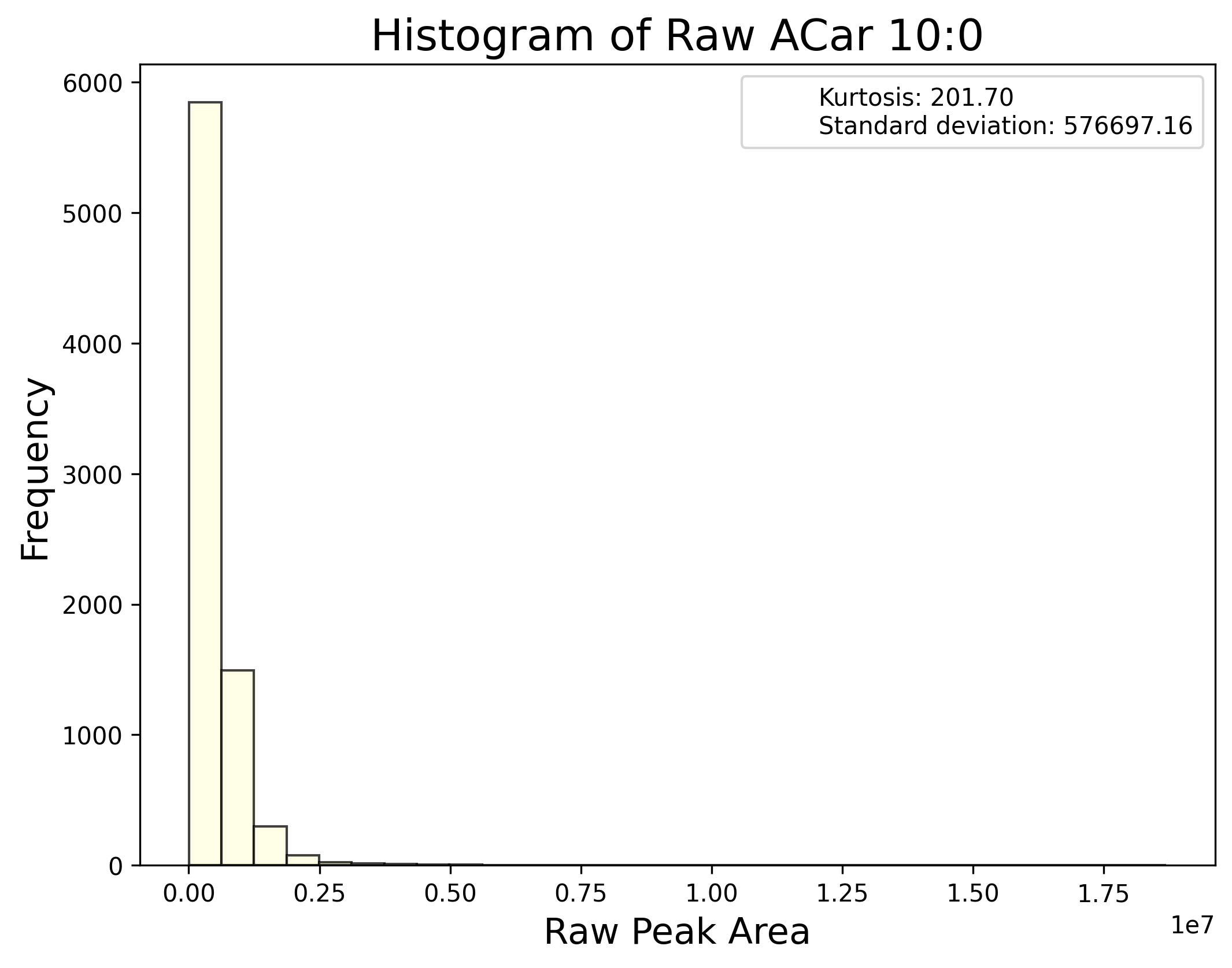

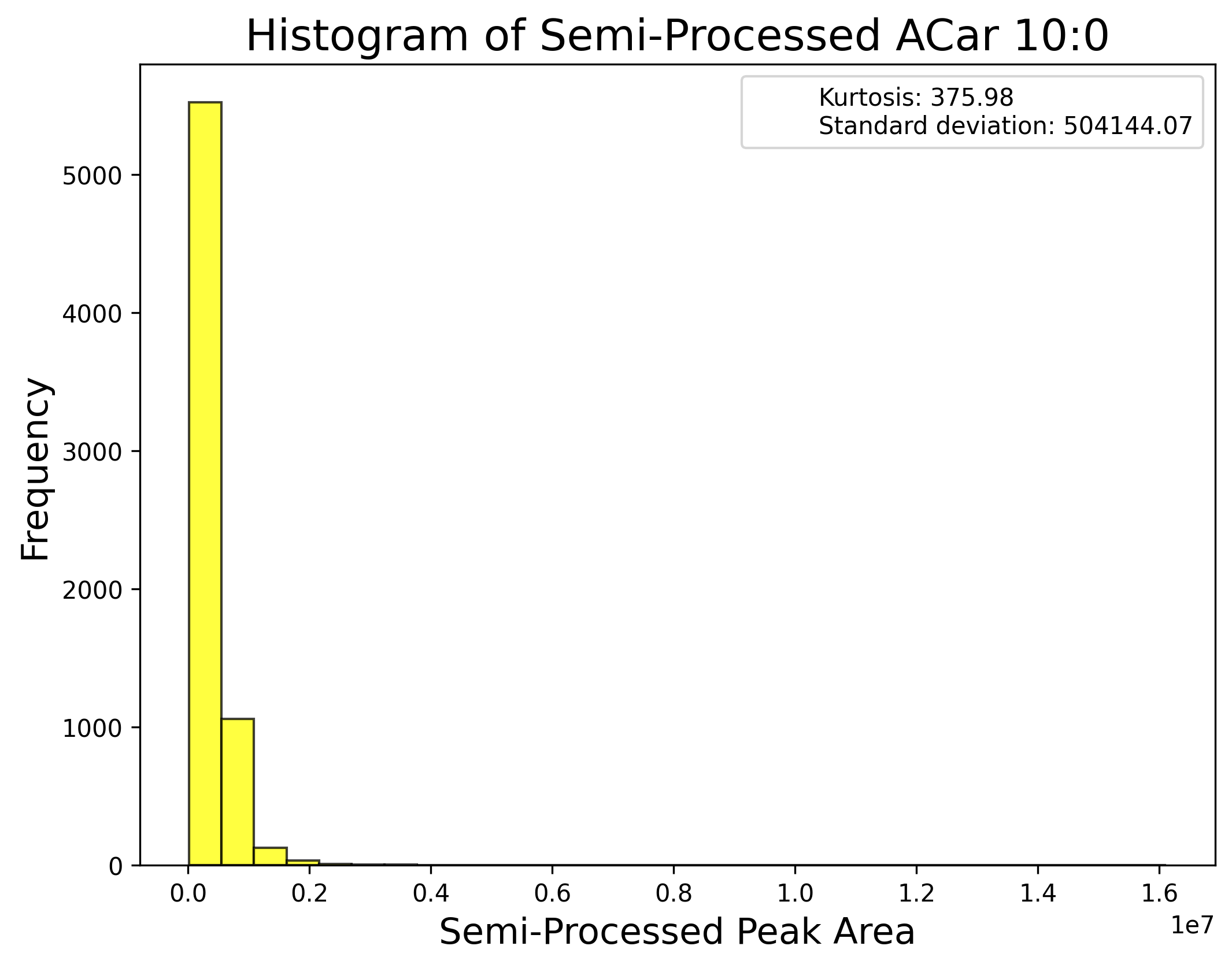

**b) Ceramides:**

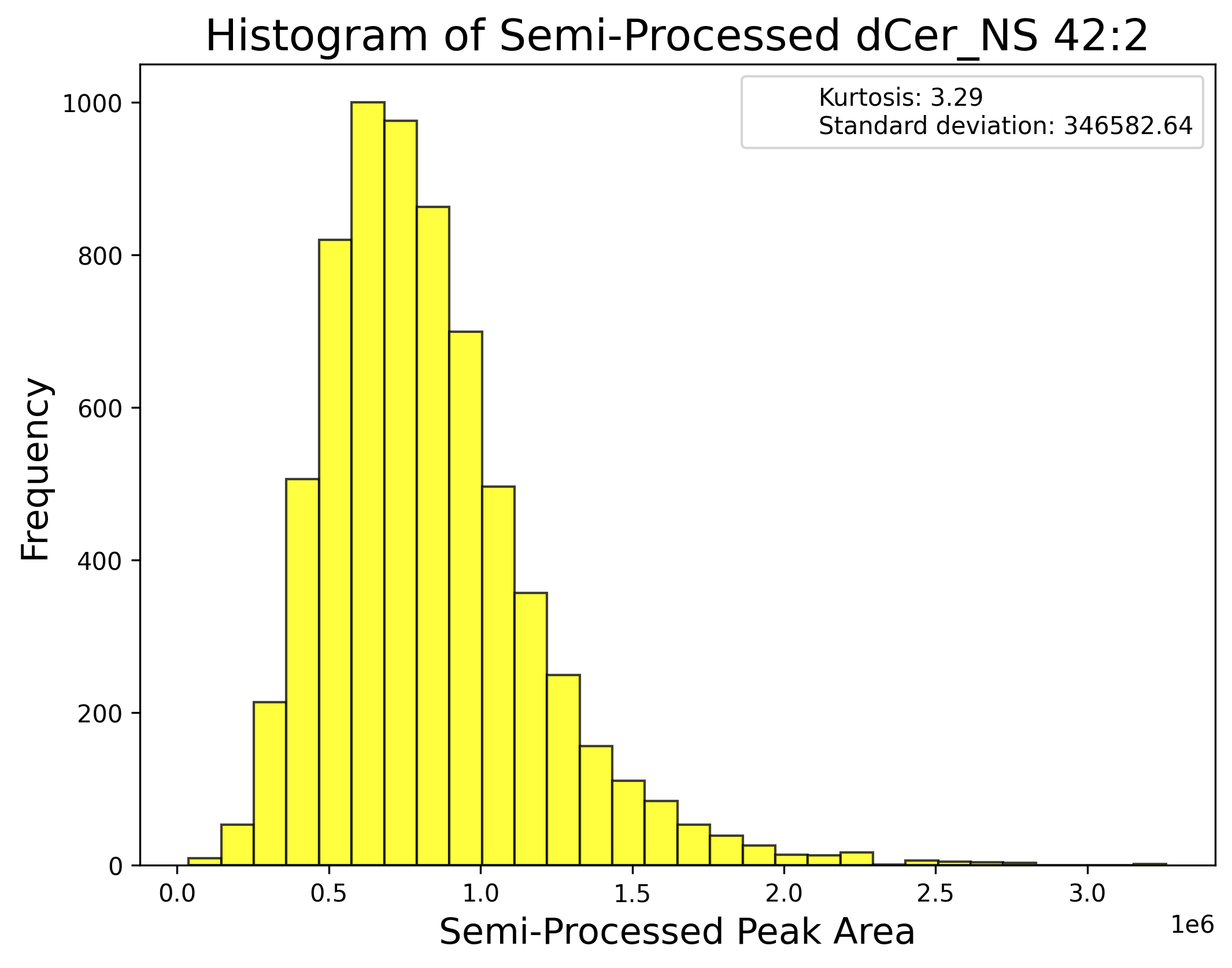

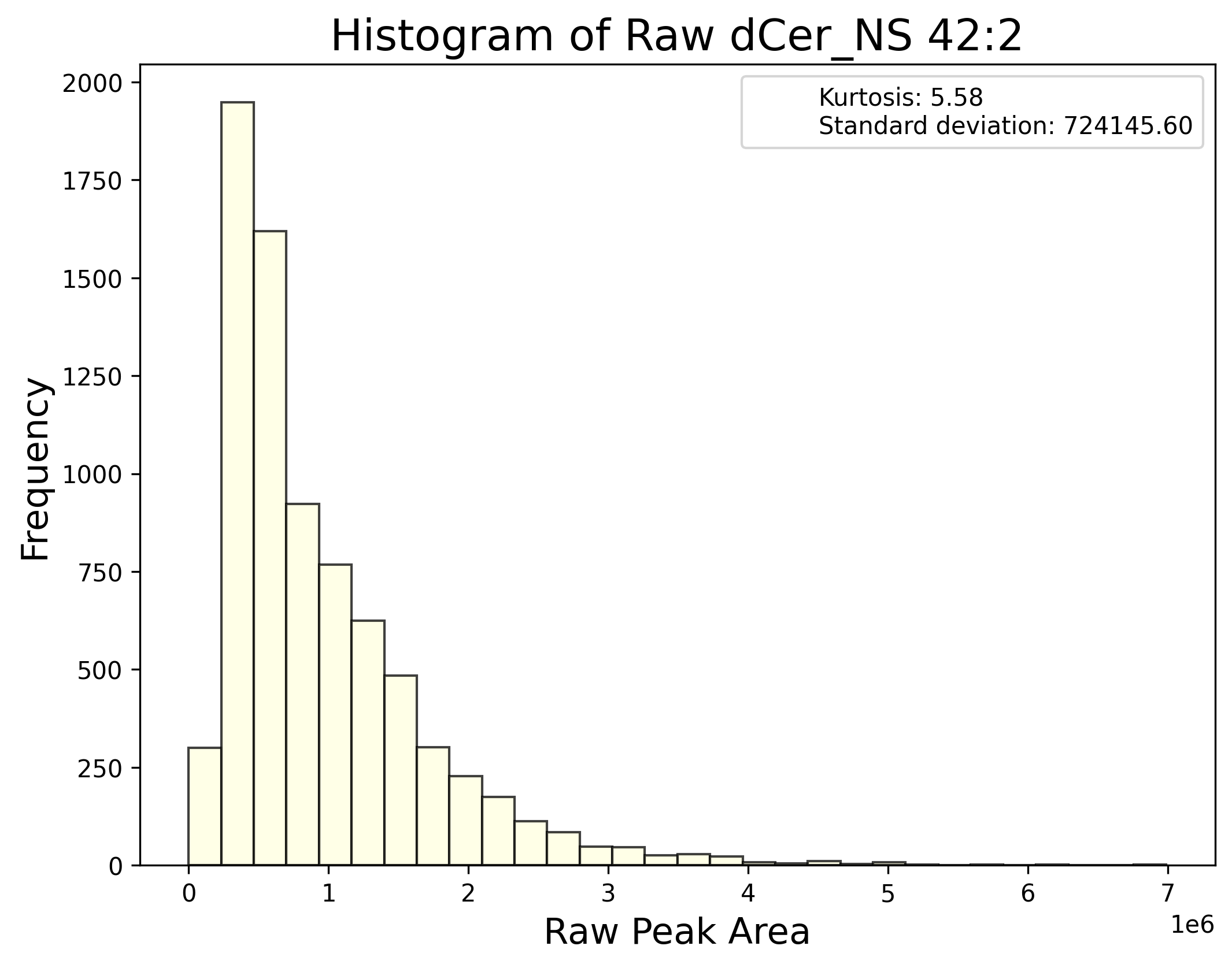

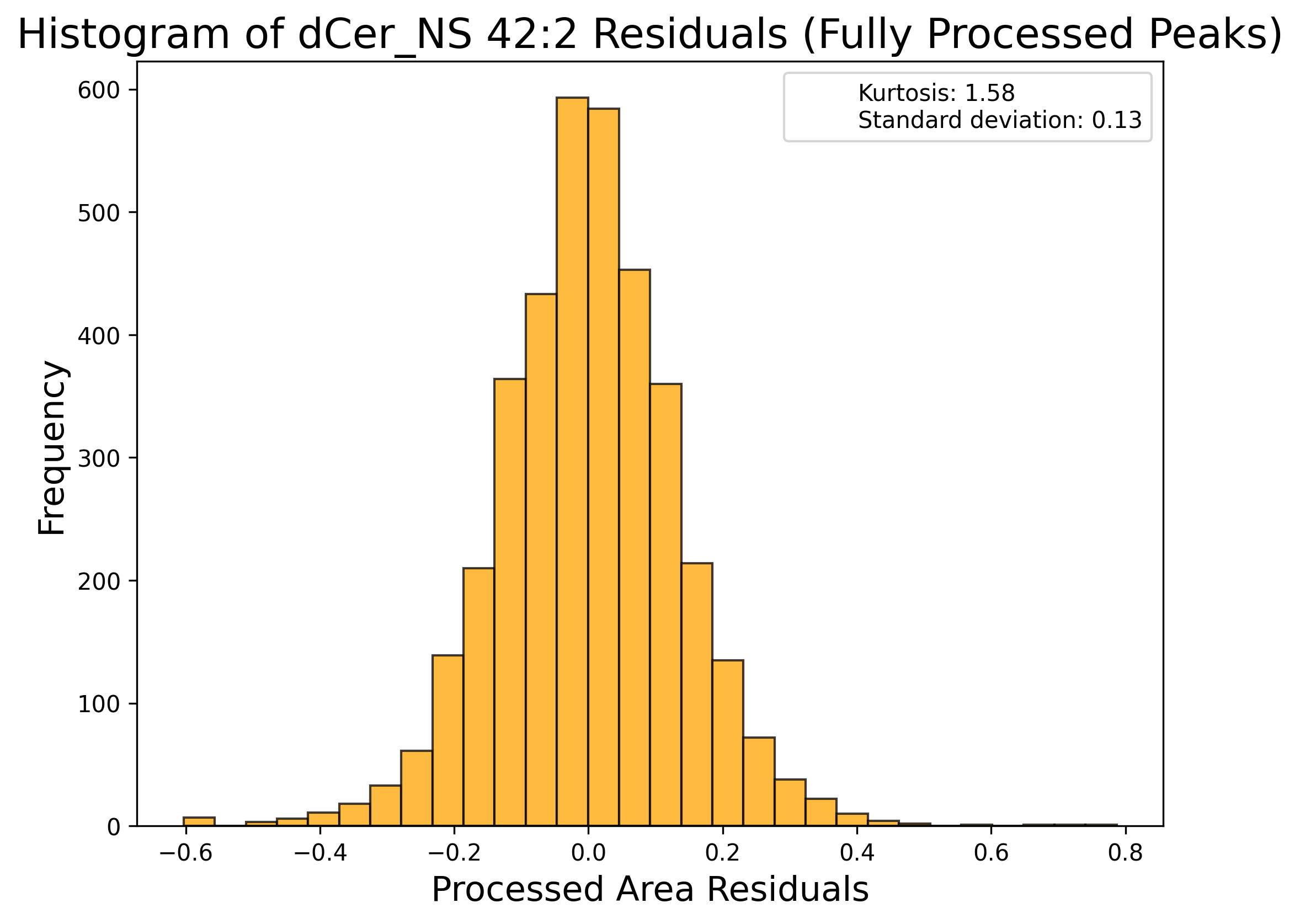

**c) Cholesterol esters:**

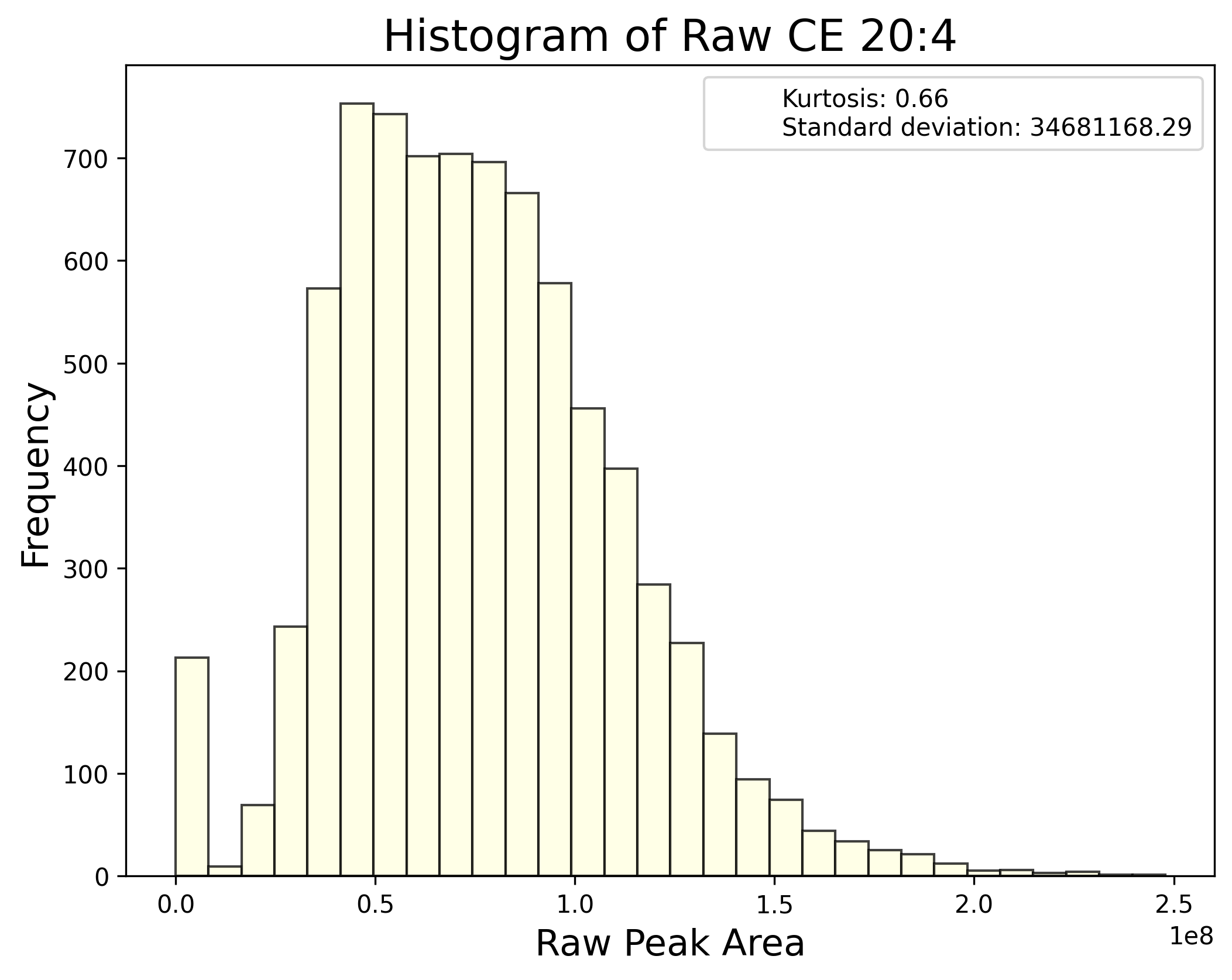

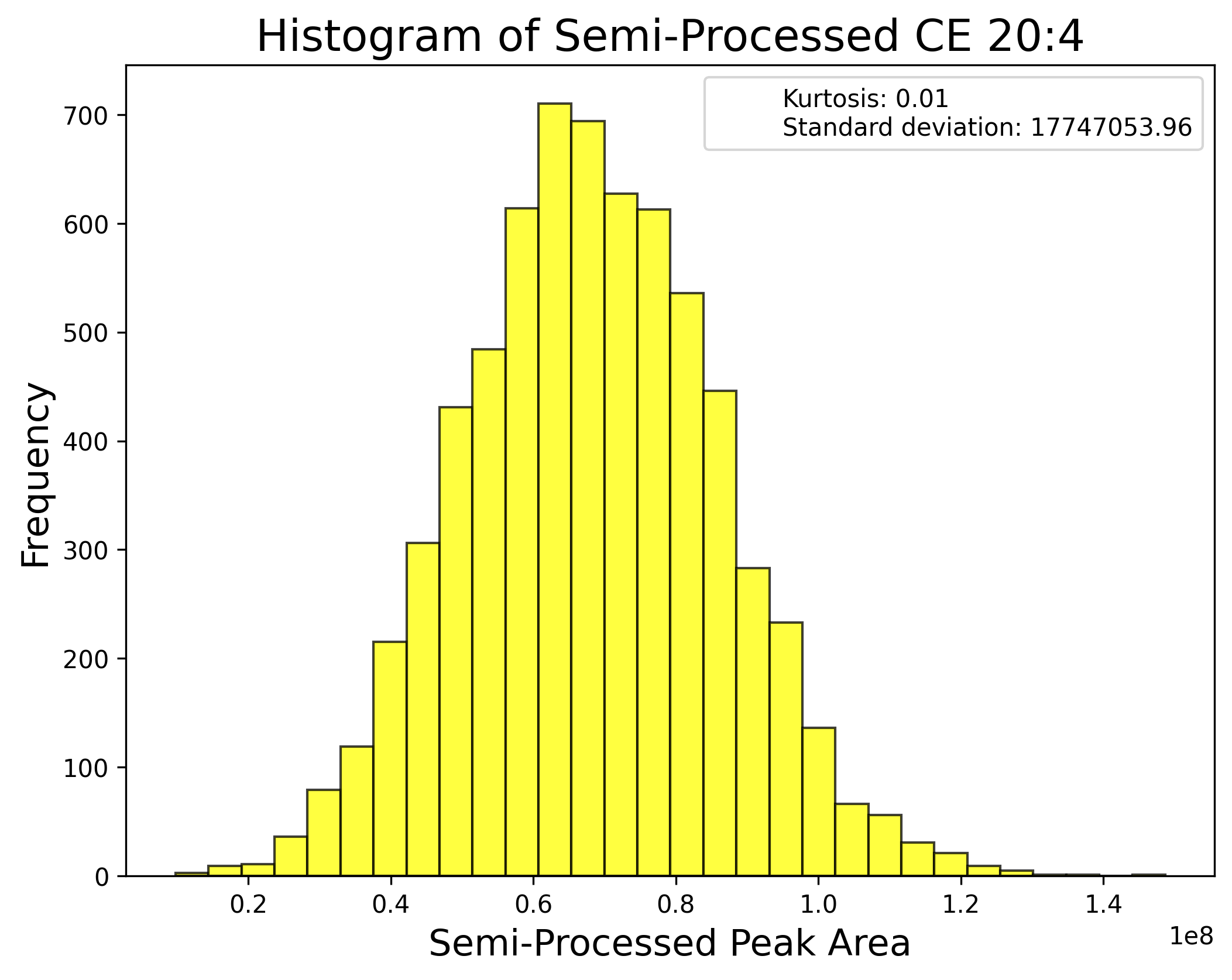

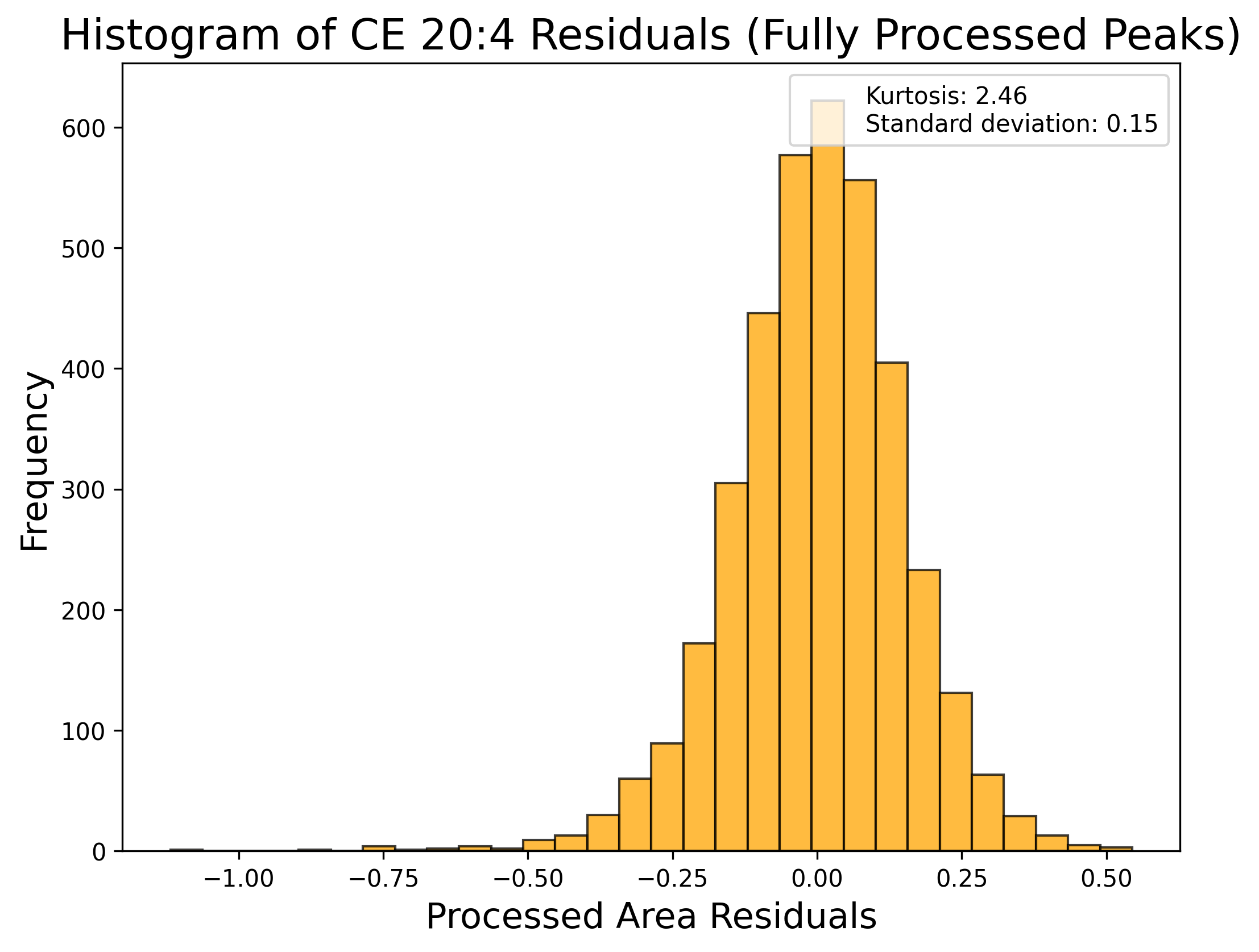

**d) Diacylglycerol:**

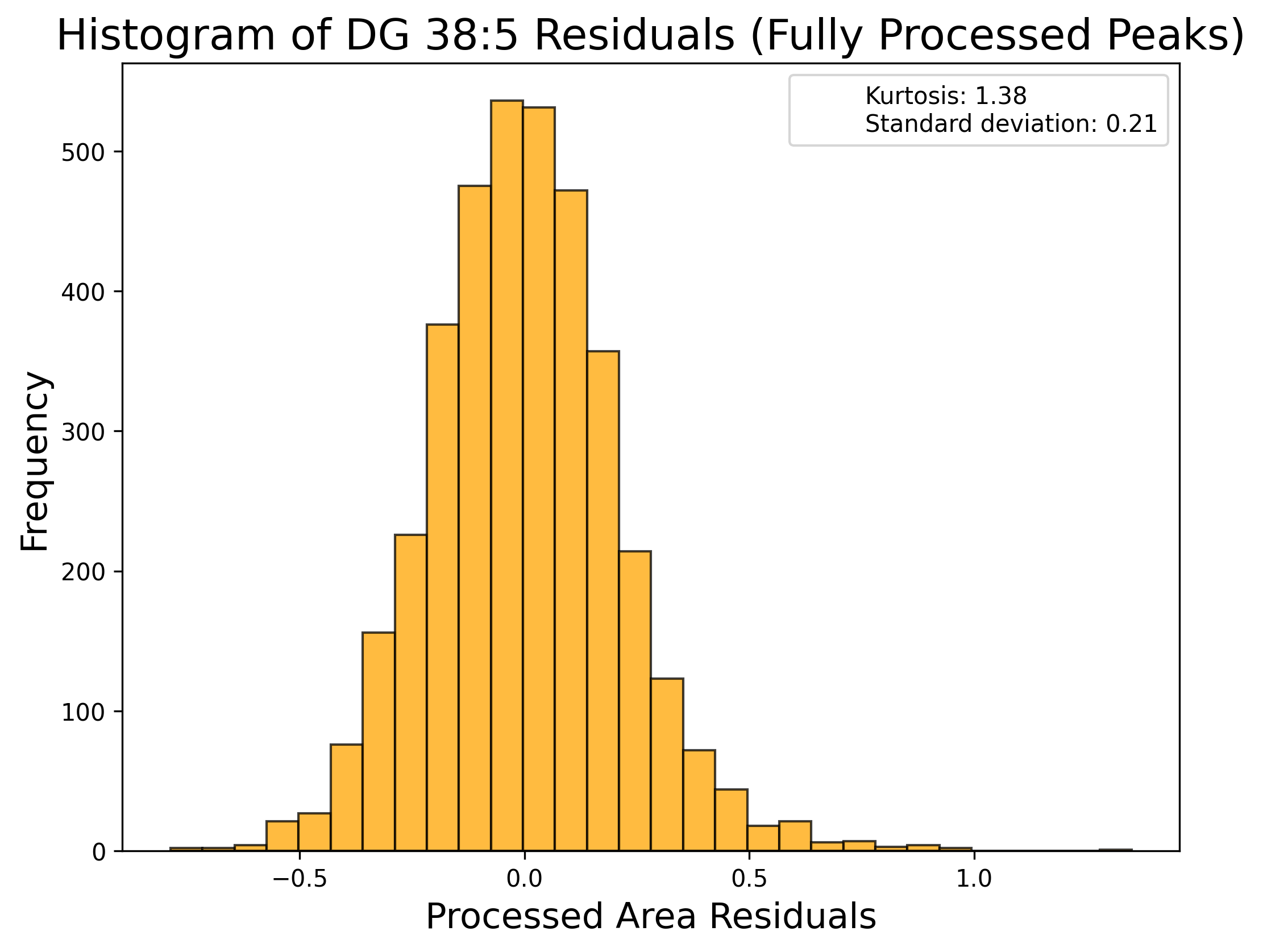

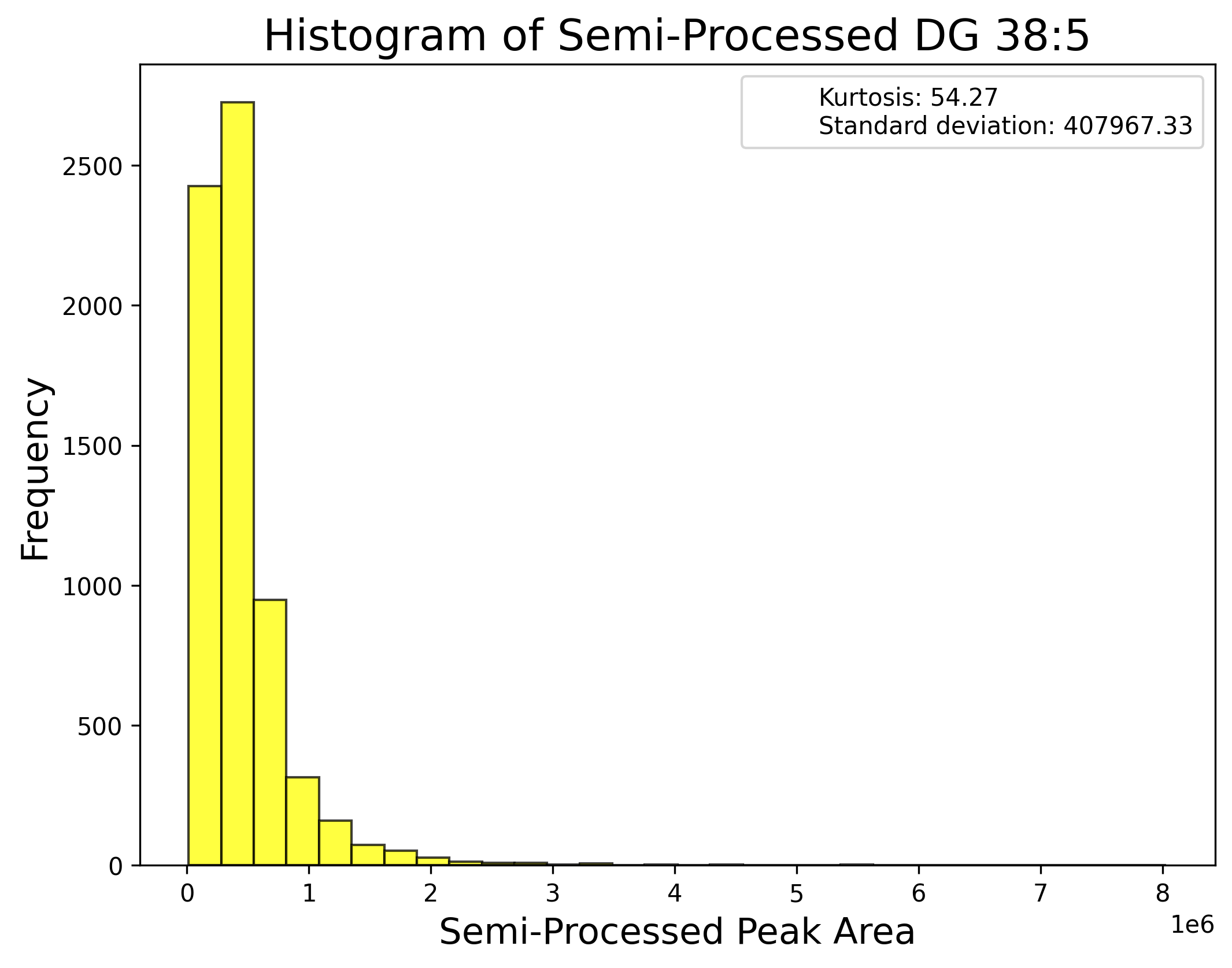

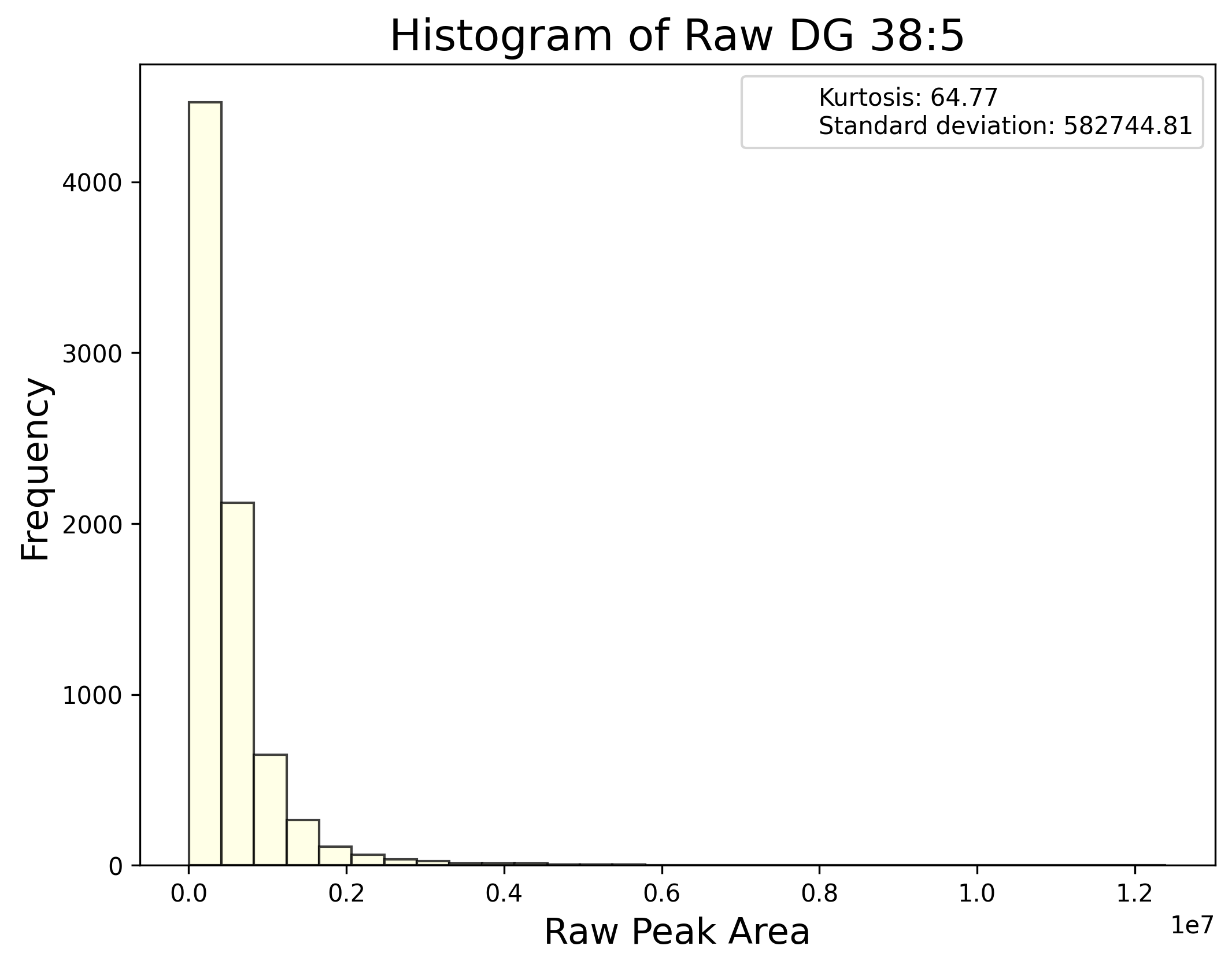

**e) Hexosylceramide:**

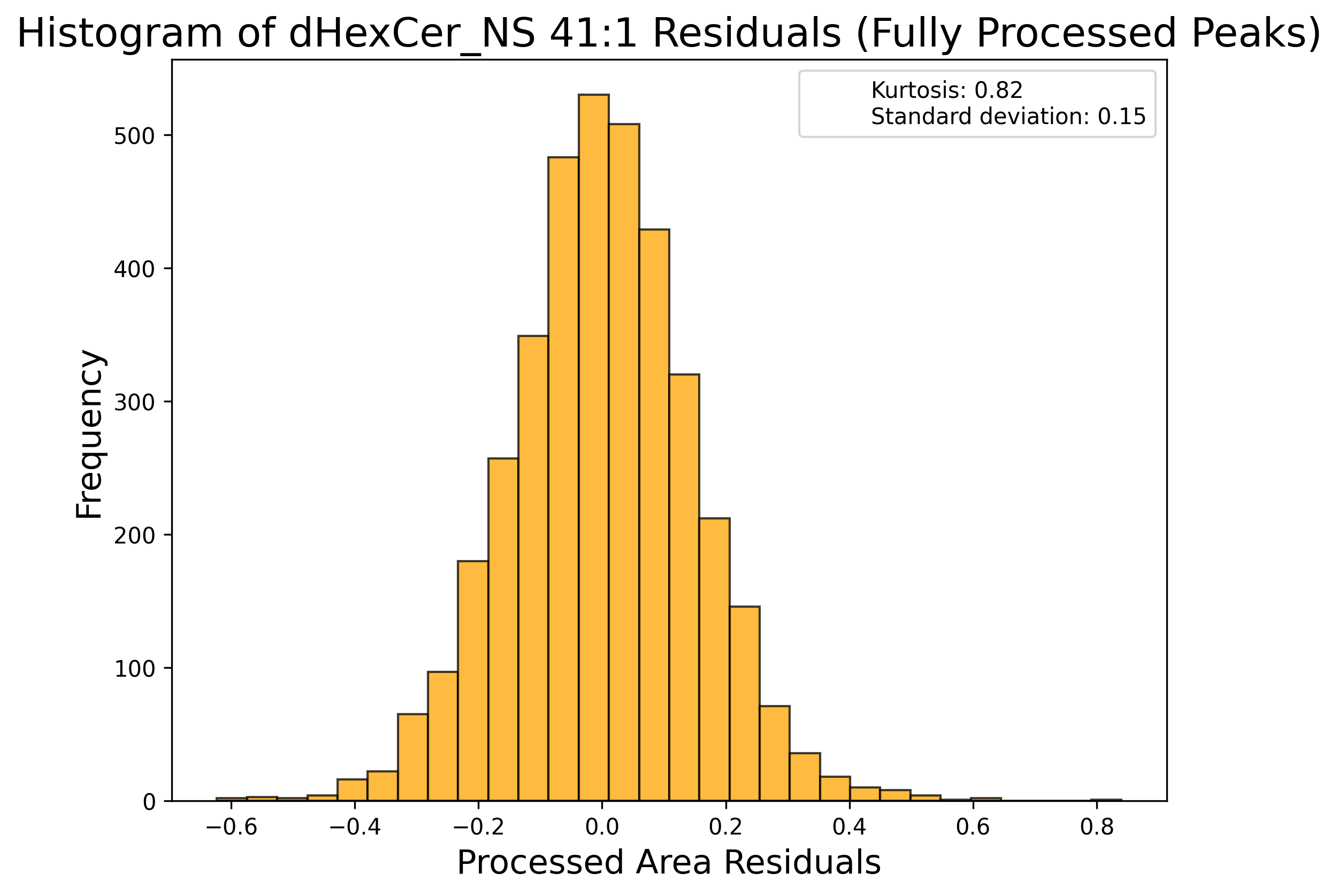

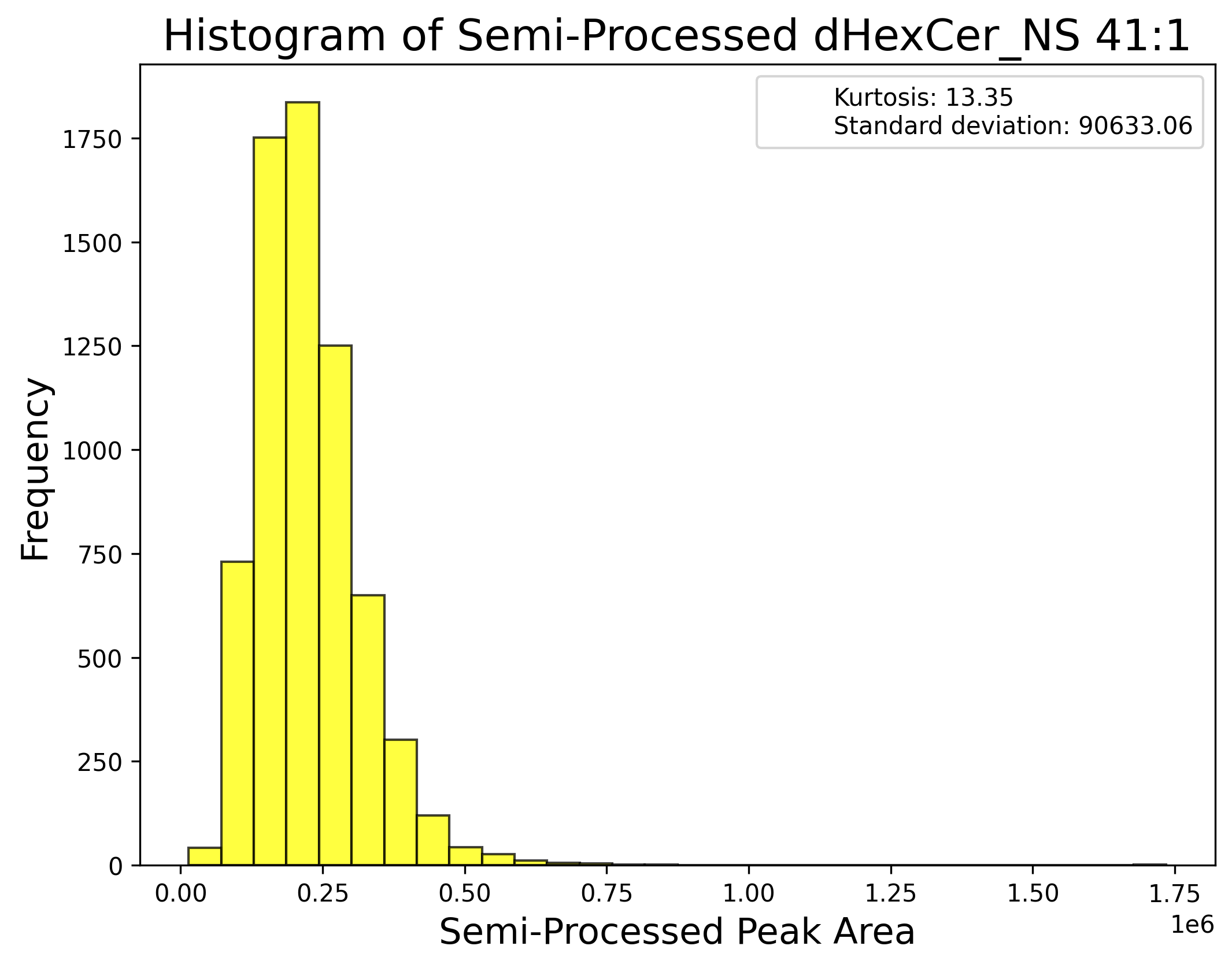

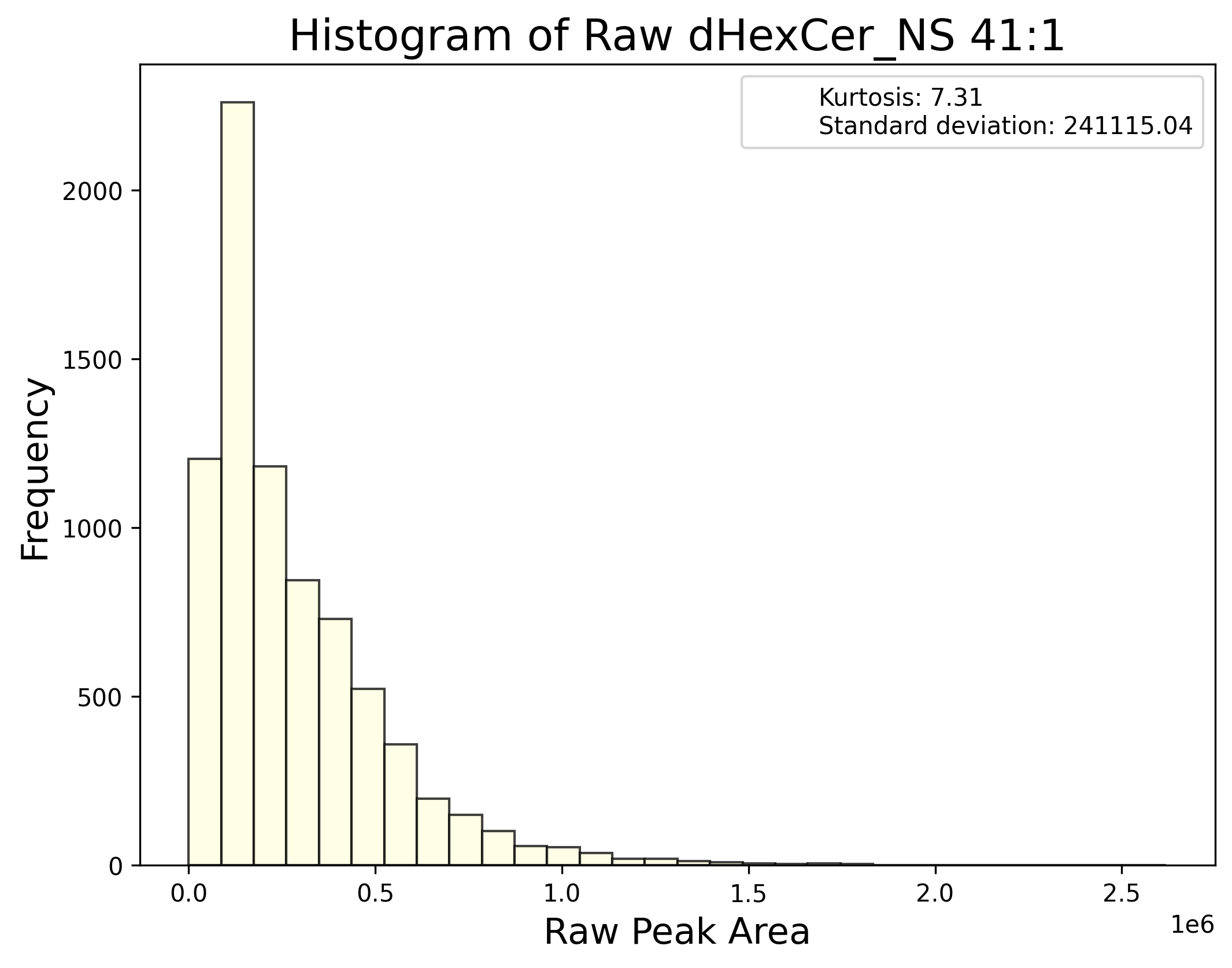

**f) Lysophosphatidylcholine:**

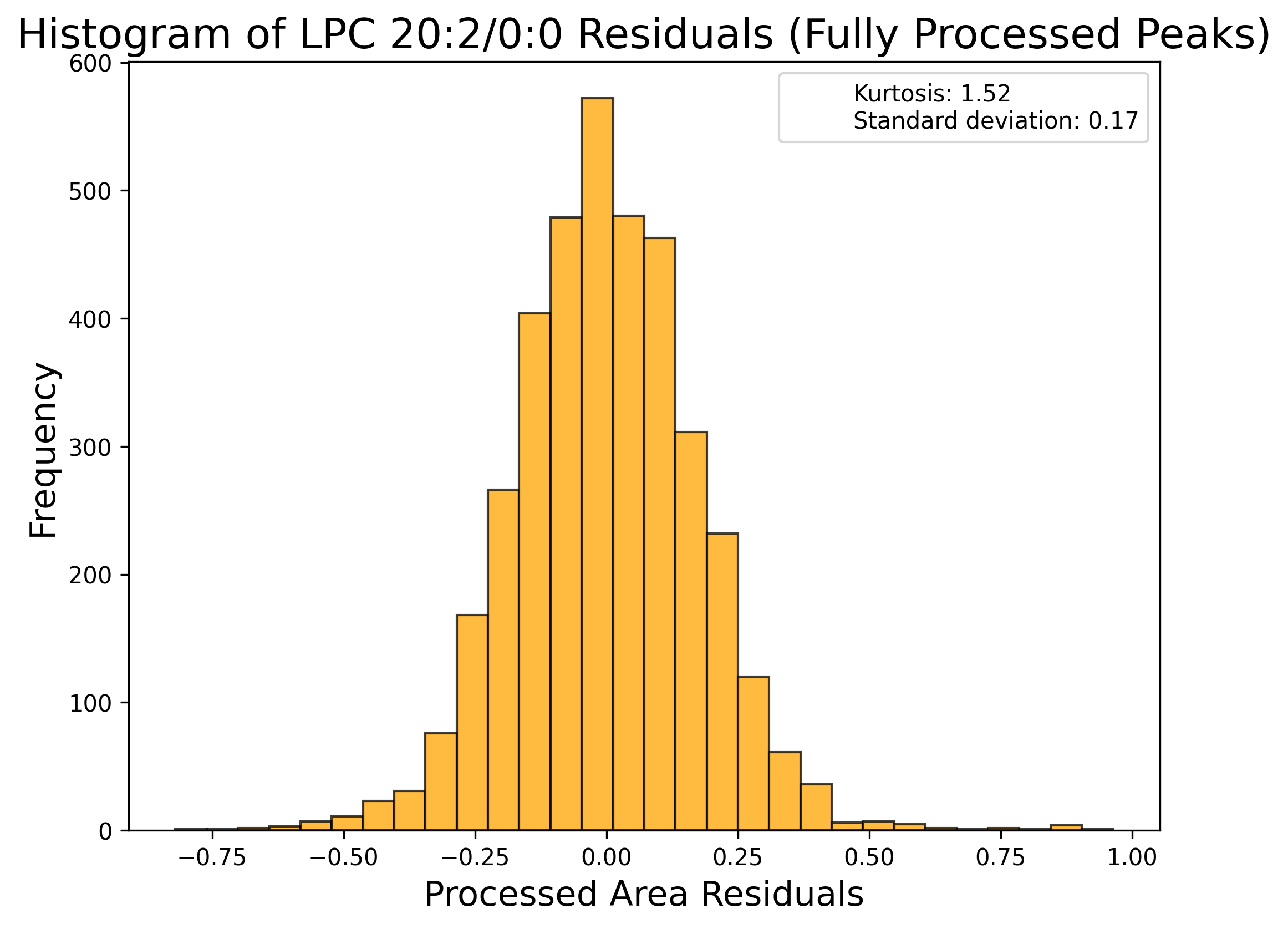

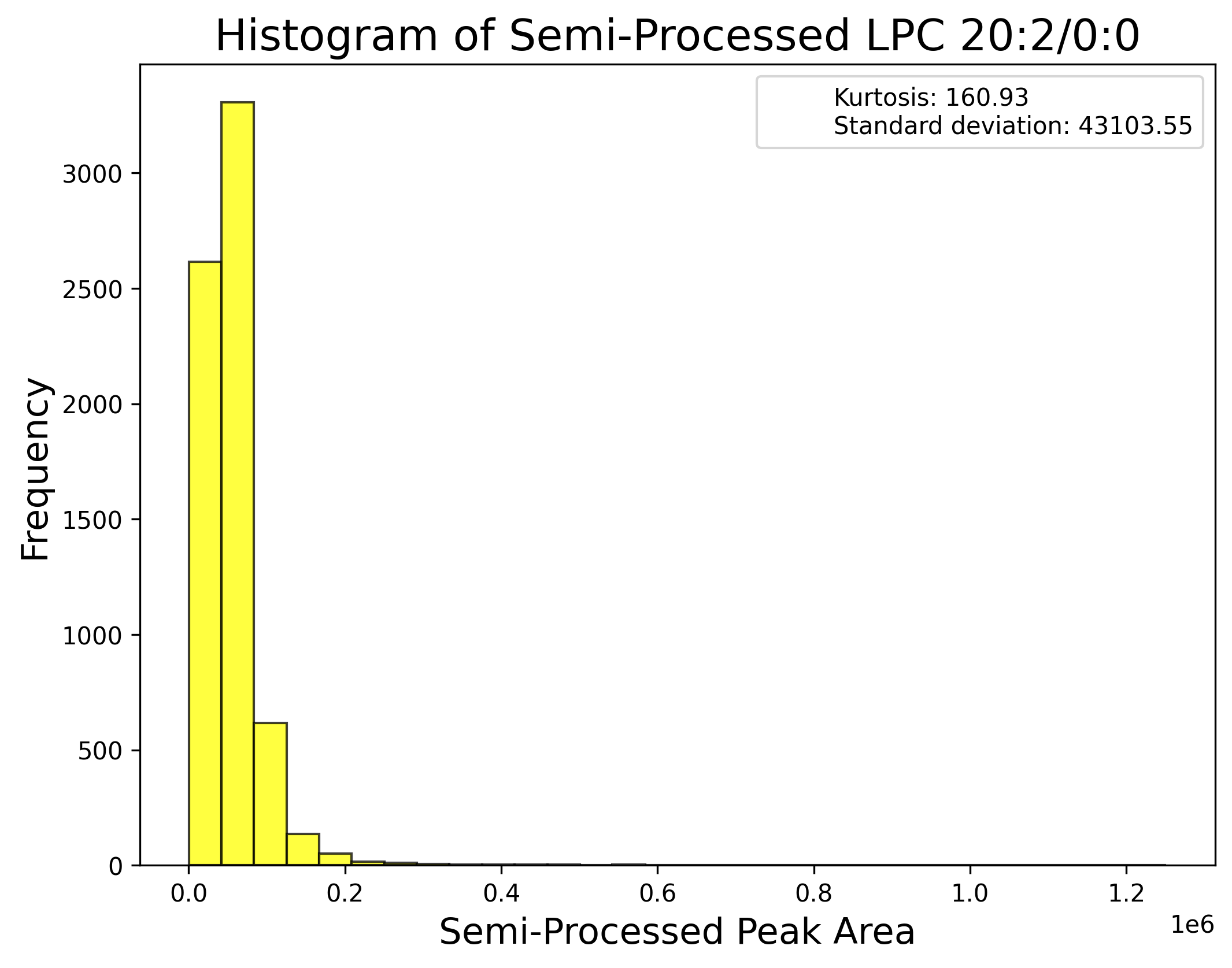

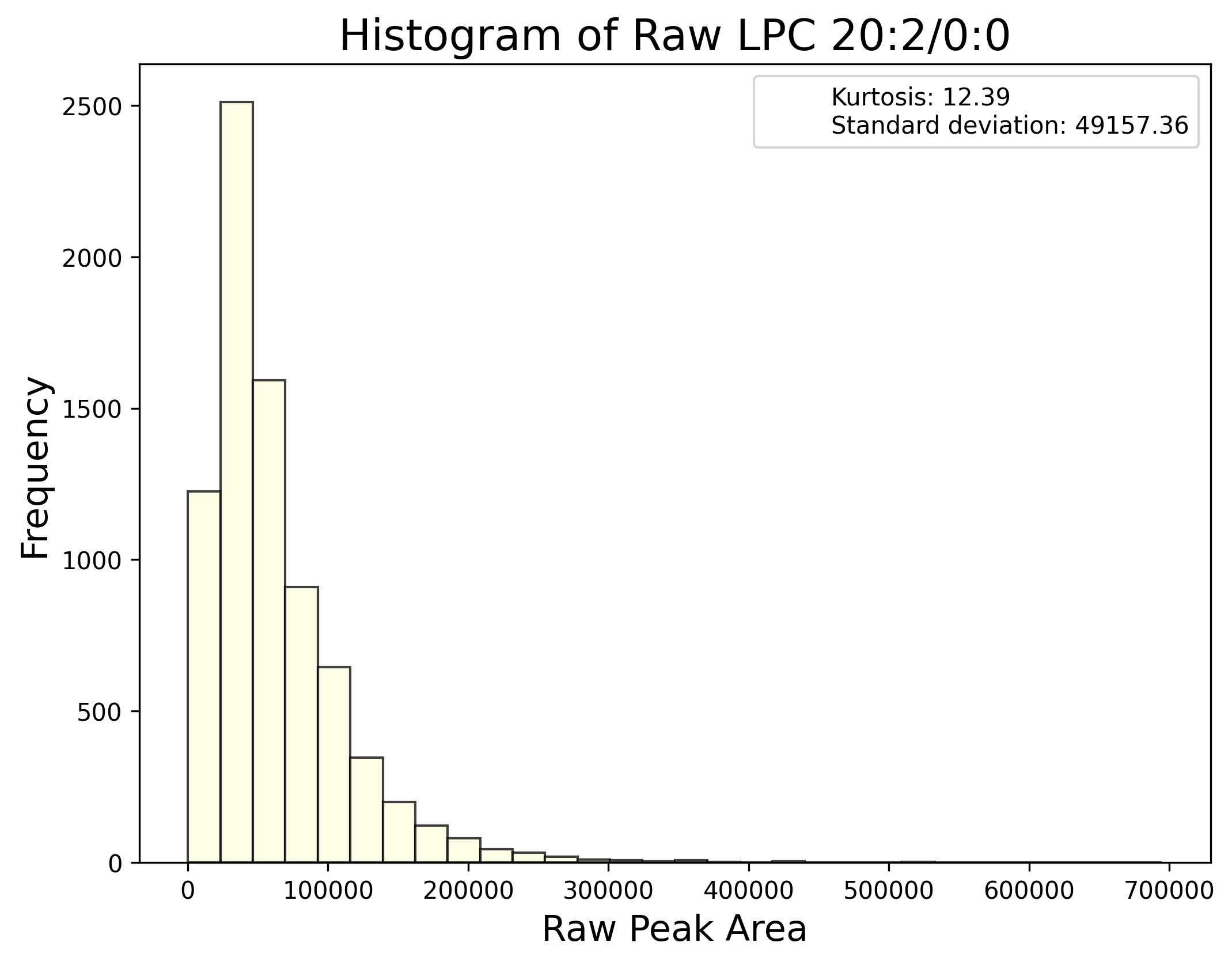

**g) Lysophosphatidylethanolamine:**

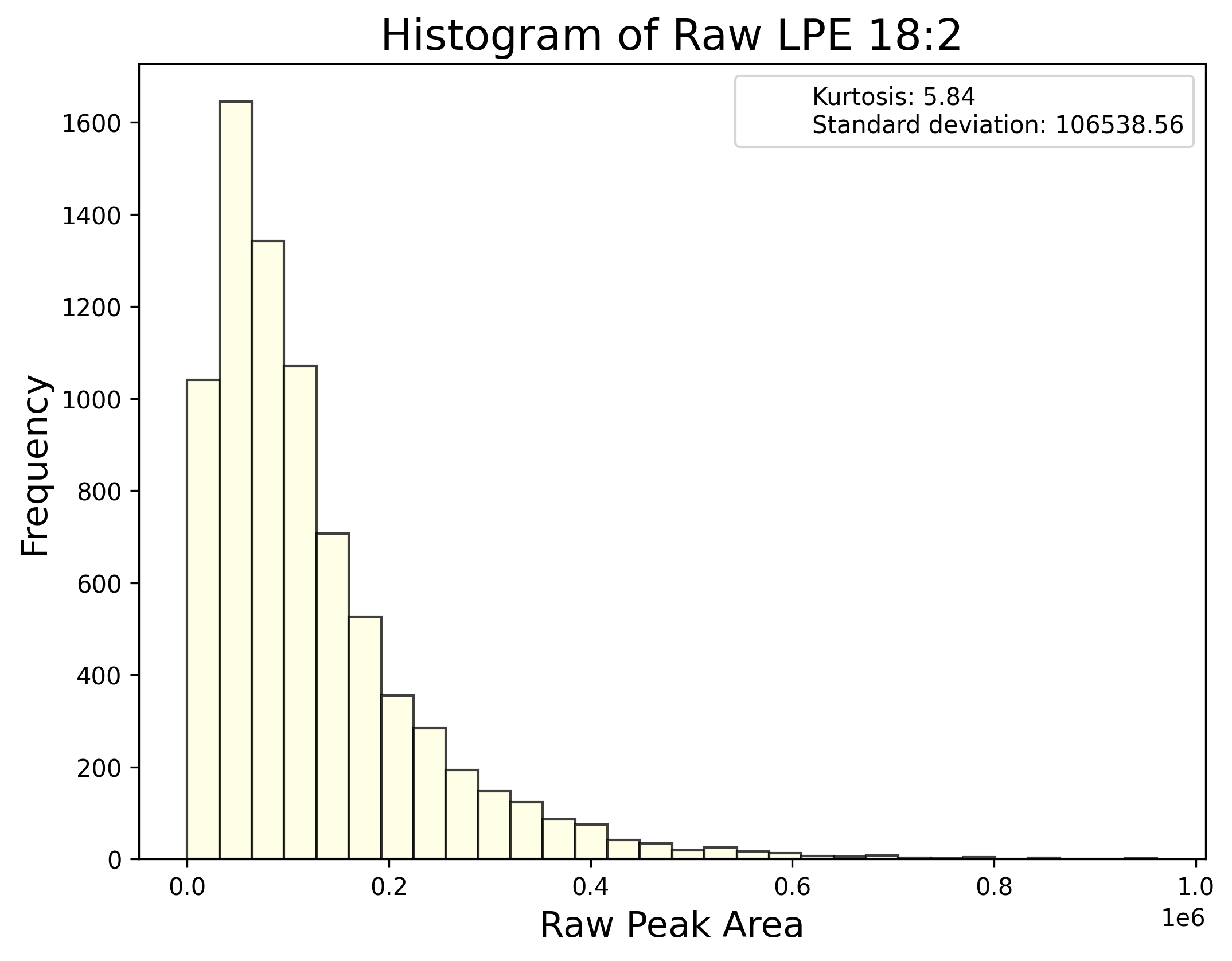

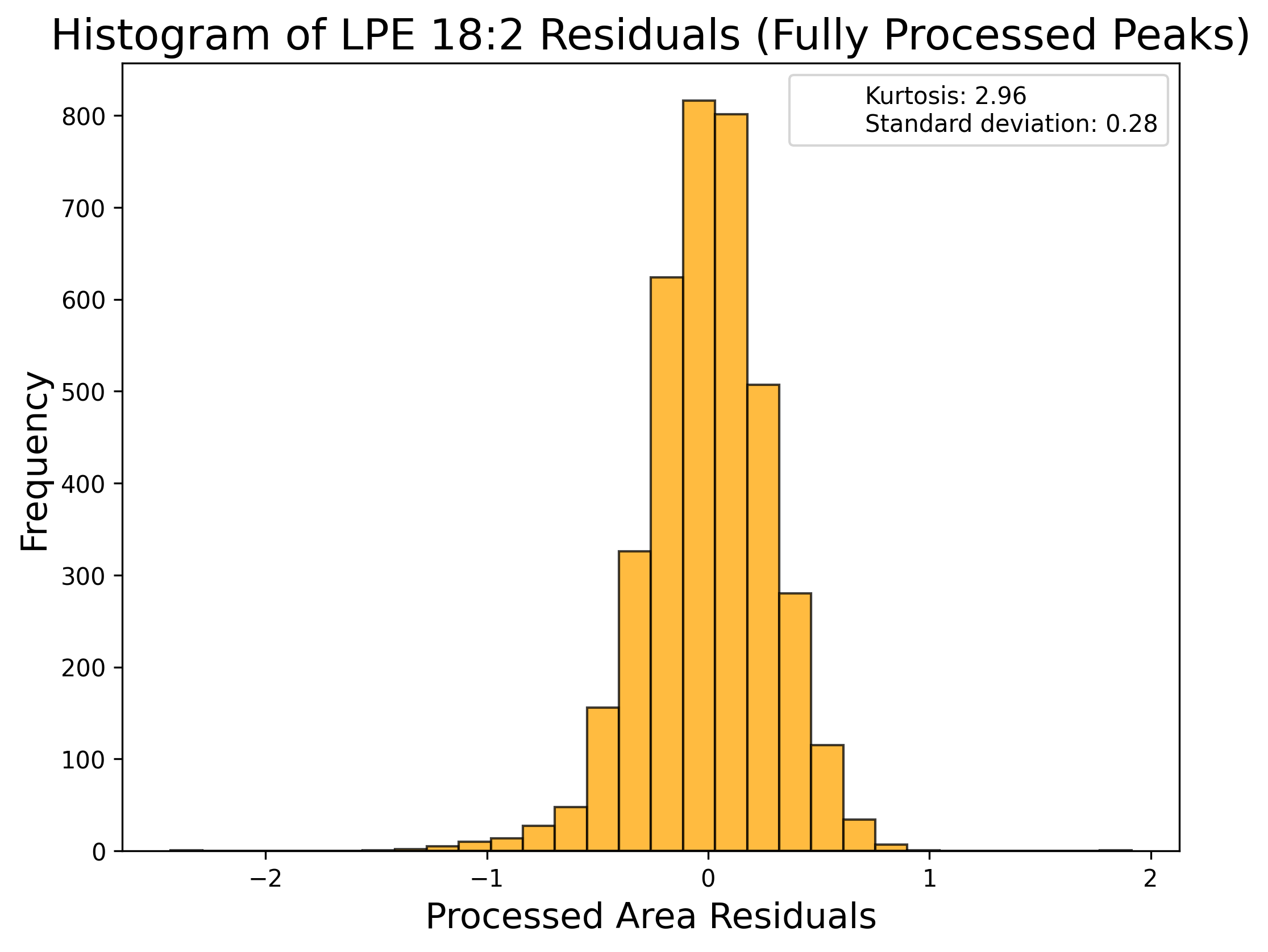

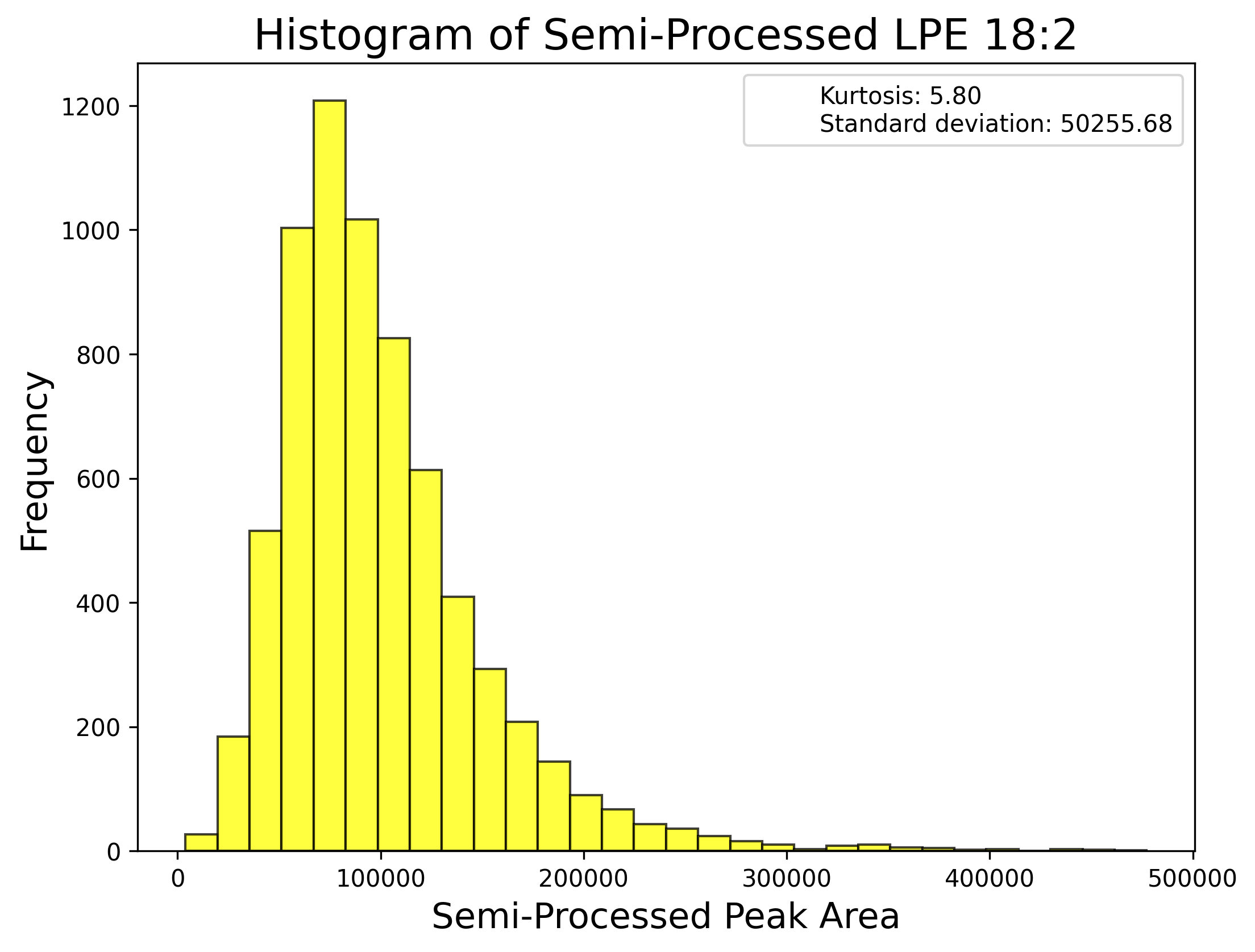

**h) Phosphatidylcholine:**

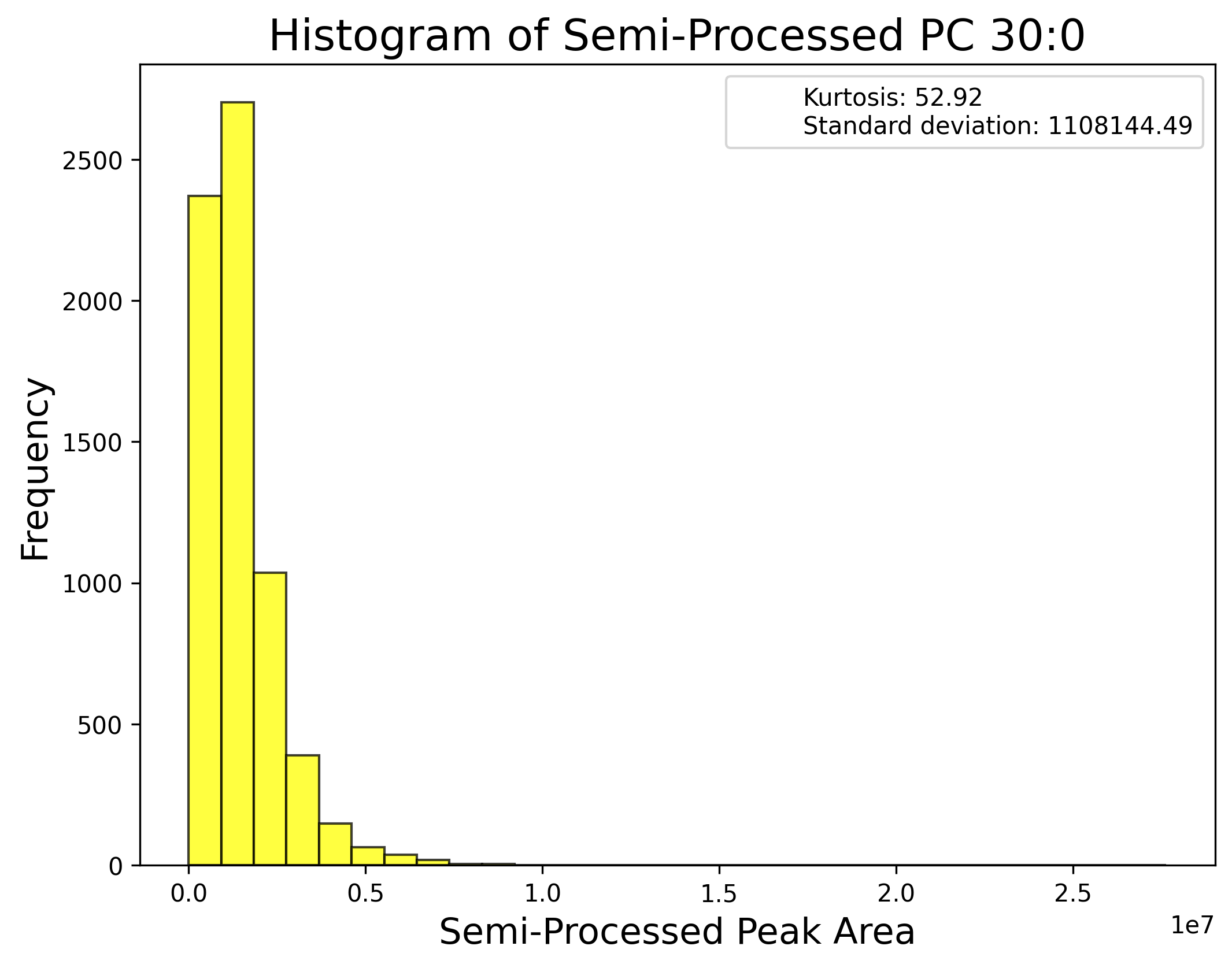

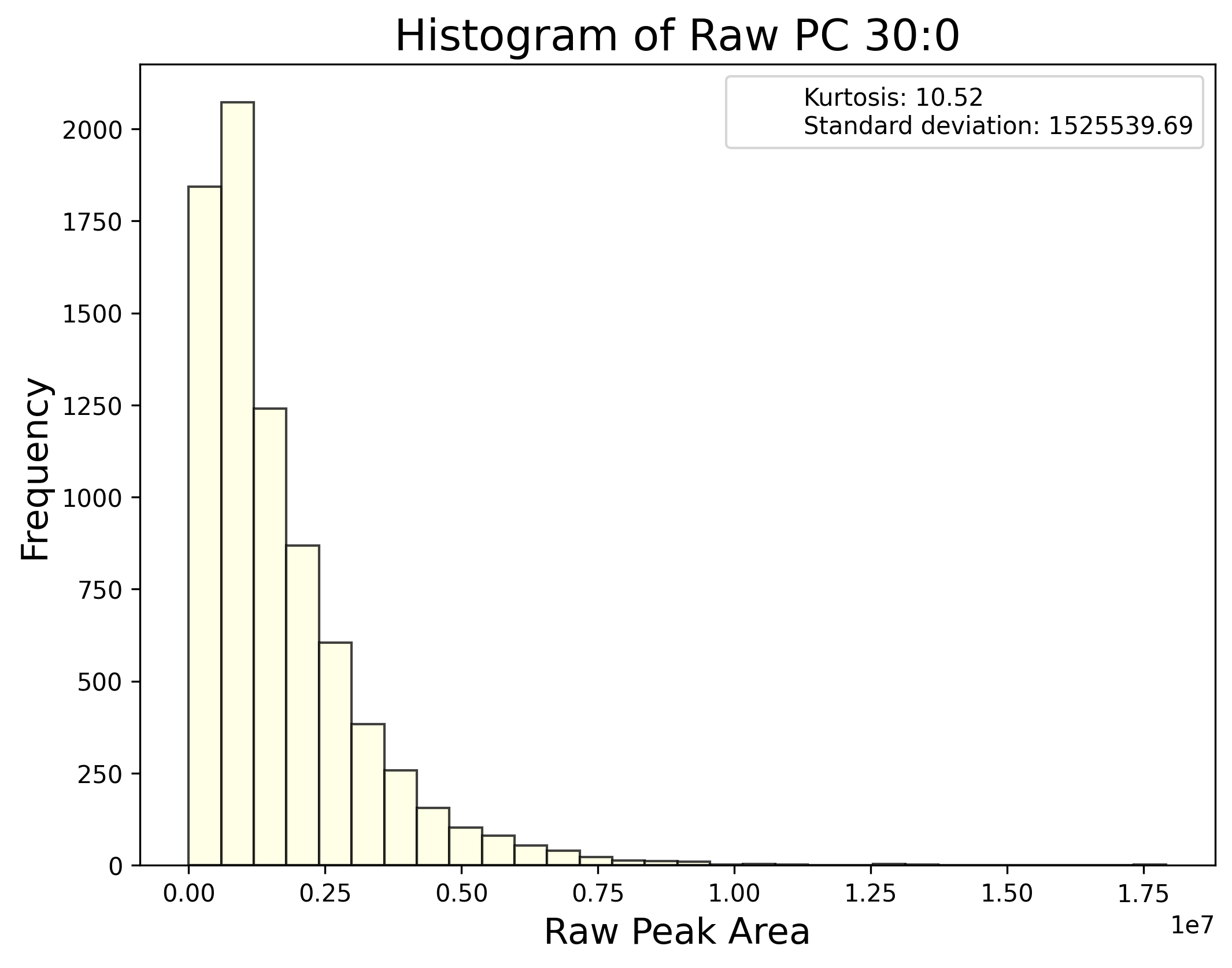

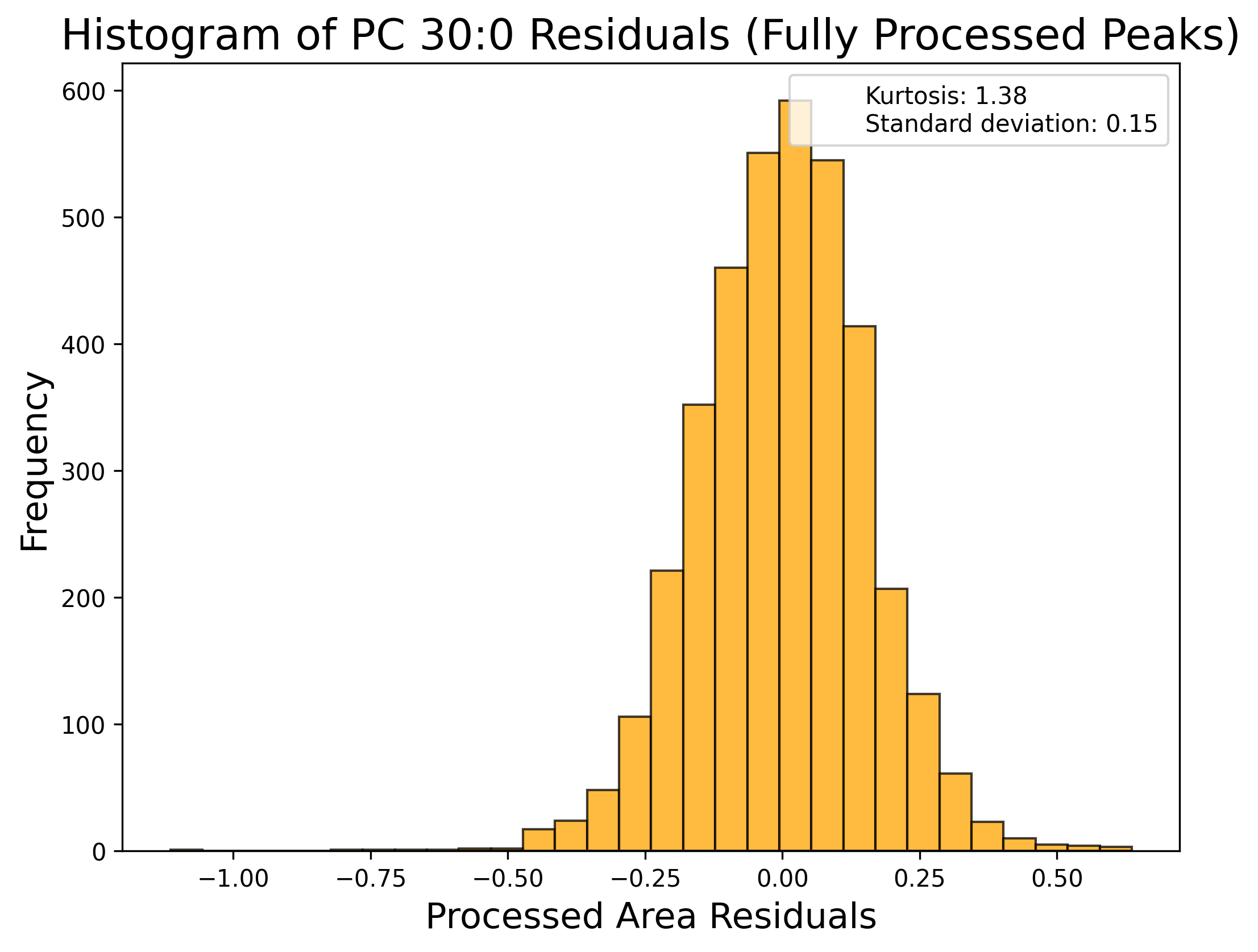

**i) Phosphatidylethanolamine:**

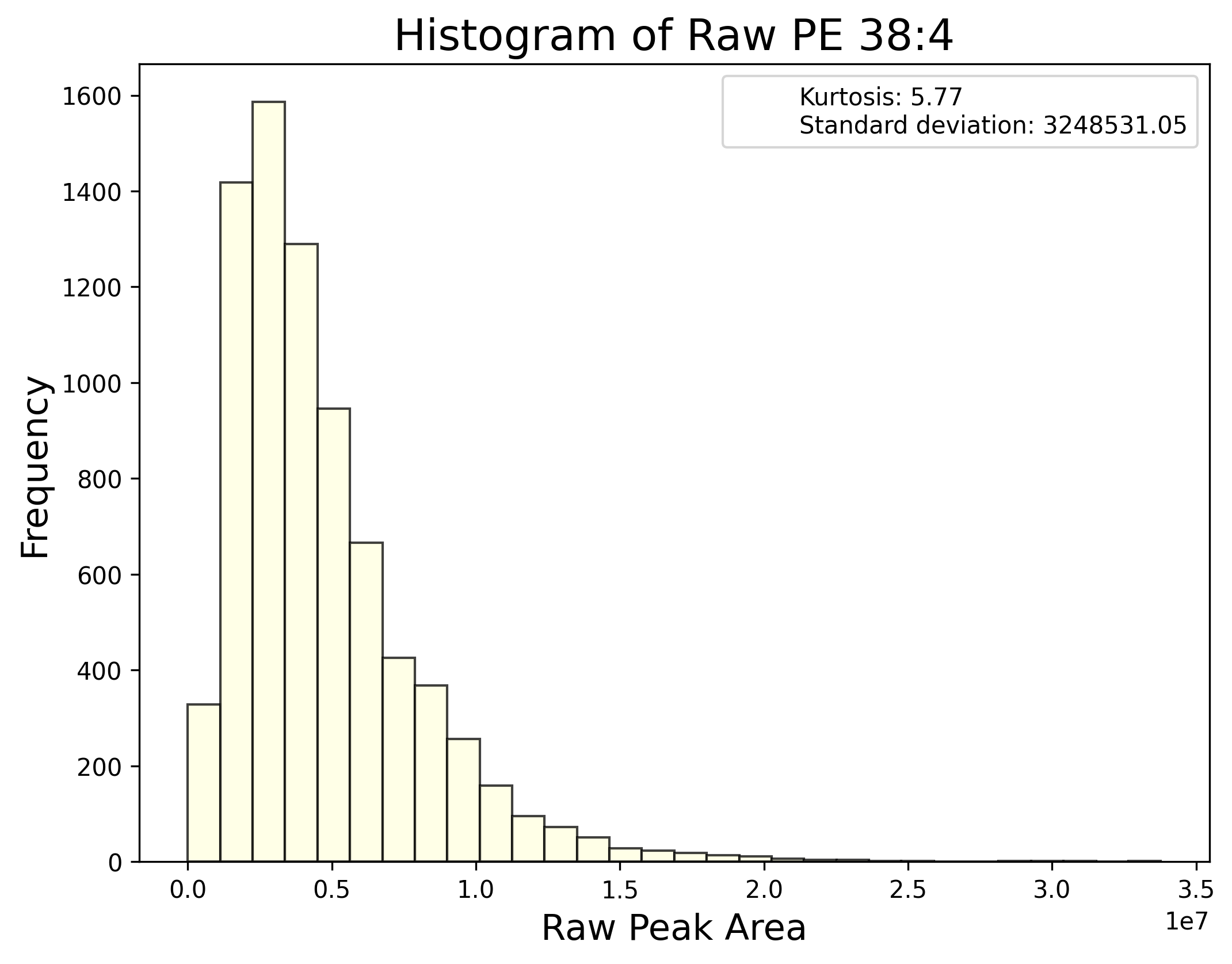

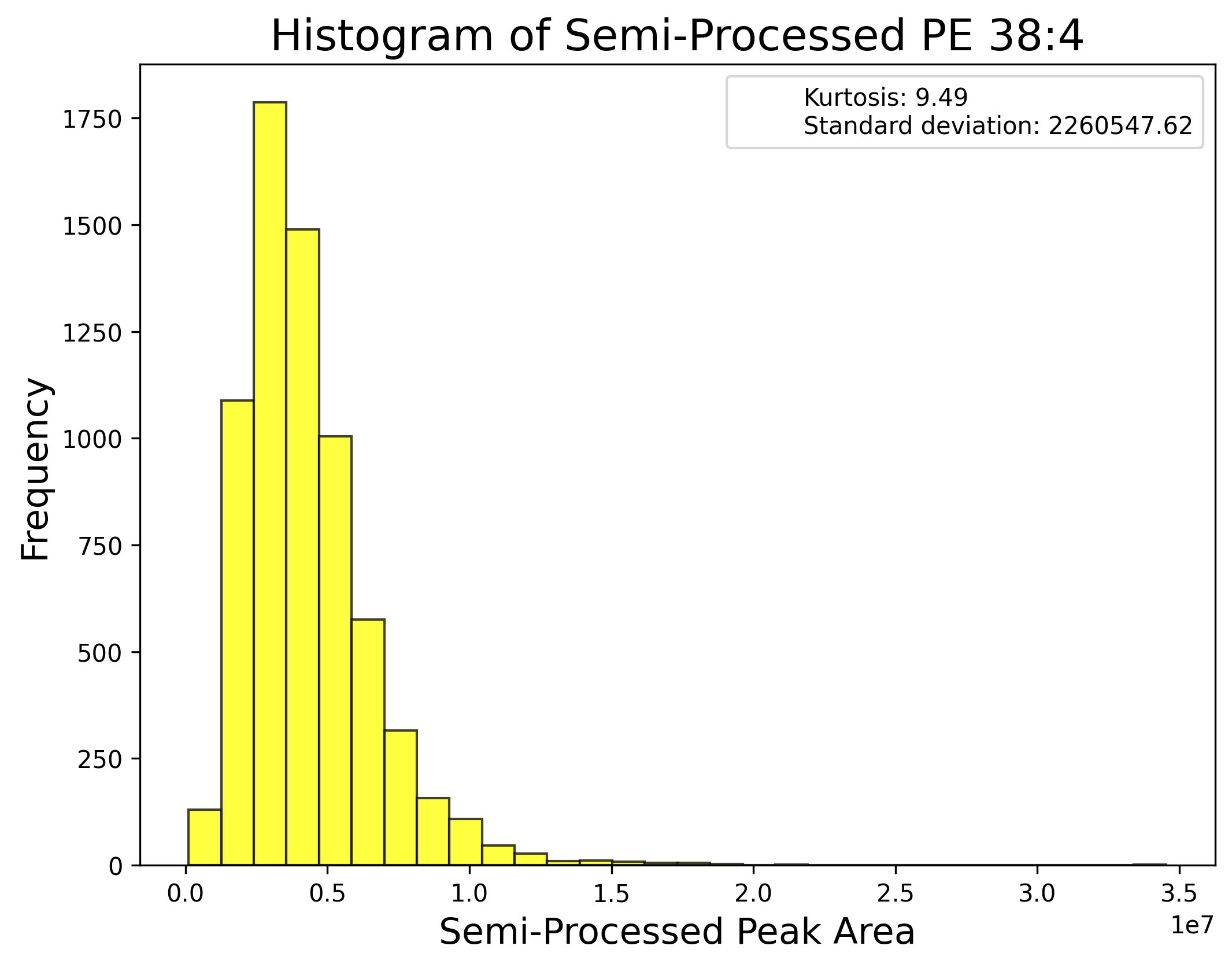

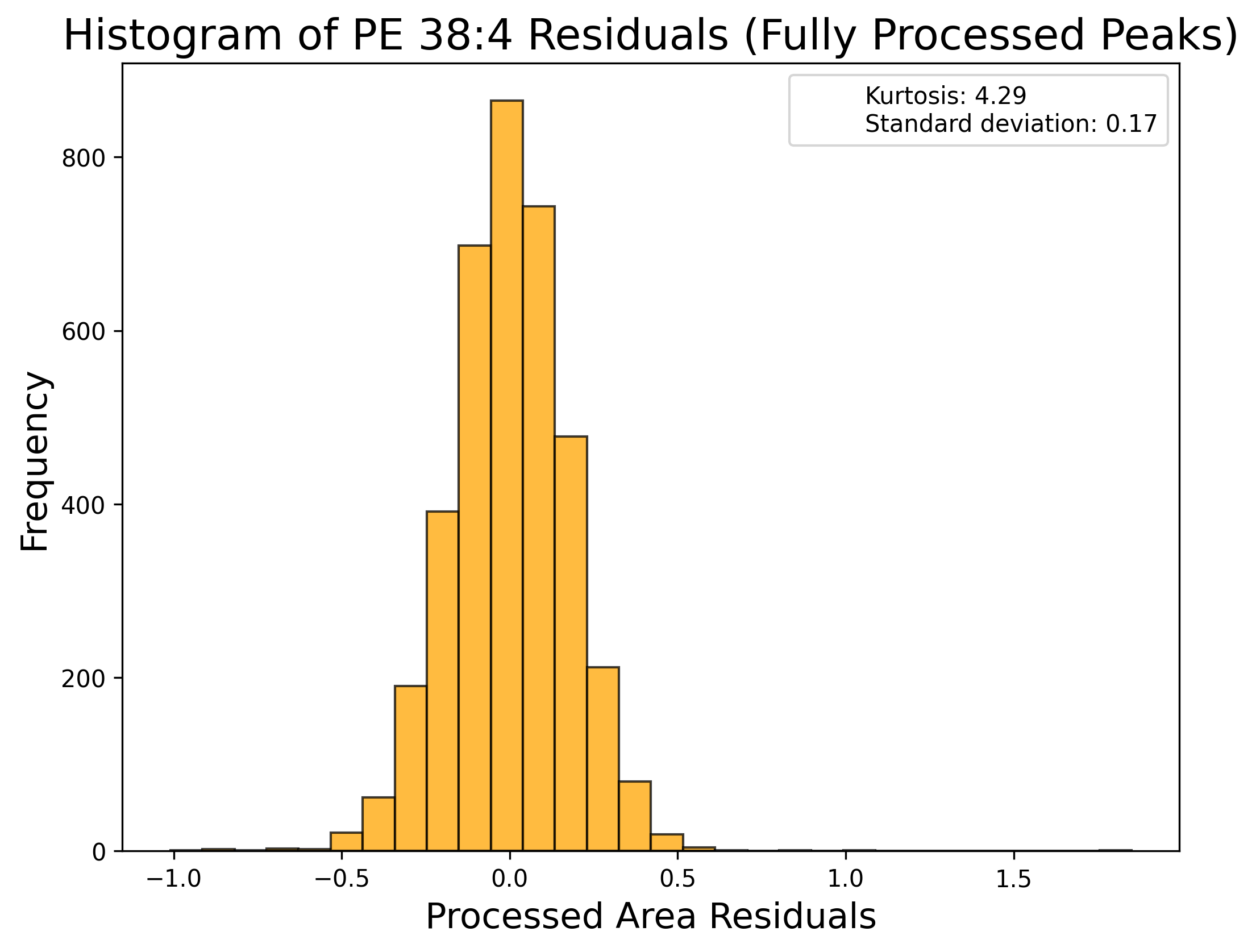

**j) Phosphatidylserine:**

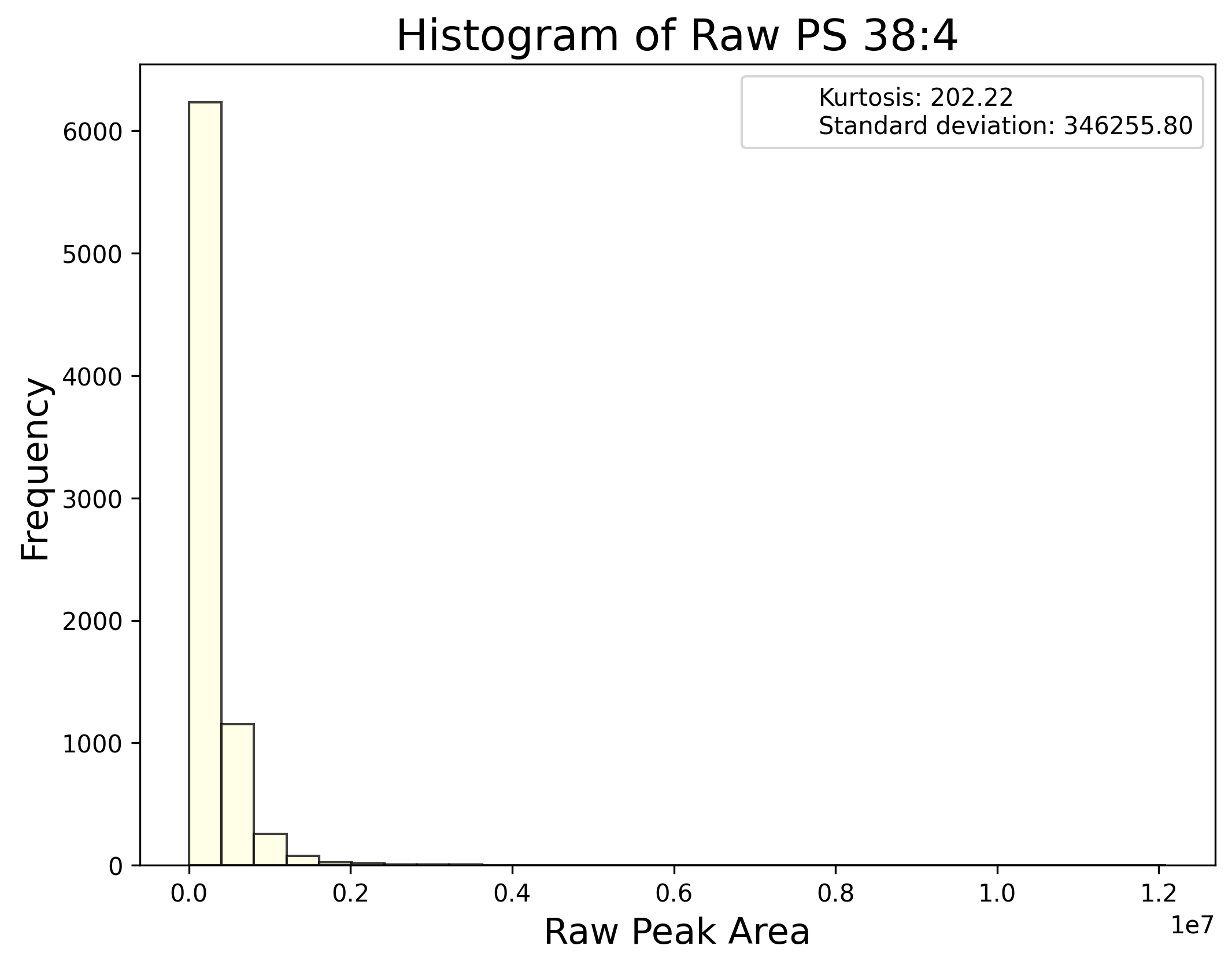

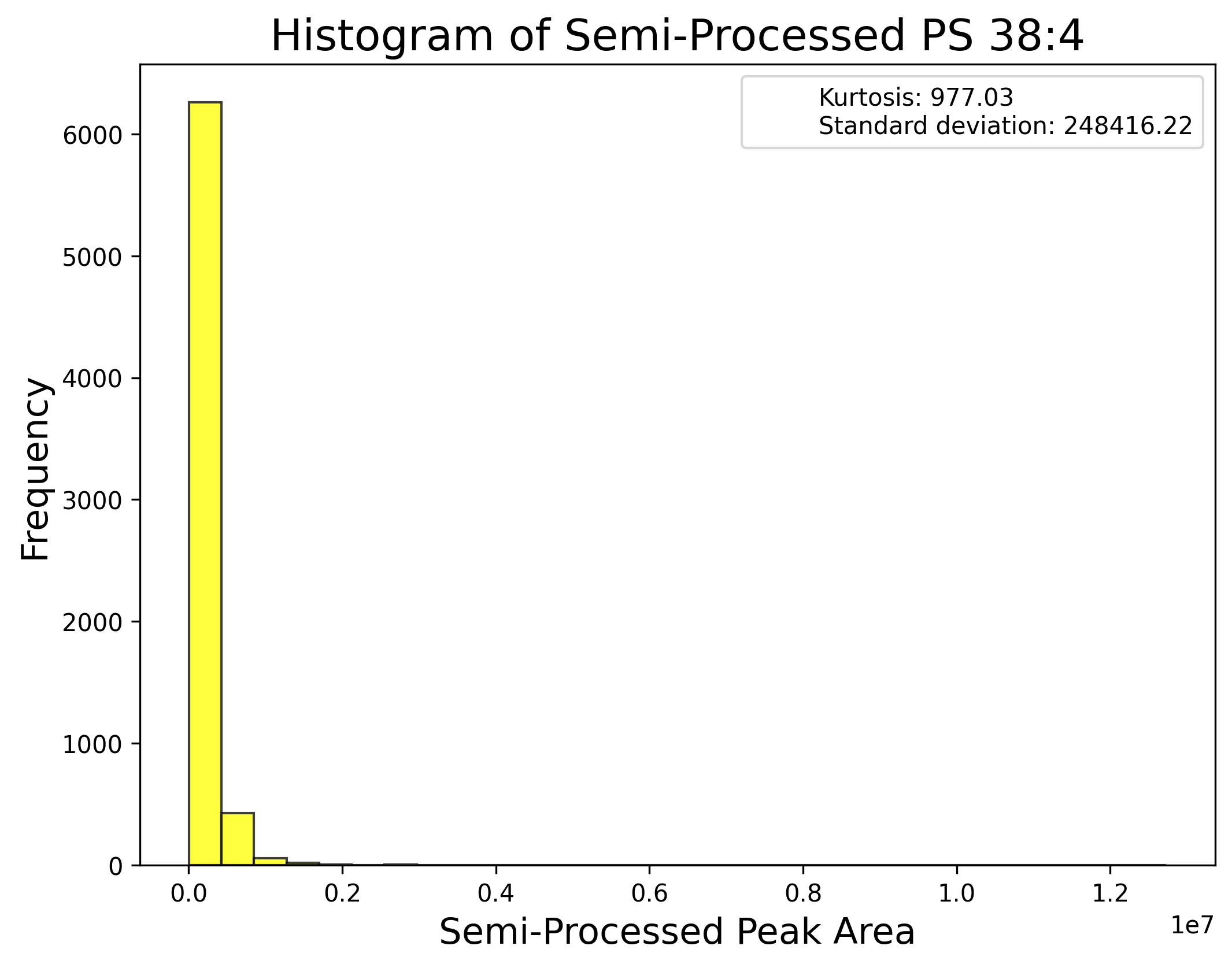

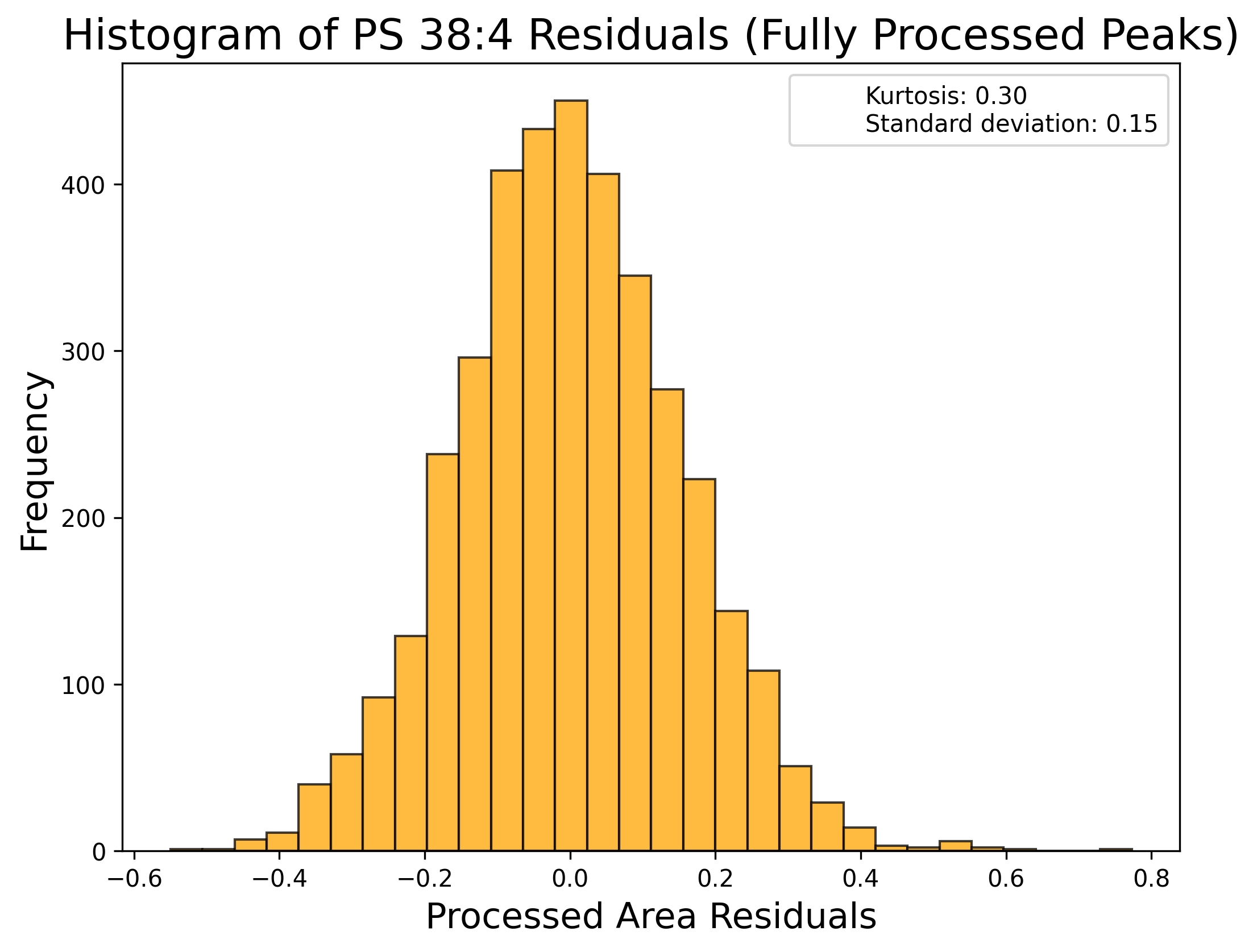

**k) Sphingomyelin:**

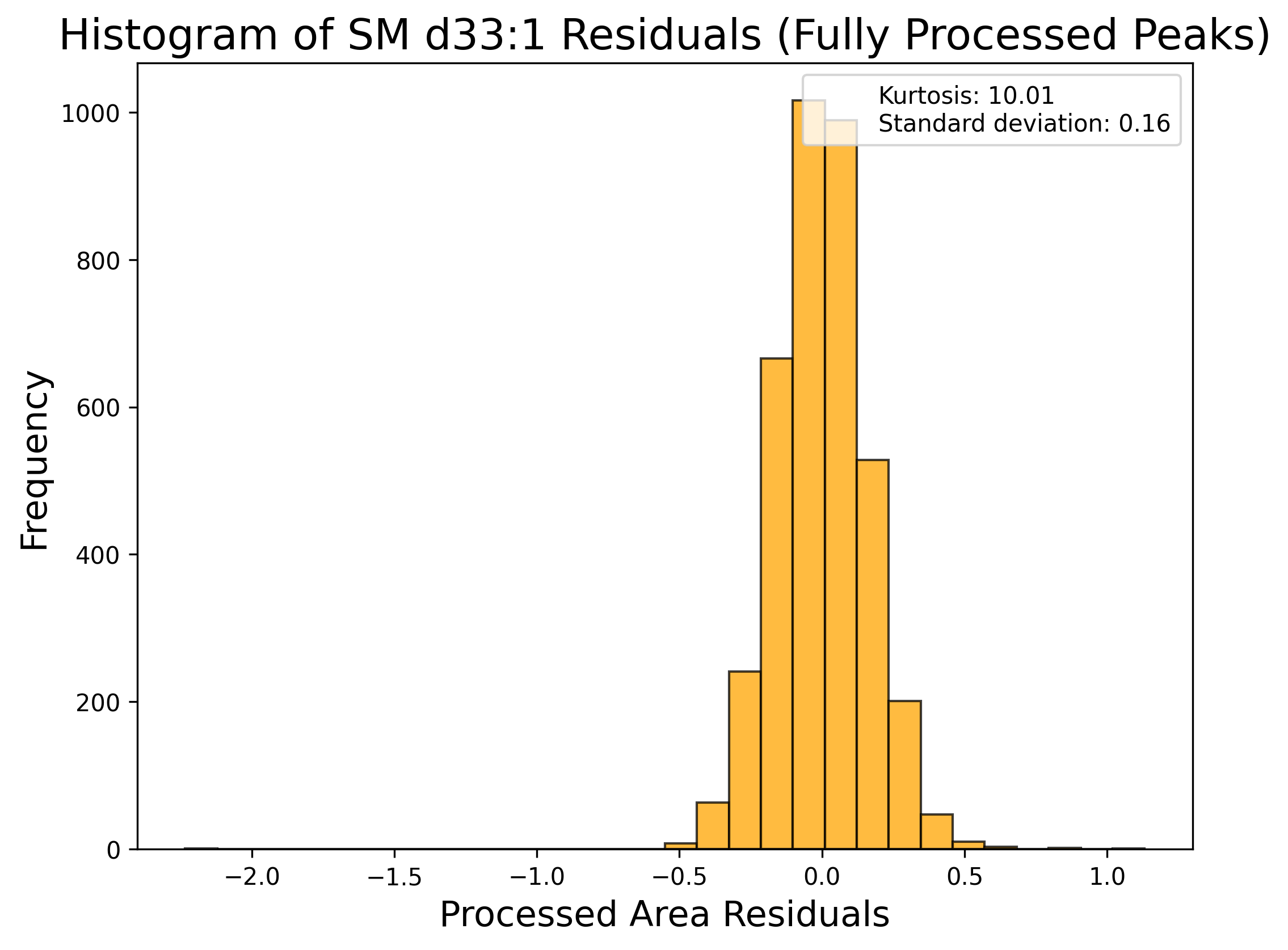

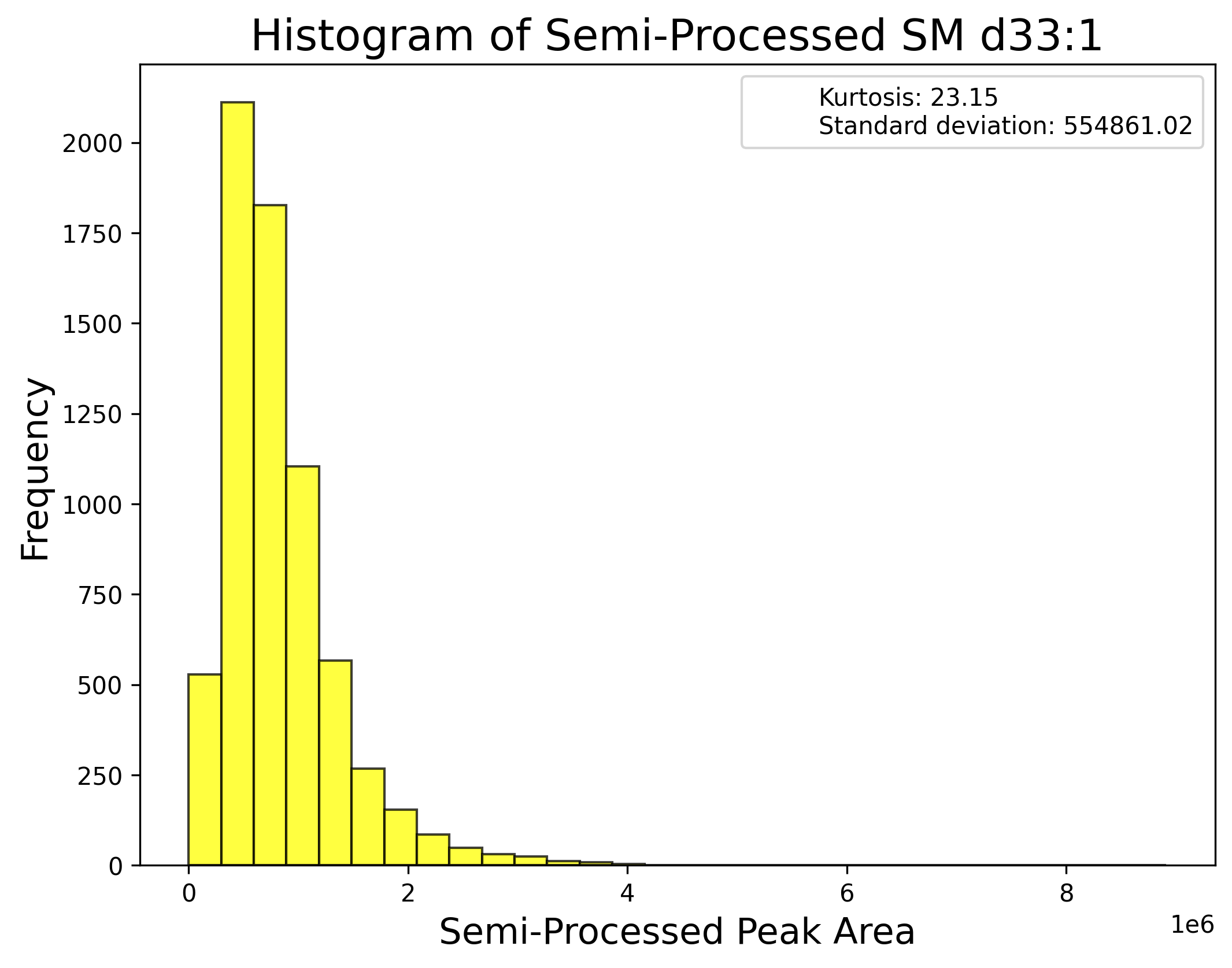

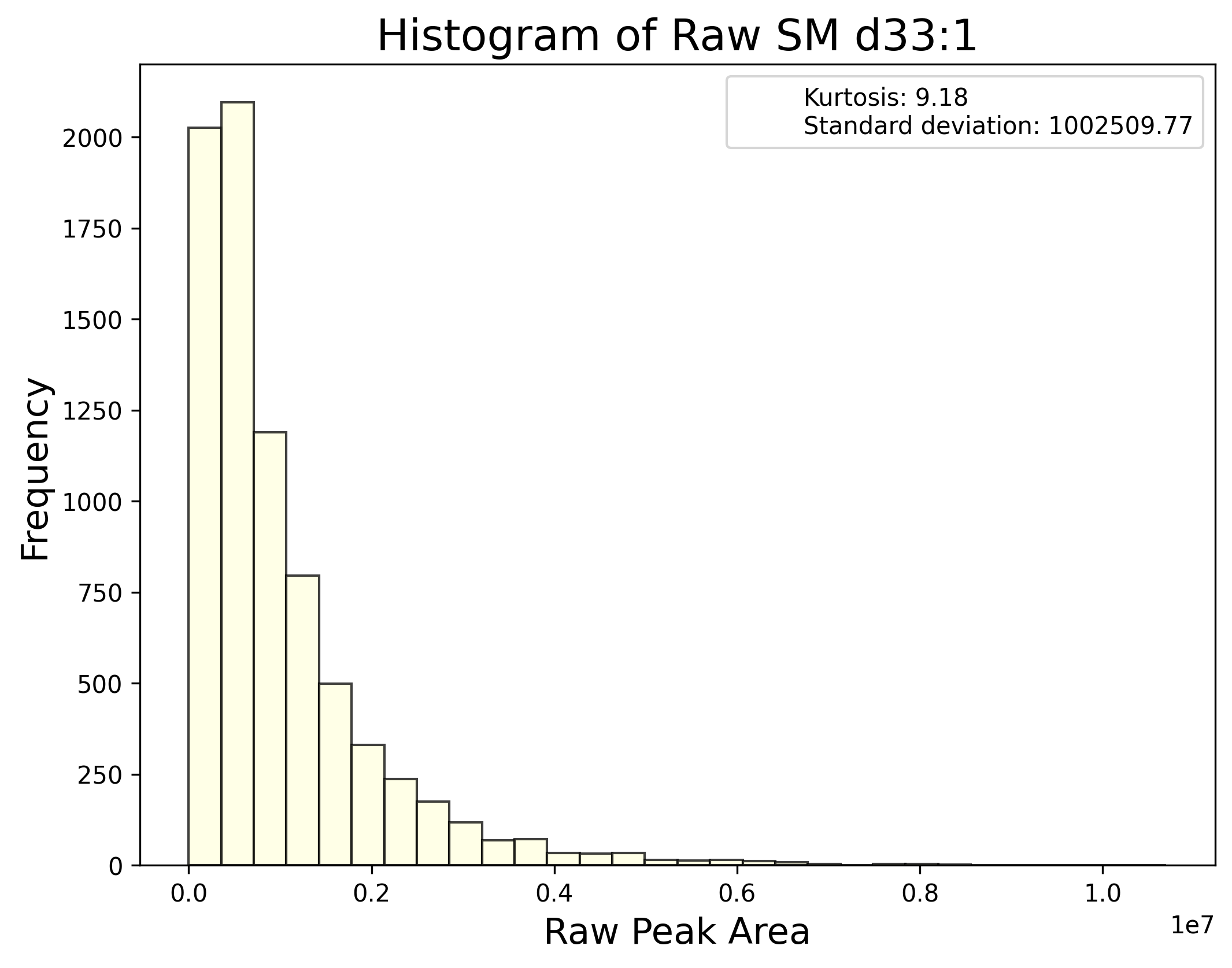

**l) Triacylglycerol**:

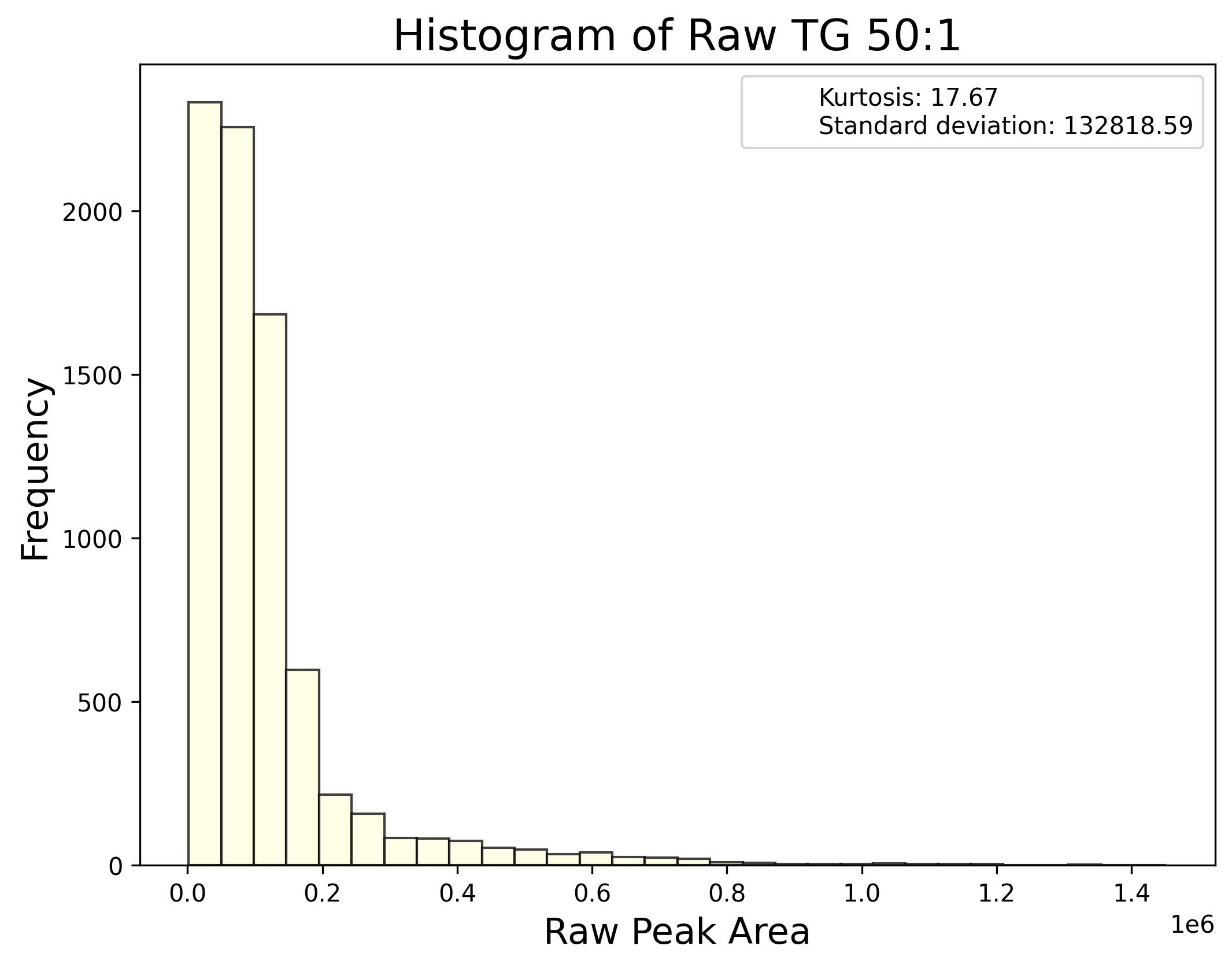

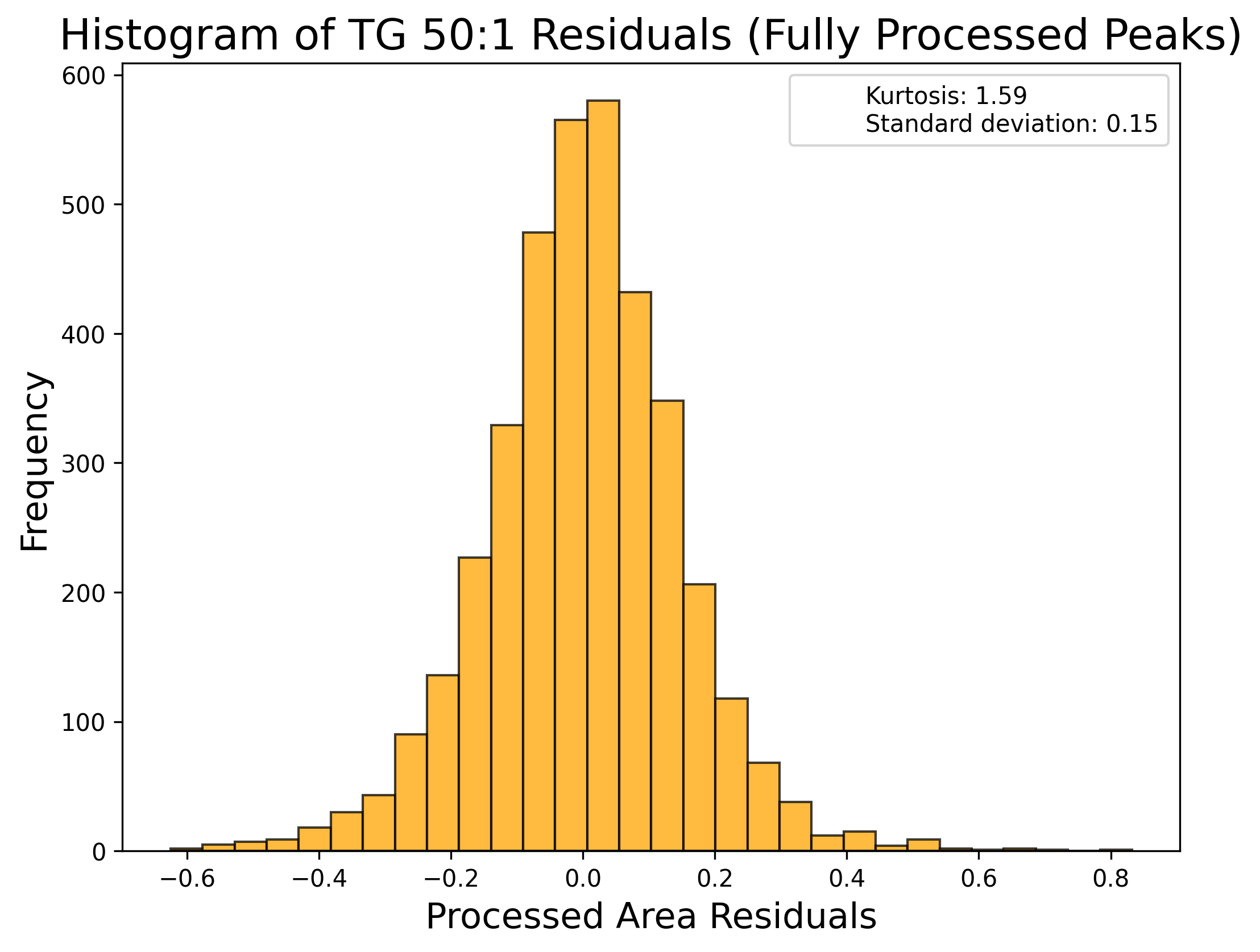

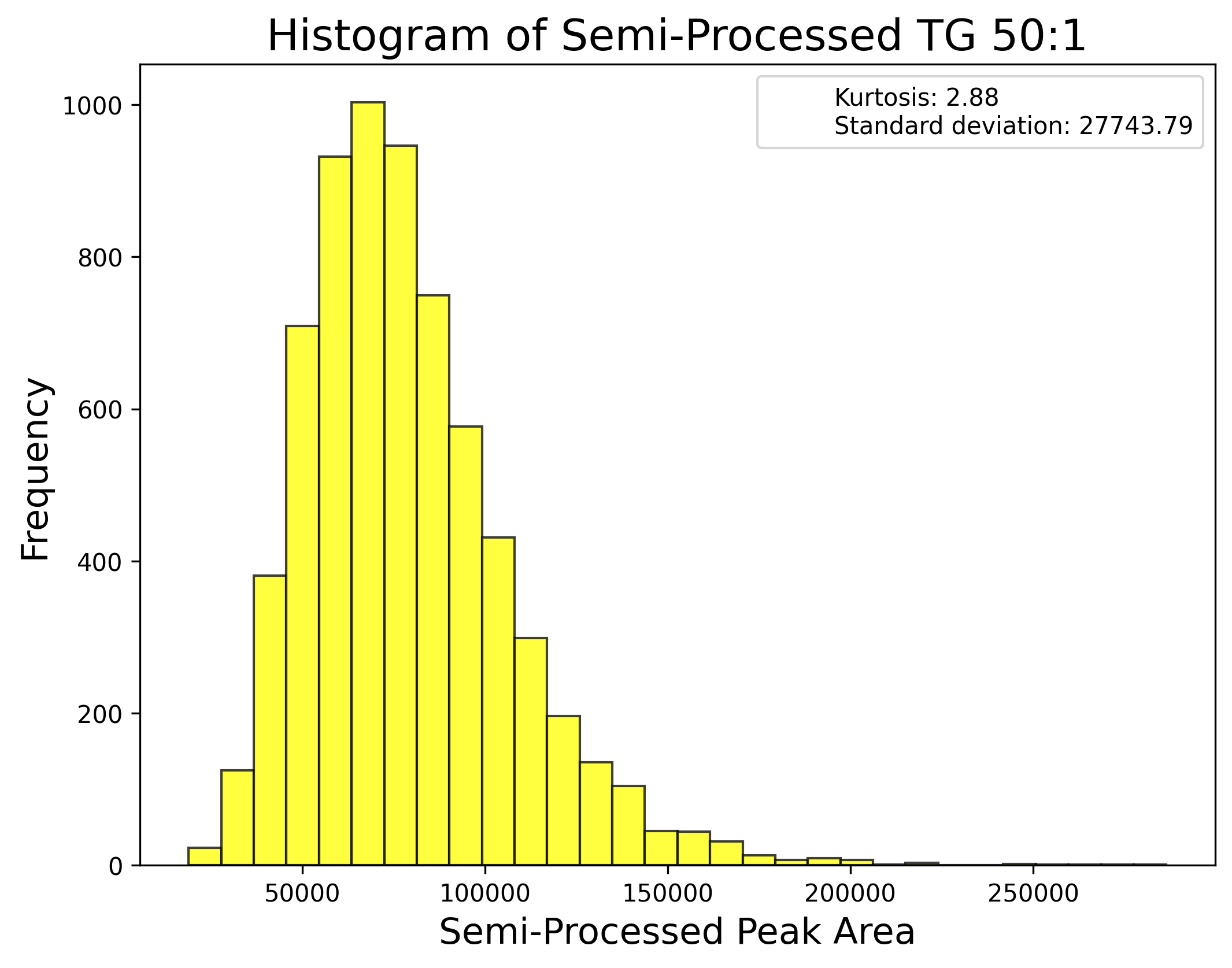

**m) Polar primary source: Diet**

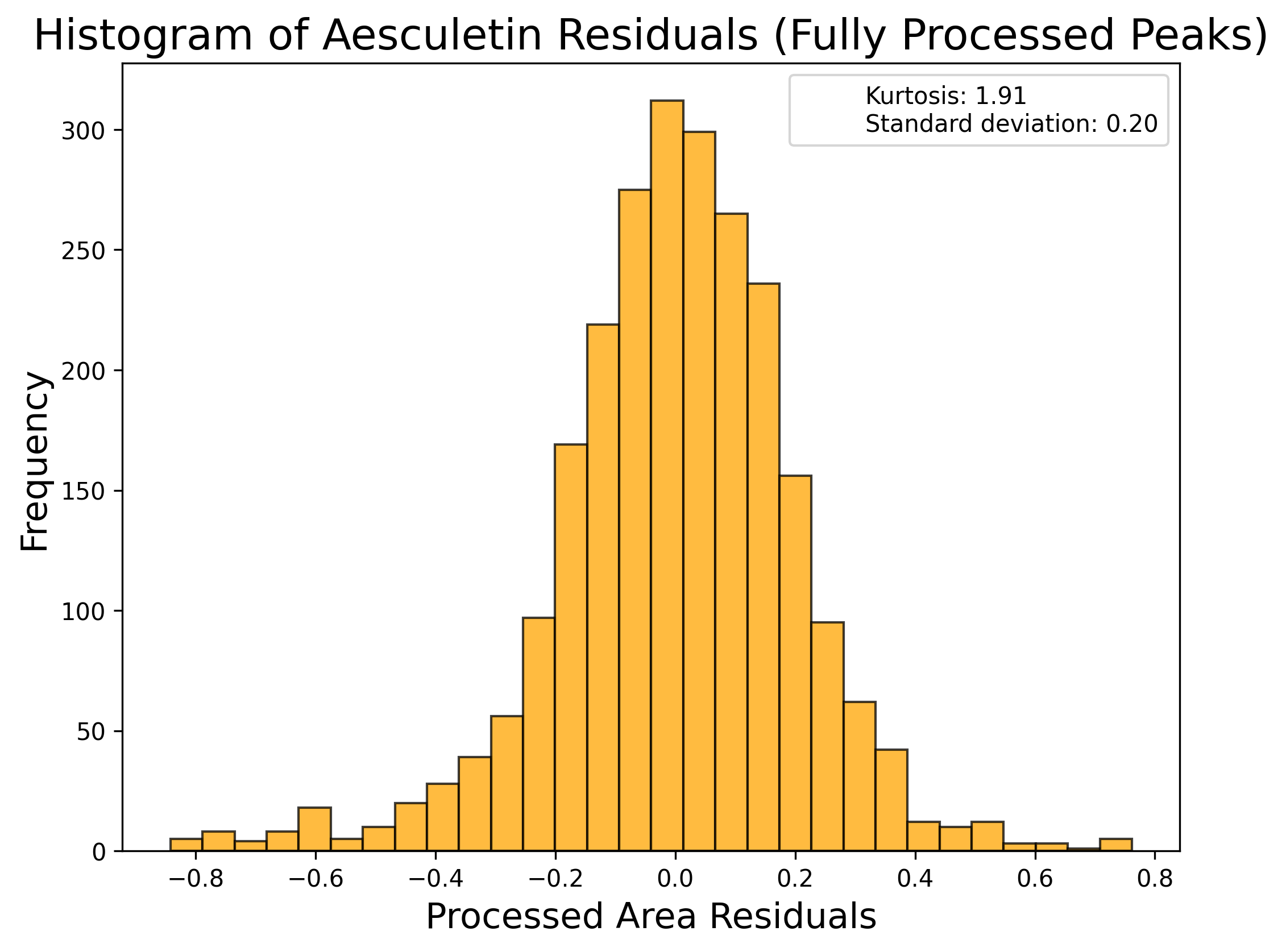

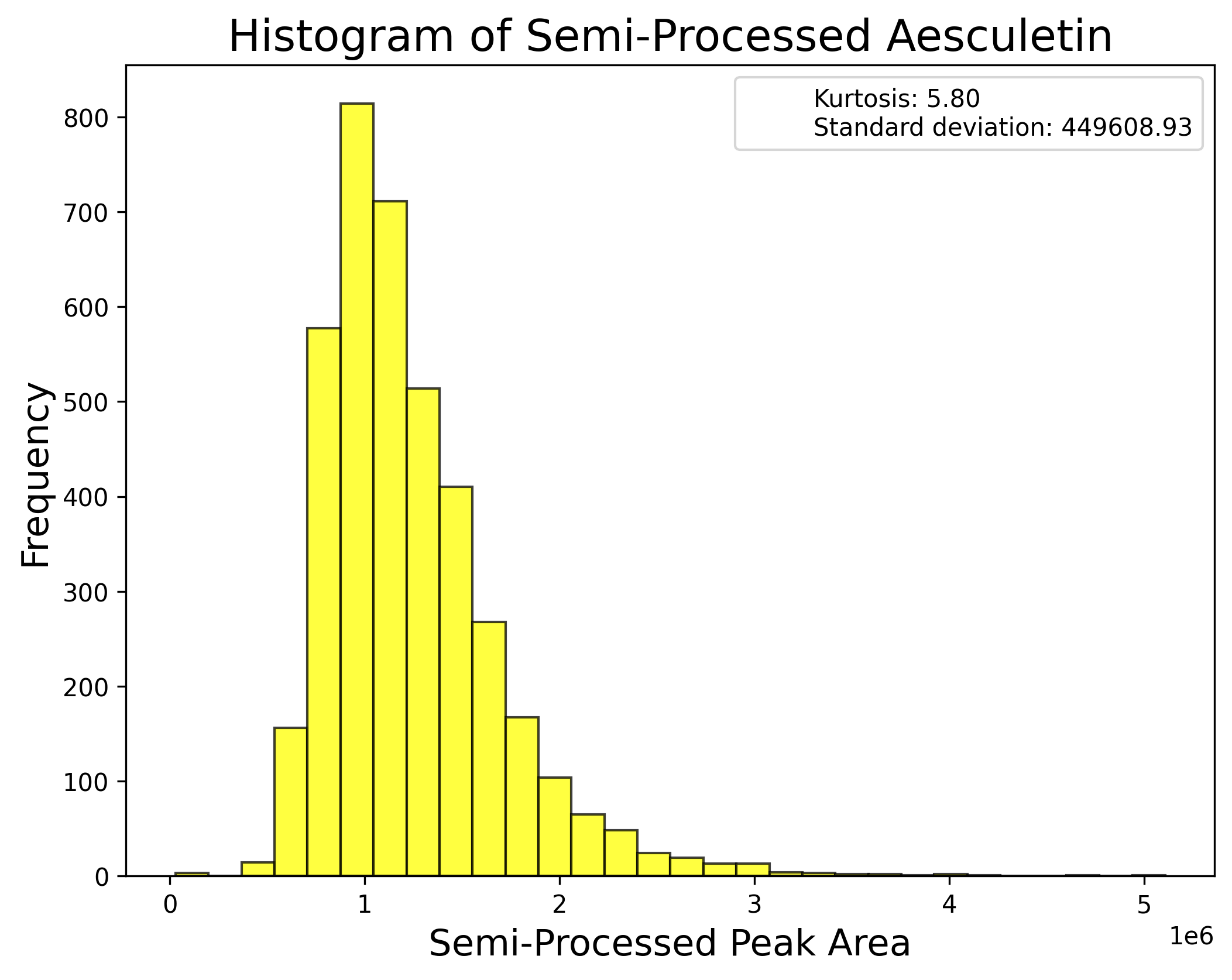

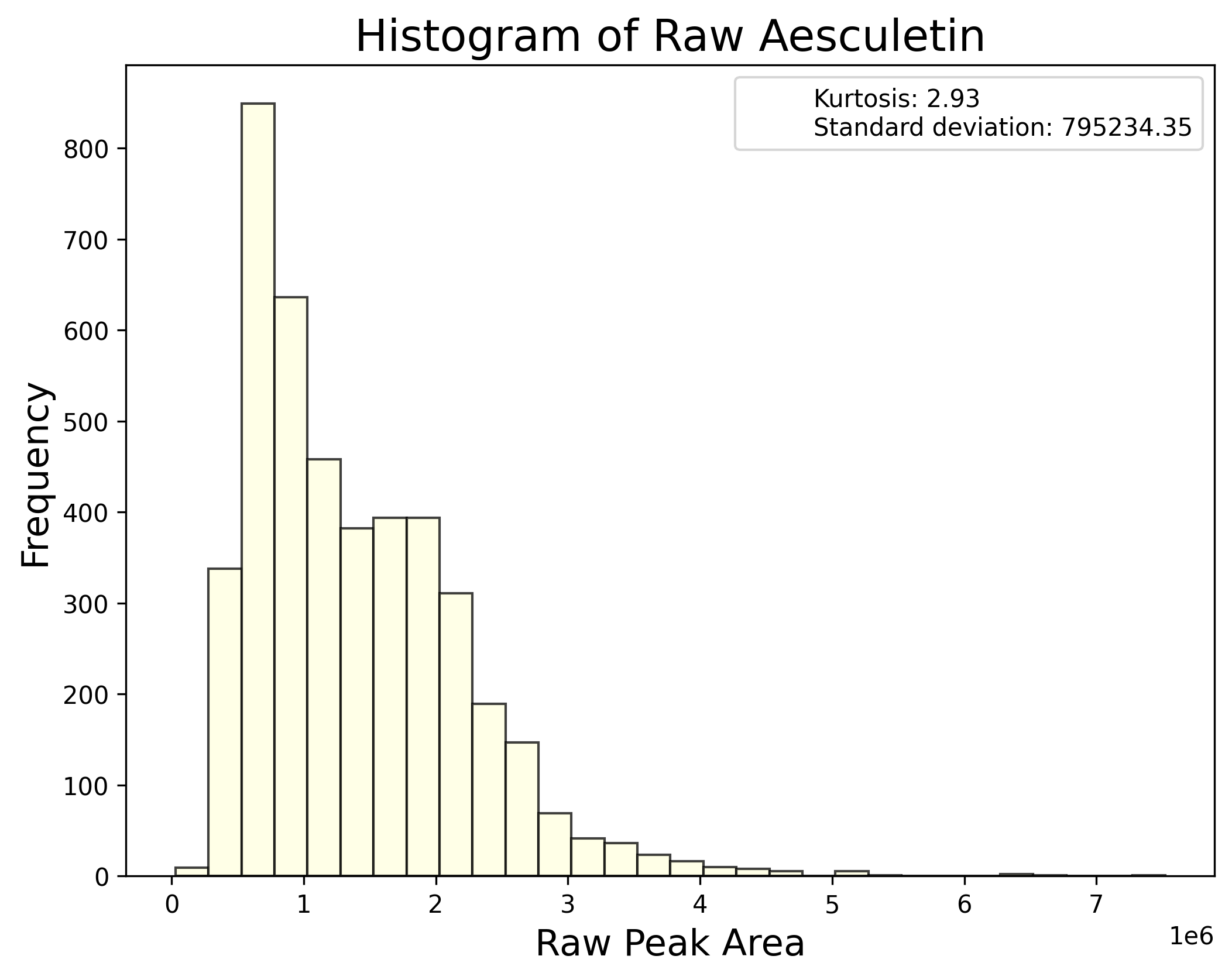

**n) Polar primary source: Diet / gut microbiome**

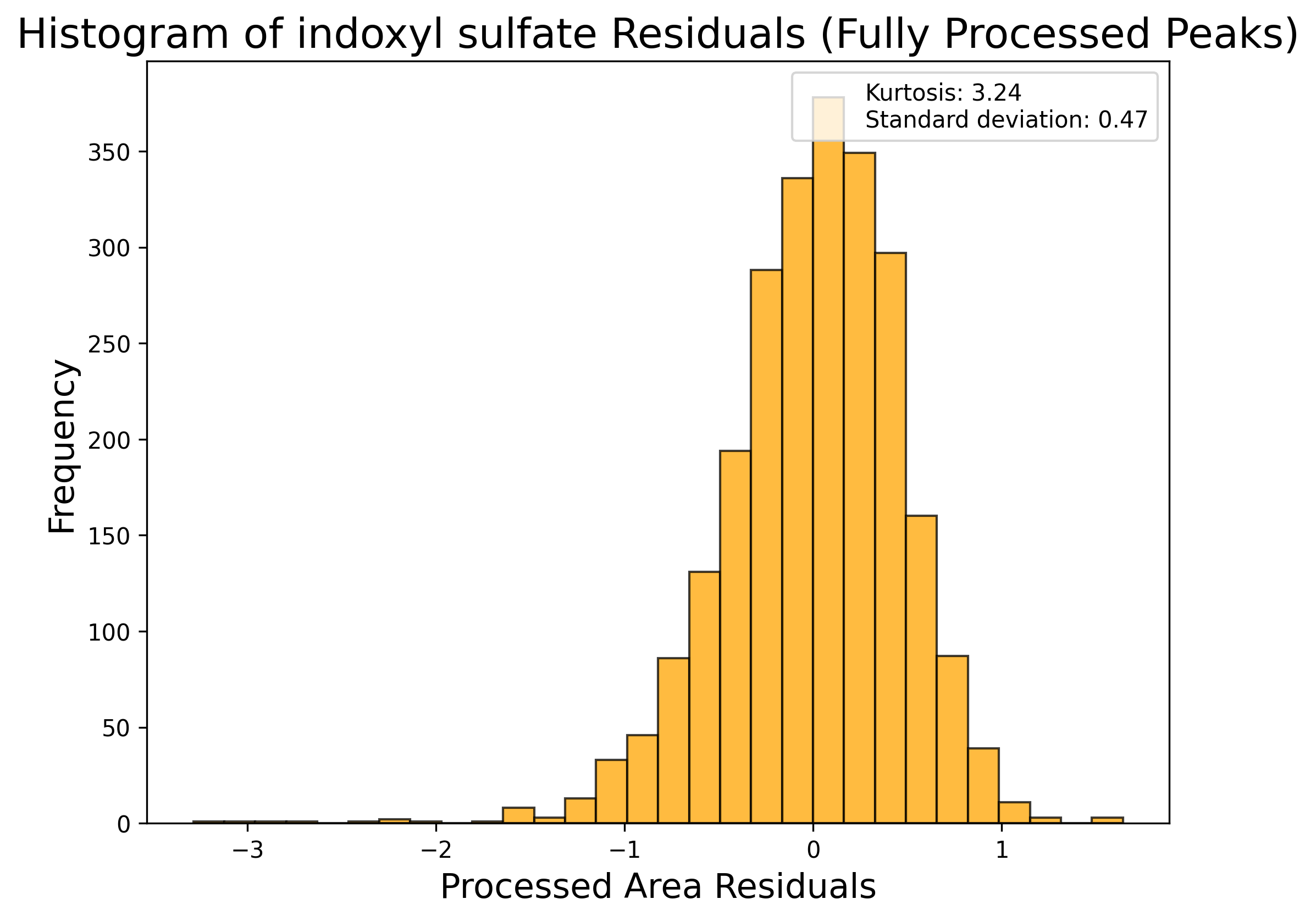

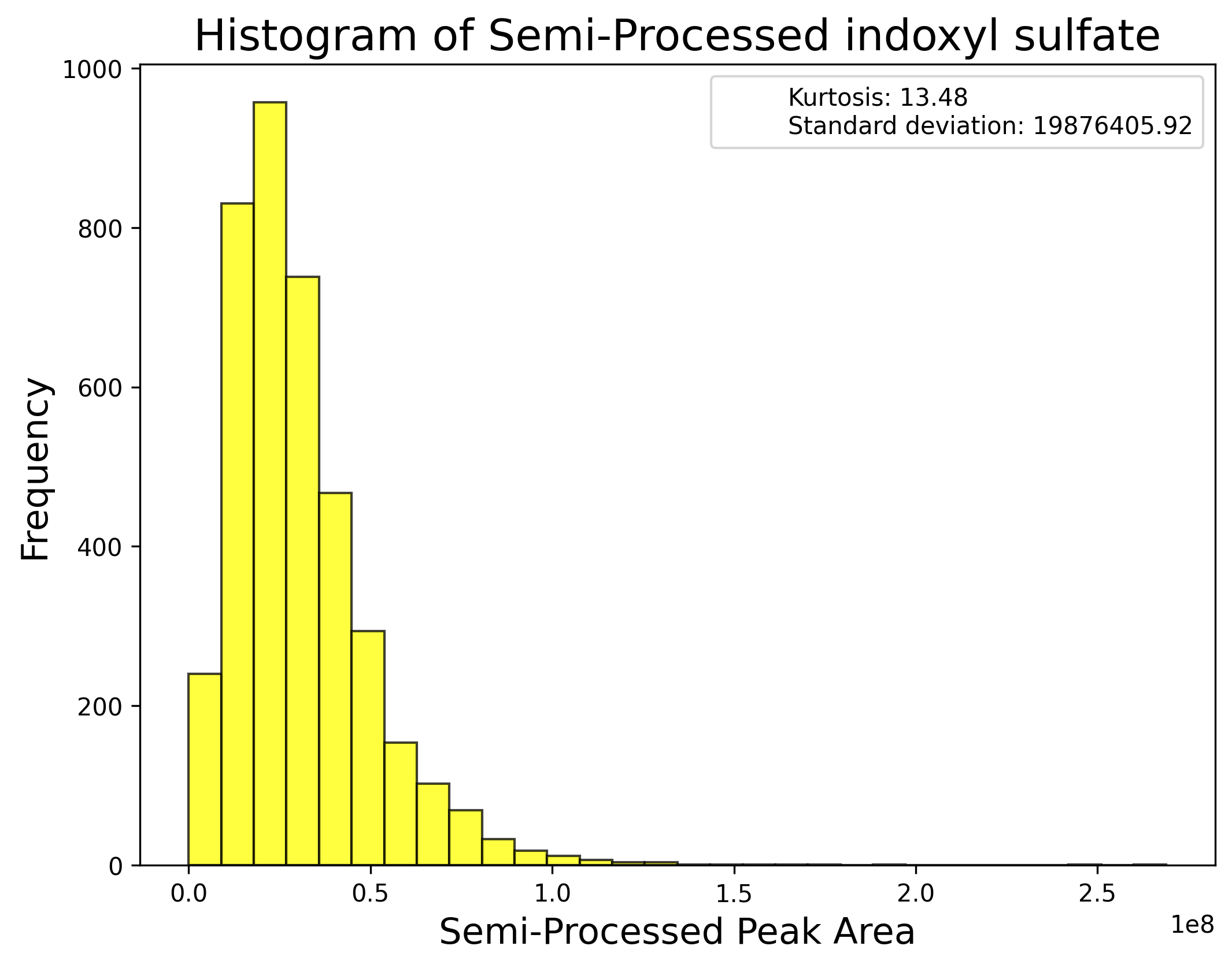

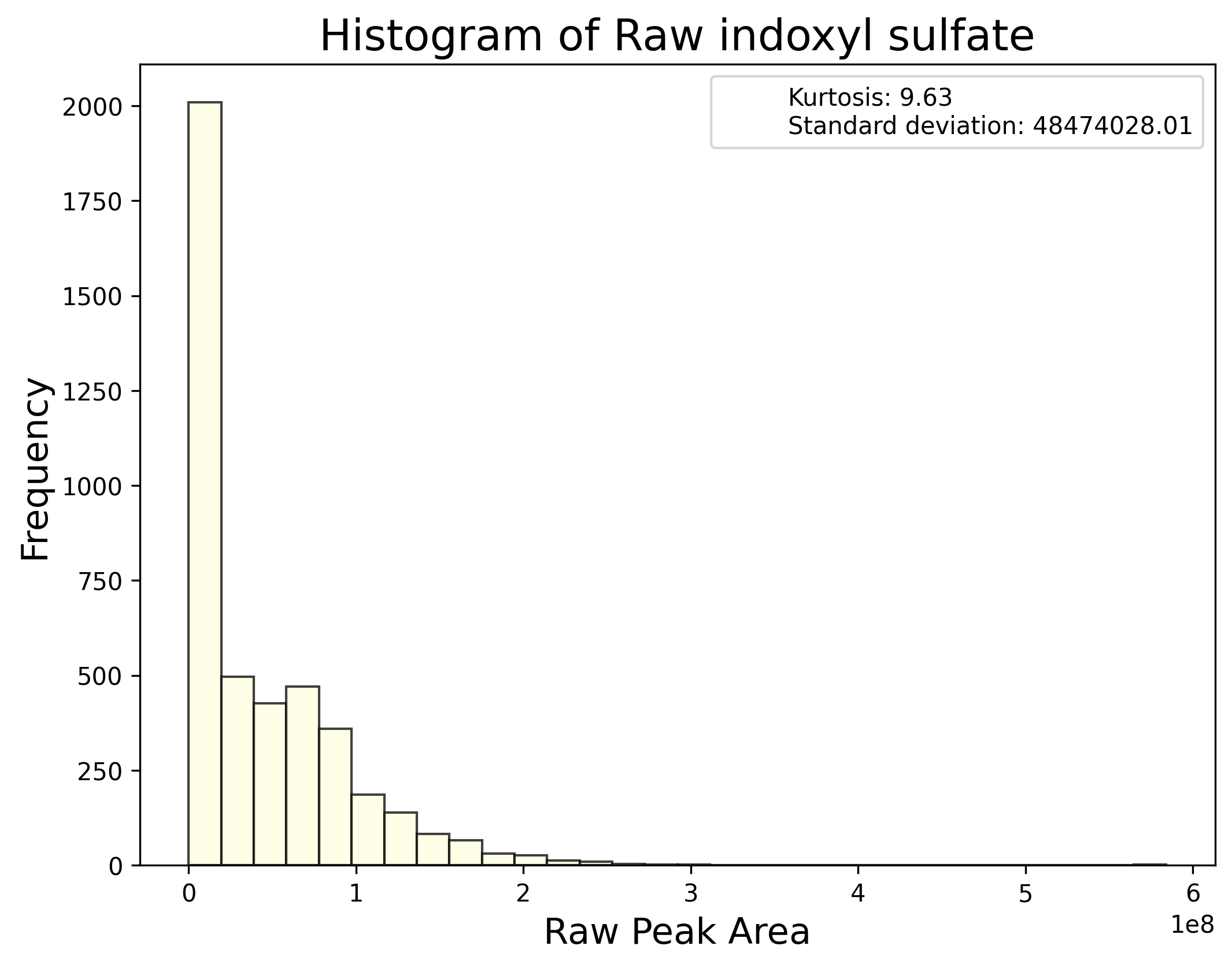

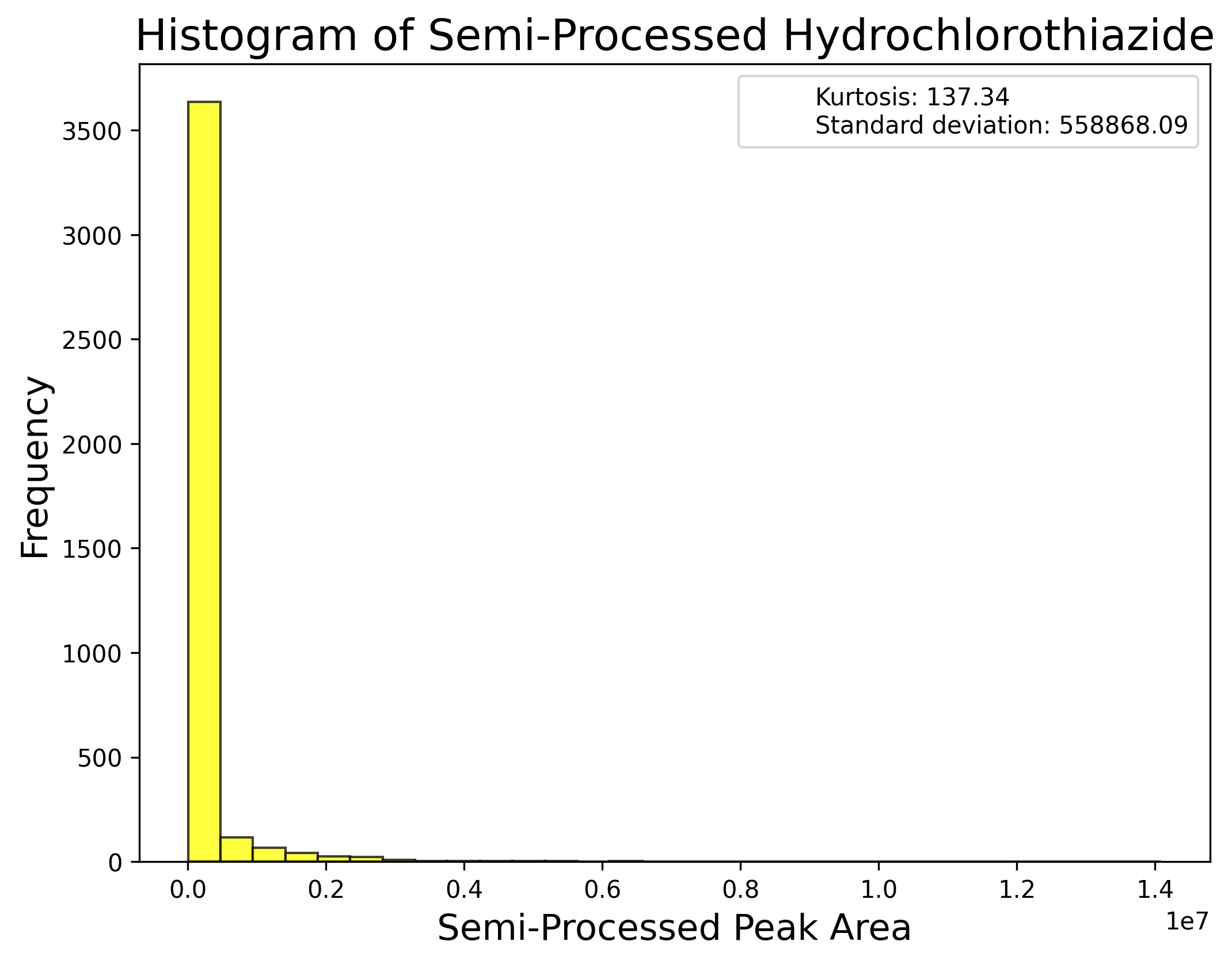
**o)** **Polar primary source: Drugs**

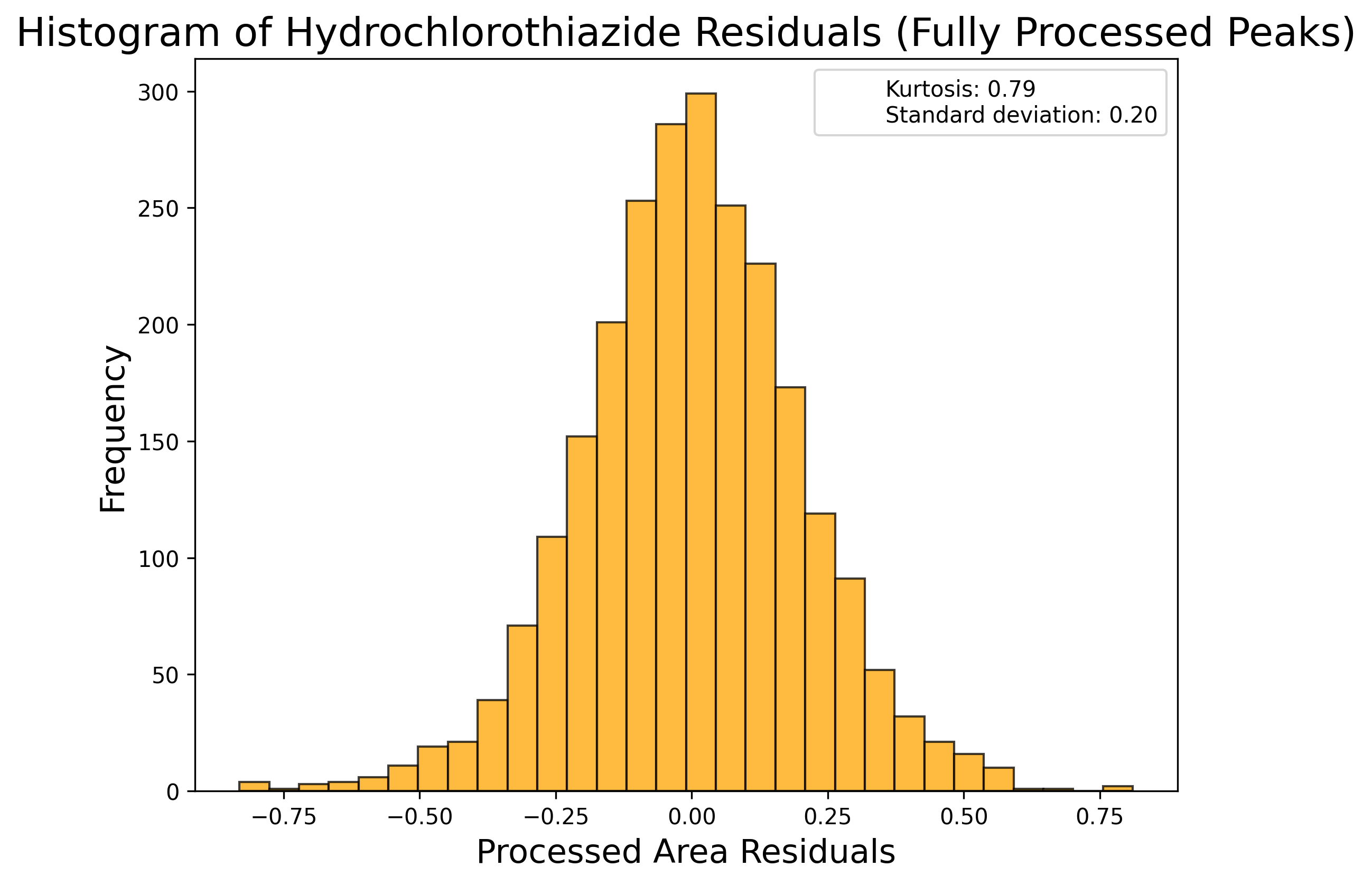

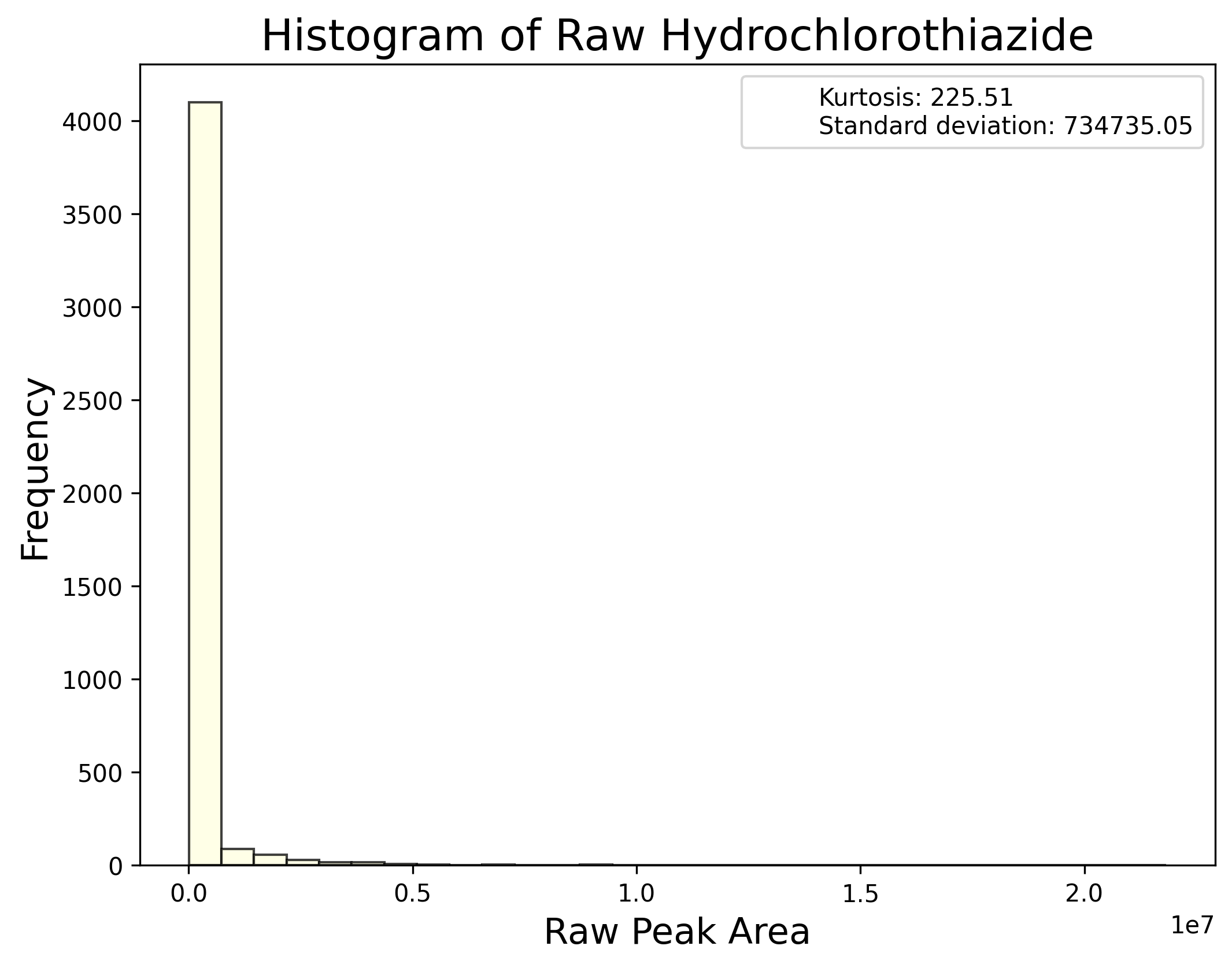

**p) Polar primary source: Endogenous**

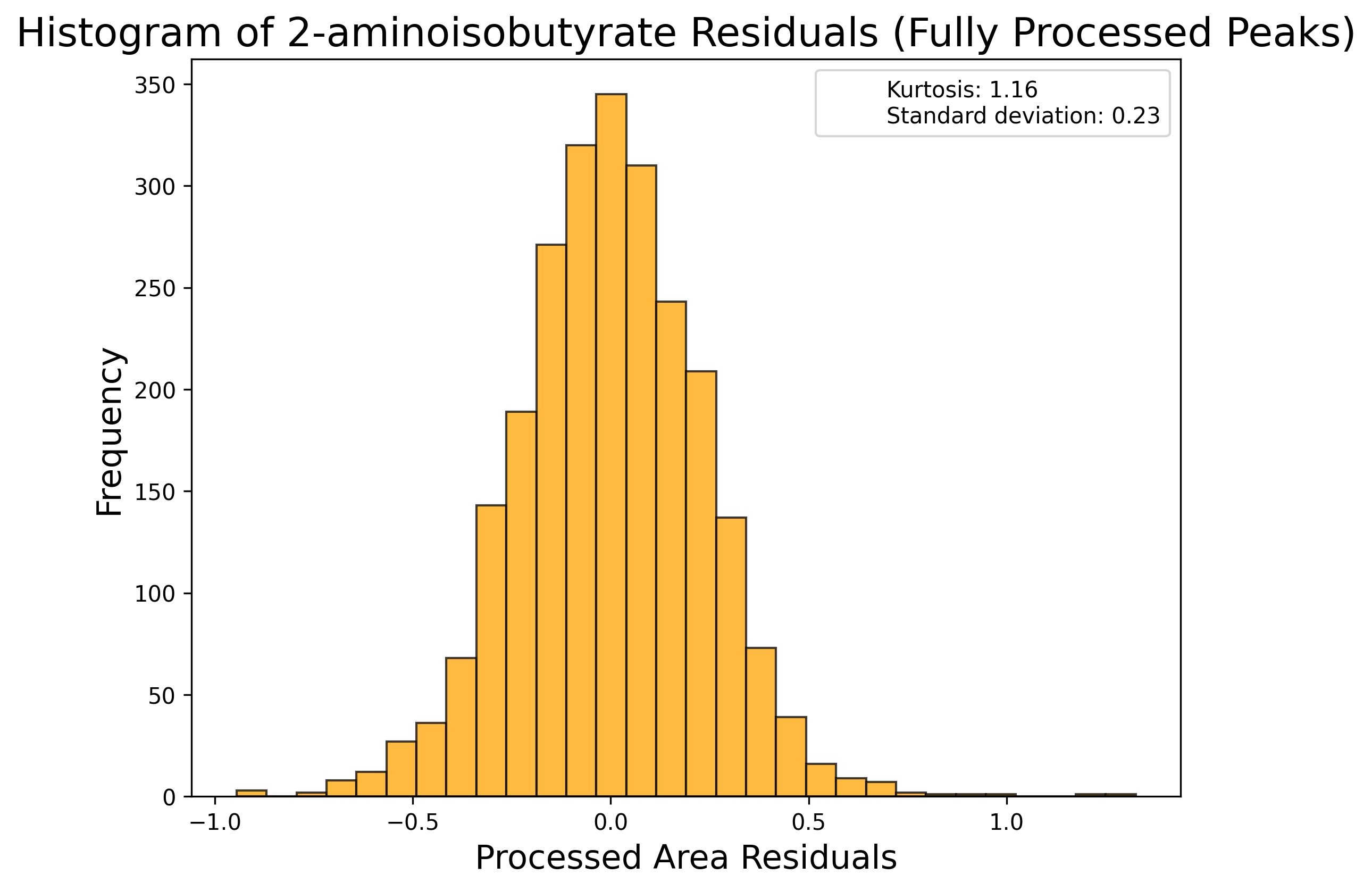

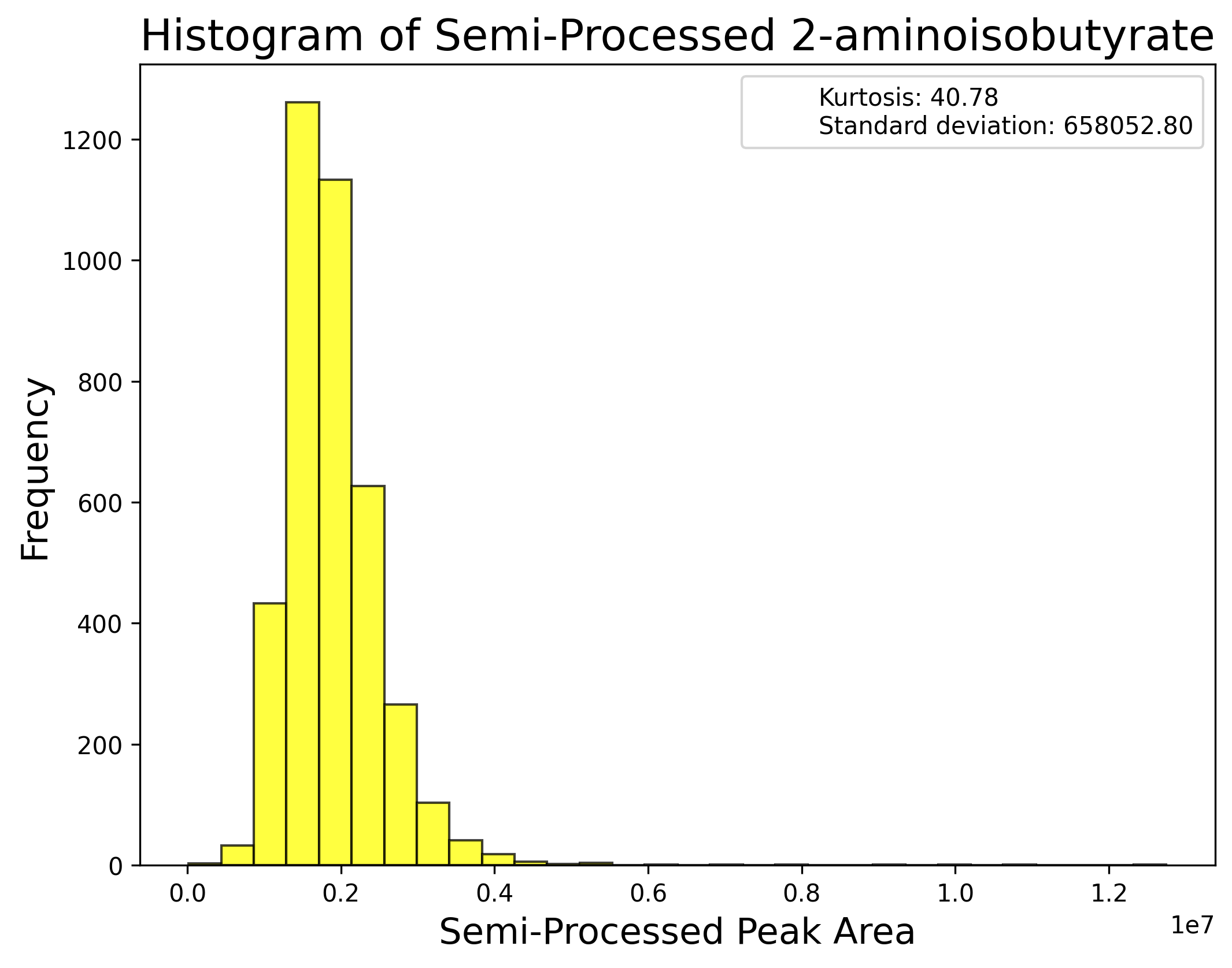

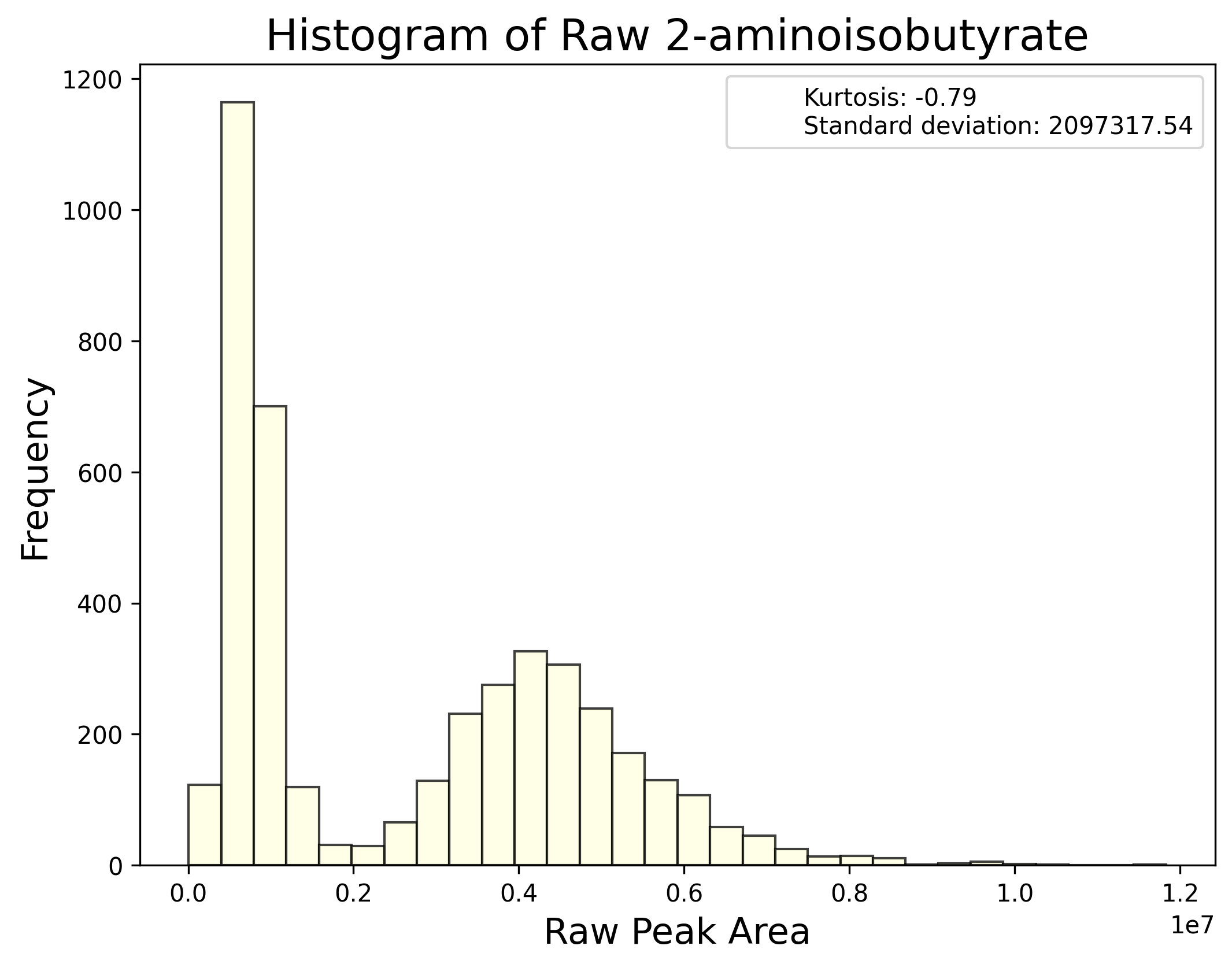

**q) Polar primary source: Exposures**

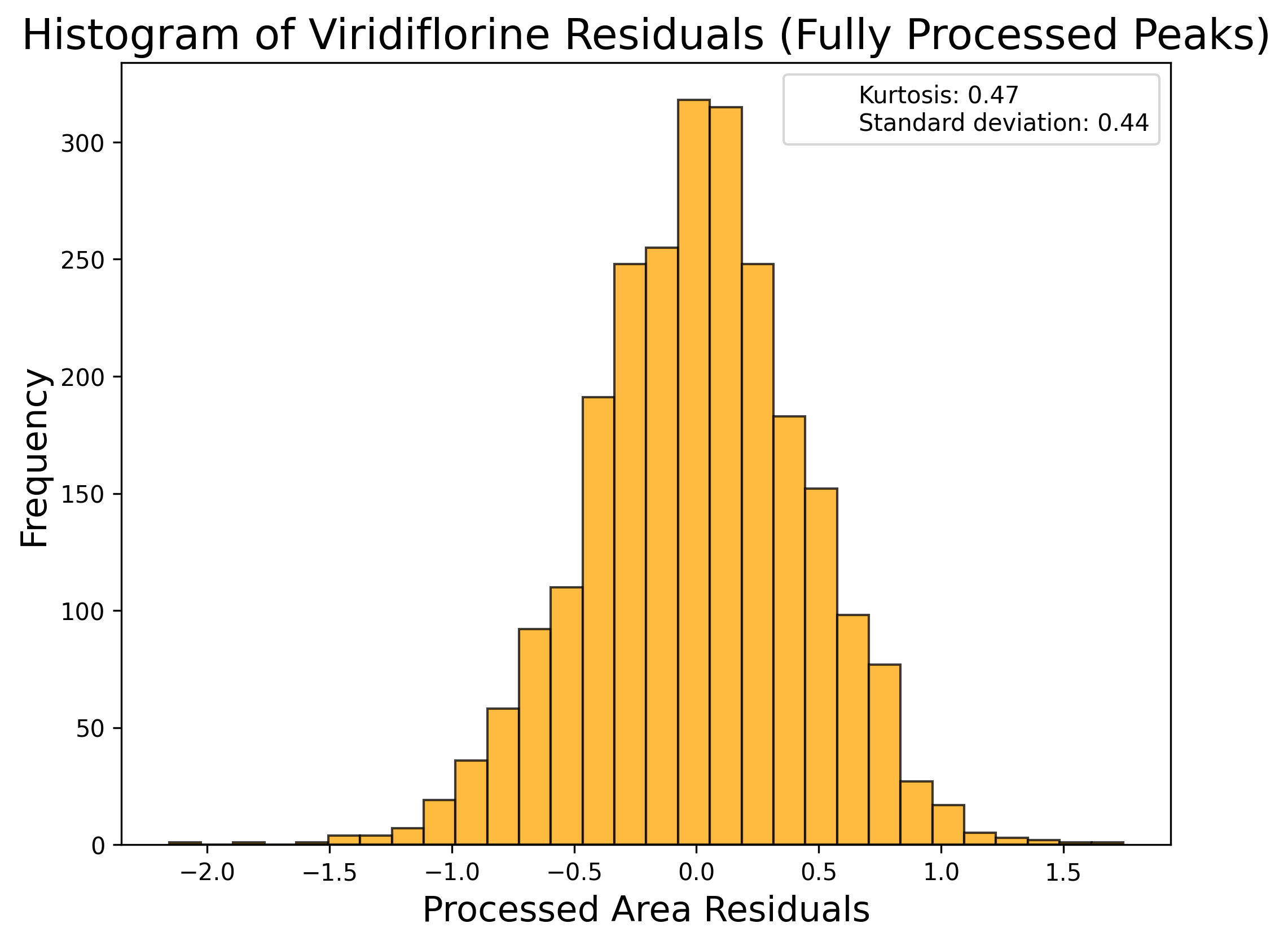

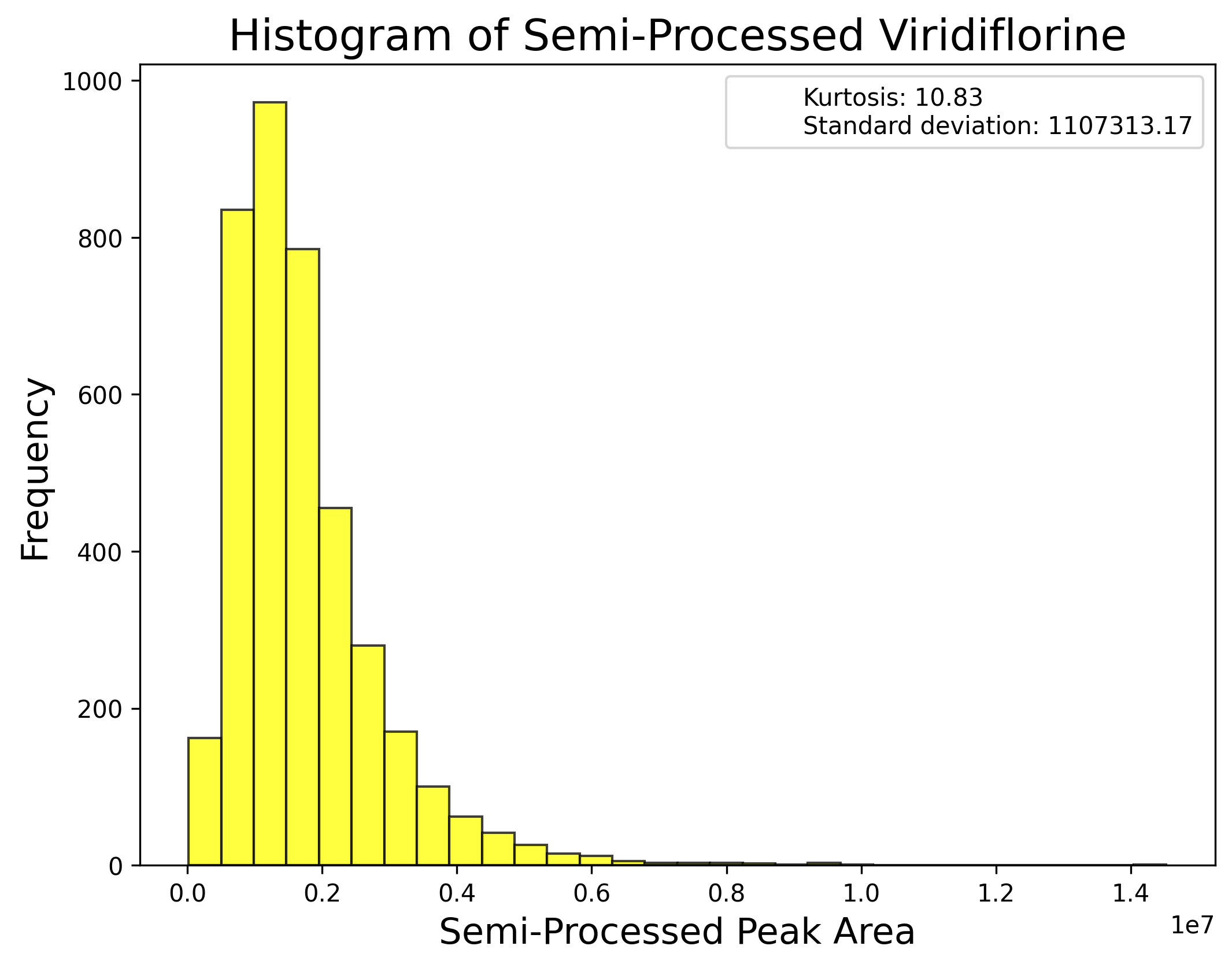

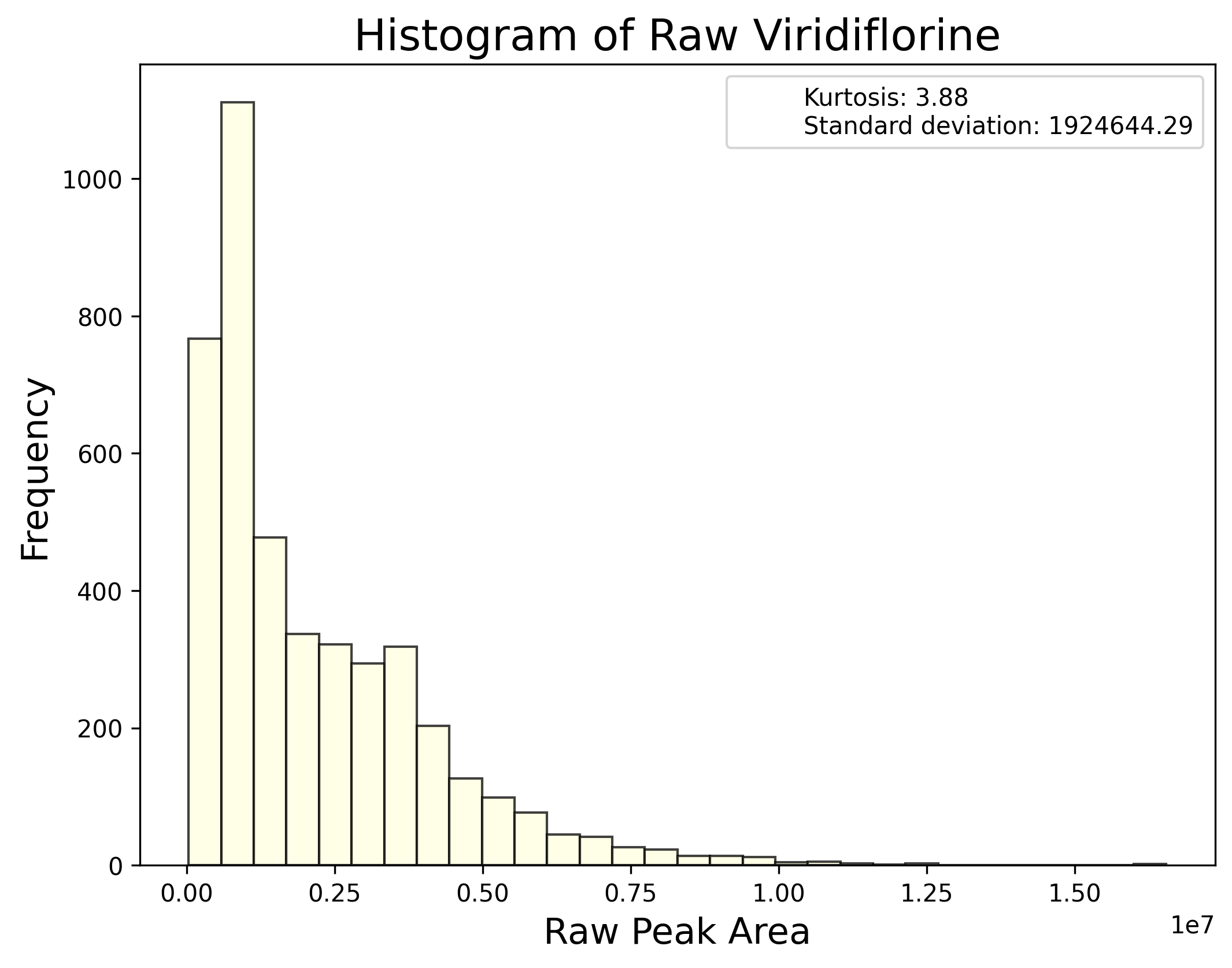

**r) Polar primary source: Mixed**

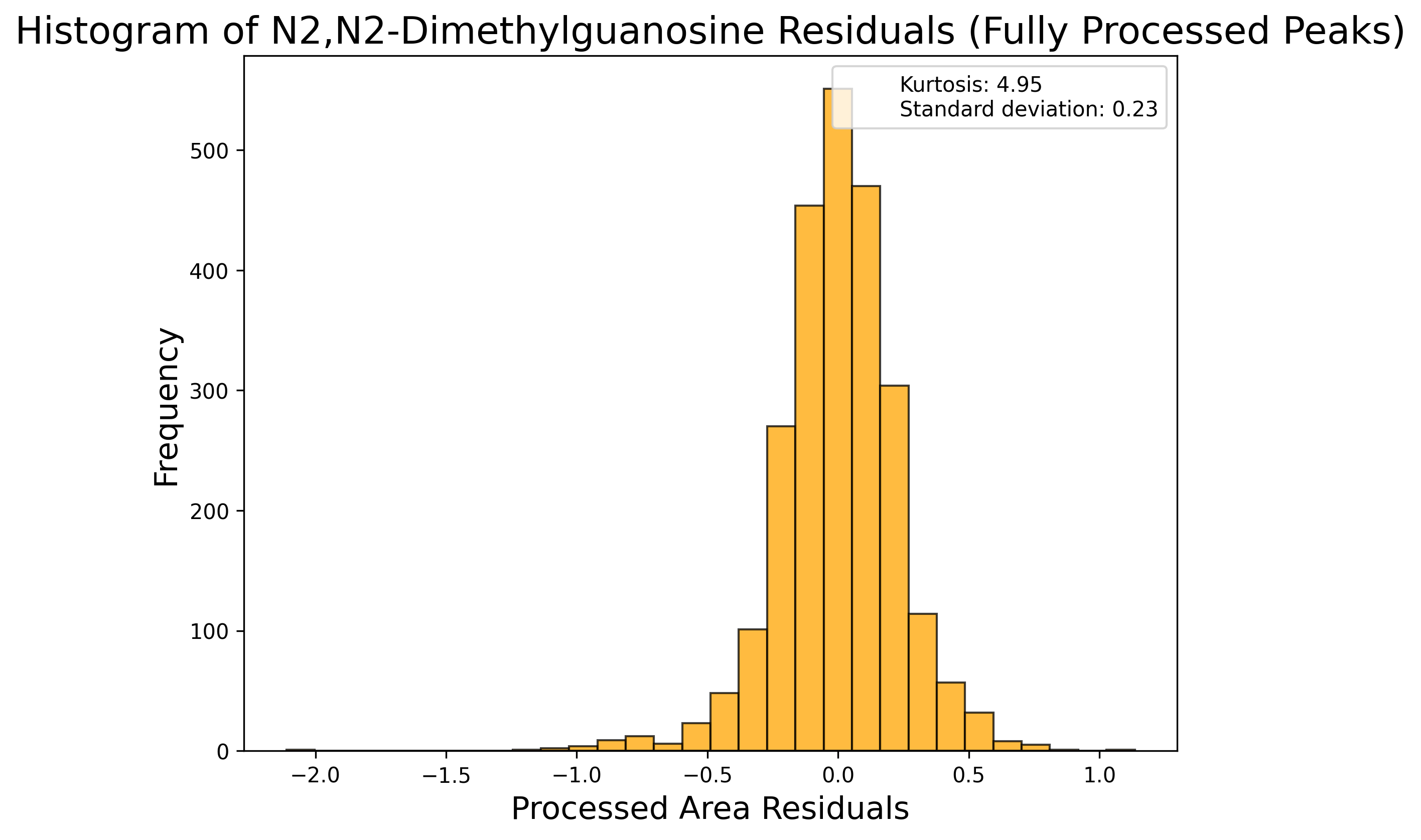

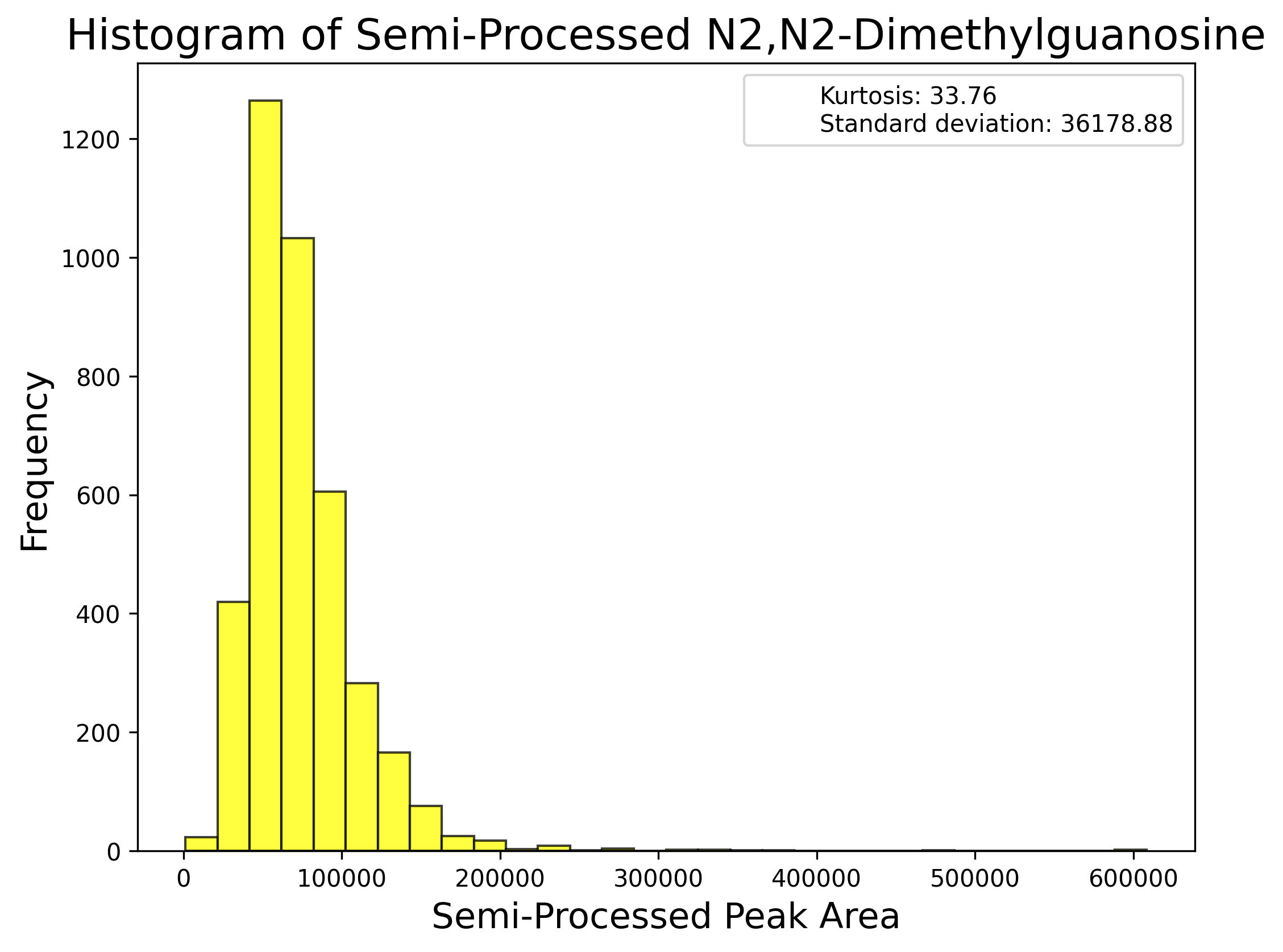

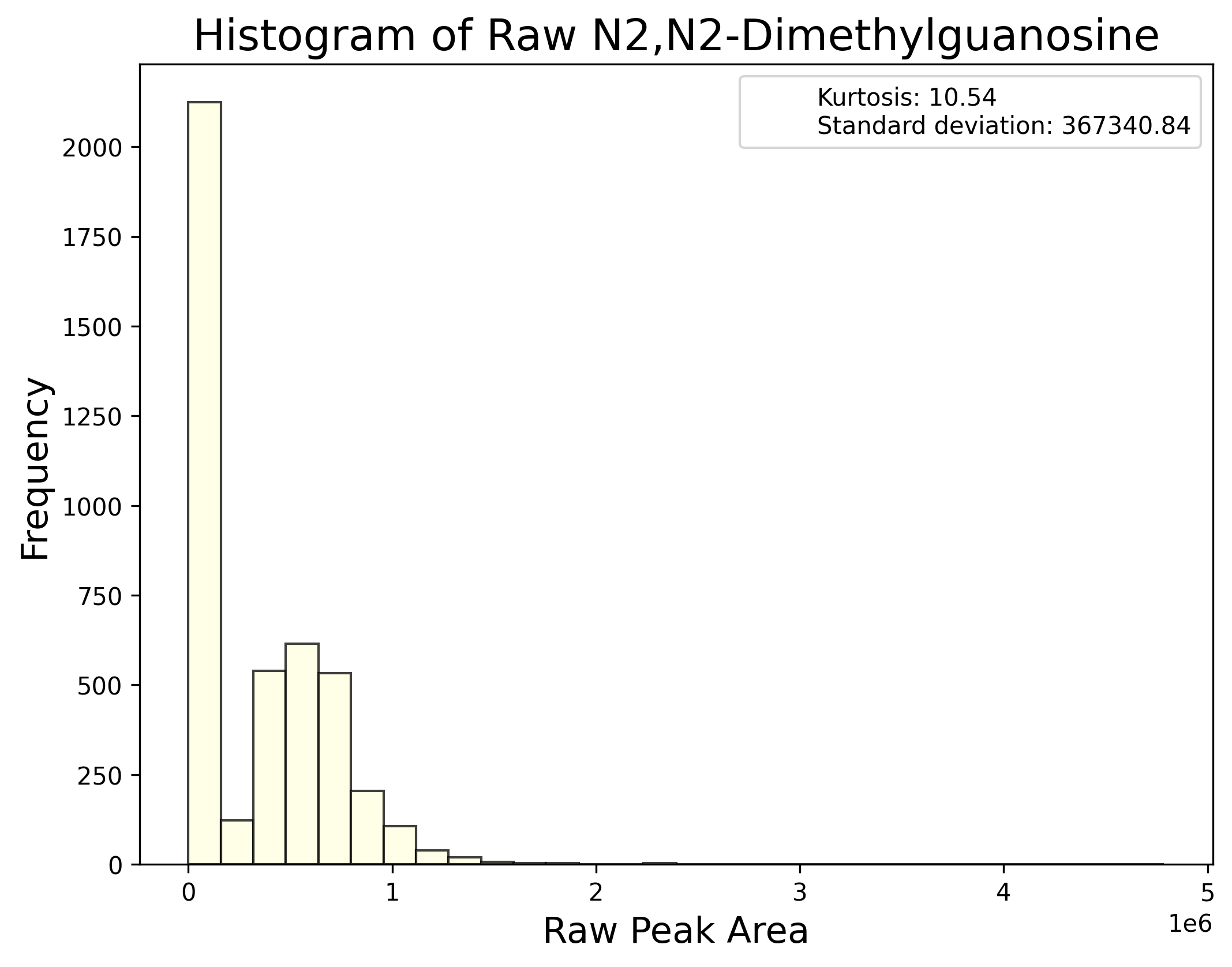

**Supplementary Figure 2. Correlations between metabolic phenotypes.** **a**) Insulin sensitivity (IS) & BMI, **b**) IS and total triglyceride (TG), **c)** IS and interleukin 6 (IL6), **d**) BMI and TG, **e**) BMI and IL6, **f**) TG and IL6

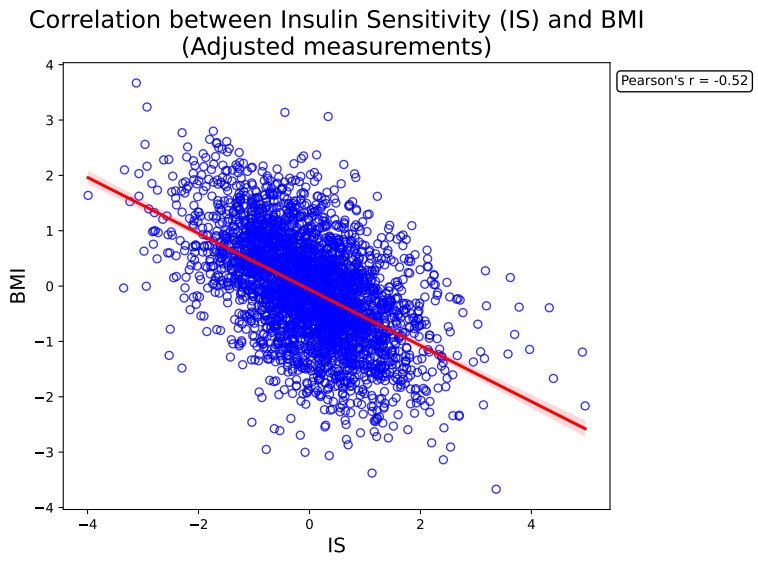

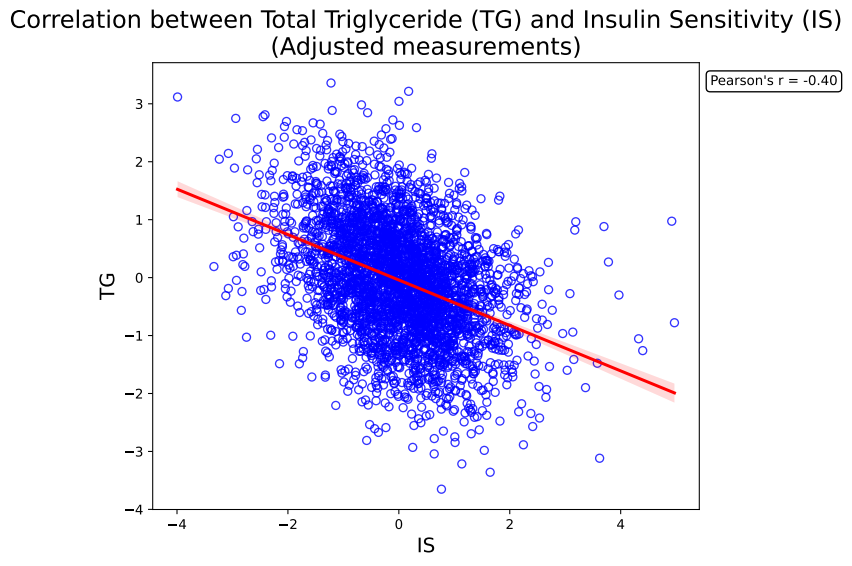

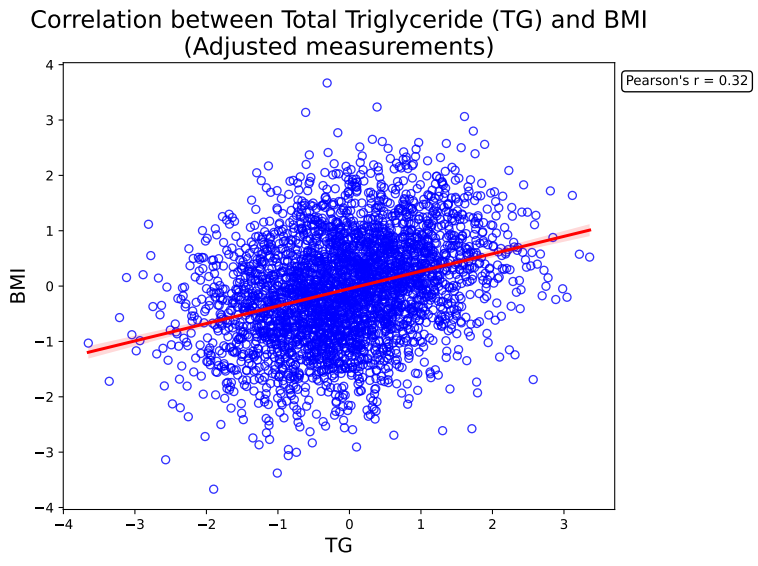

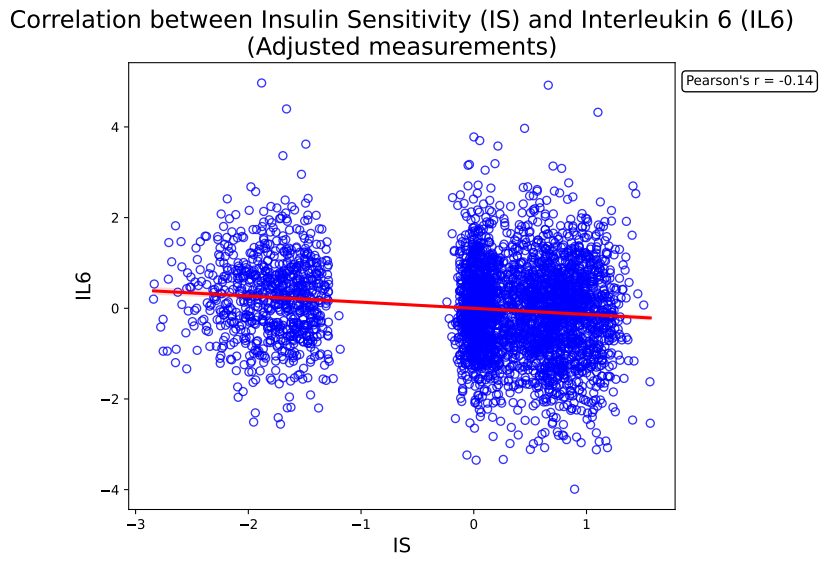

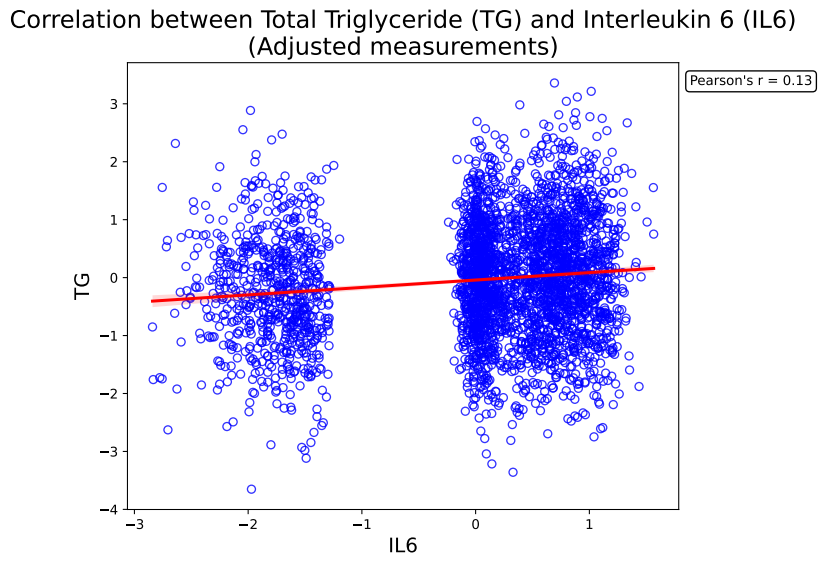

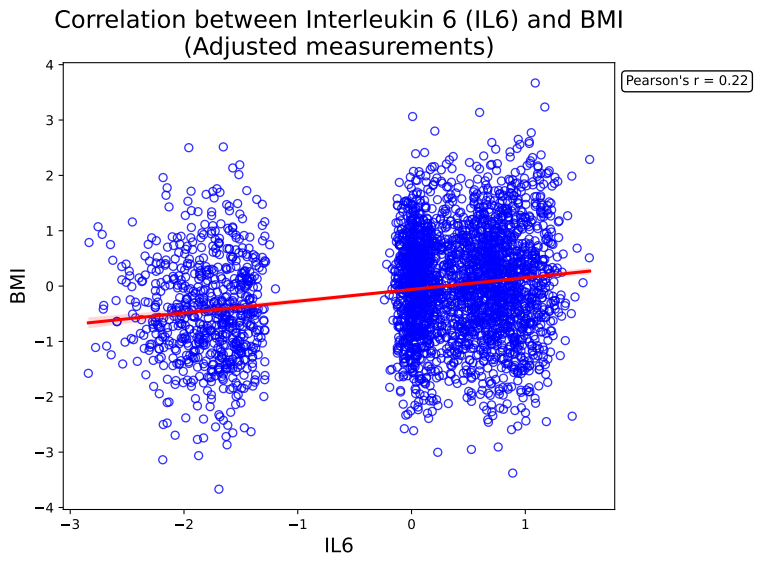

**Supplementary Figure 3. Heritability of lipids in the second clinical visit of the long life family study (LLFS).** Lipids are grouped based on their chemical families.

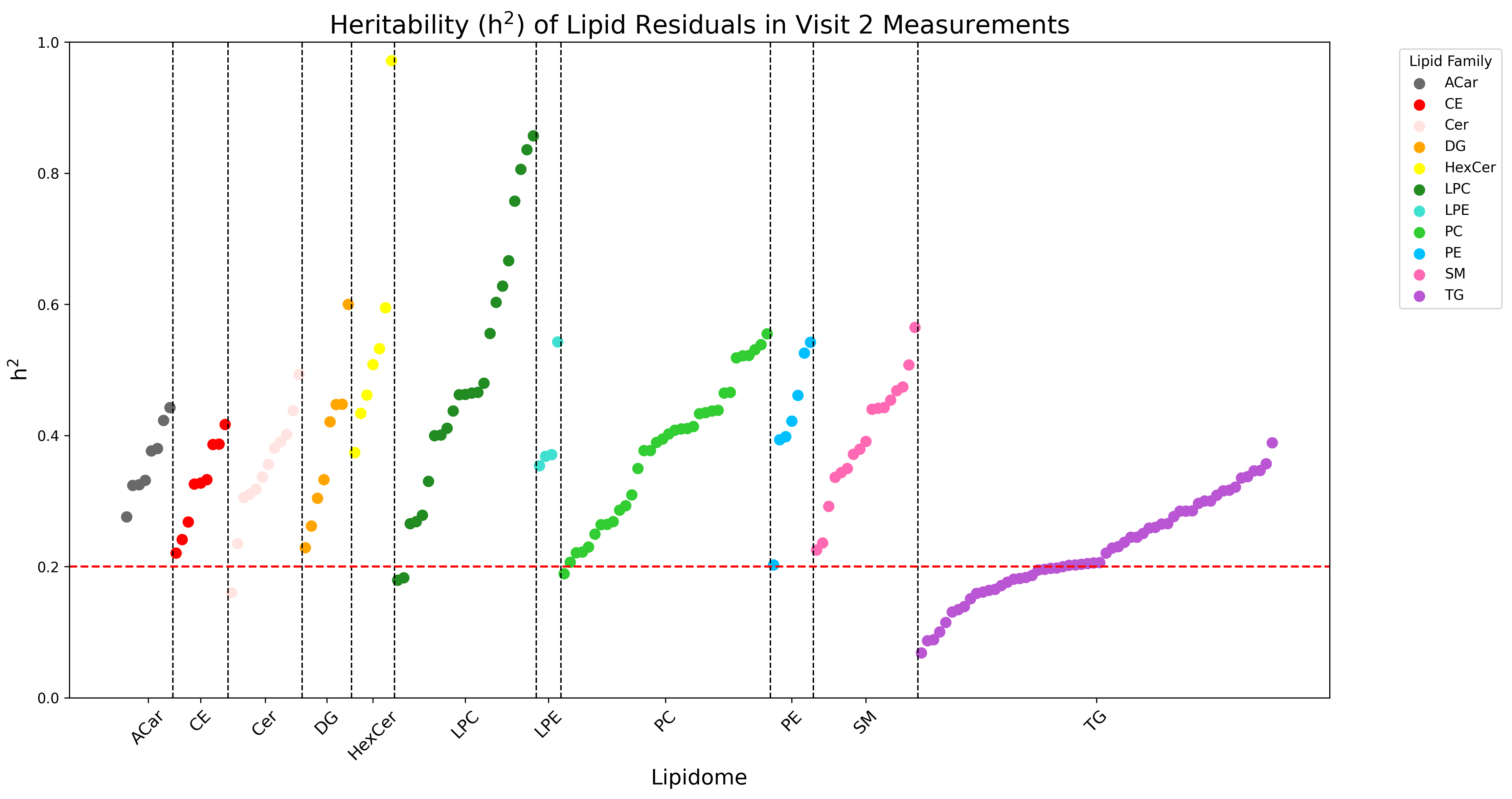

**Supplementary Figure 4. Distribution of the inflaction factors (λ) for gene-metabolome associations.**

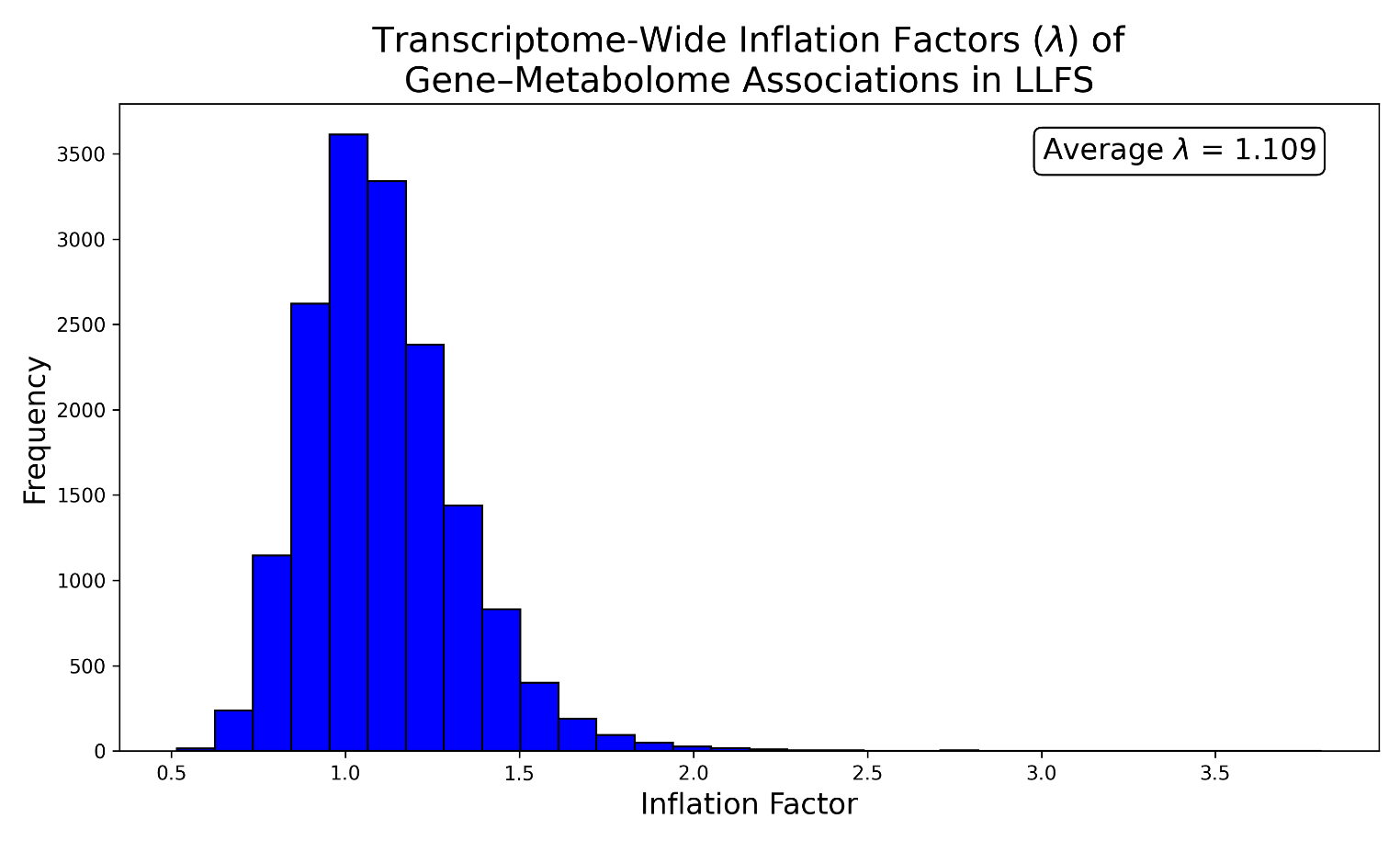

**Supplementary Figure 5. QQ-plots of the “OMICS”-wide association studies for IS.** **a**) transcriptome-wide association study (TWAS), **b)** metabolome-wide association study (MWAS): polar SMs, **c**) MWAS: lipid SMs. All QQ-plots are adjusted for inflation by Bacon.

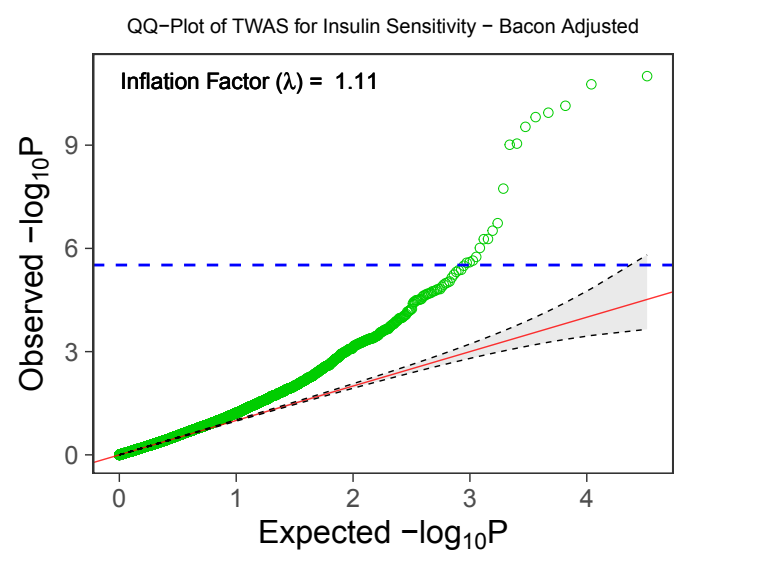

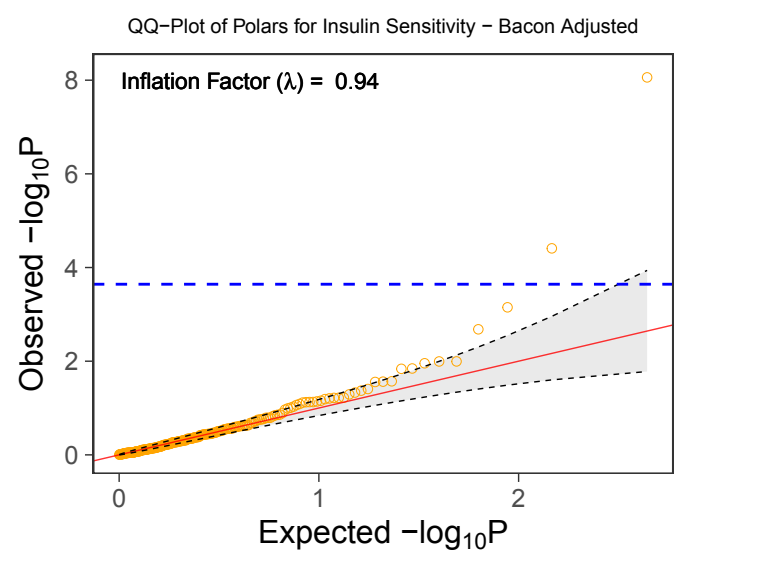

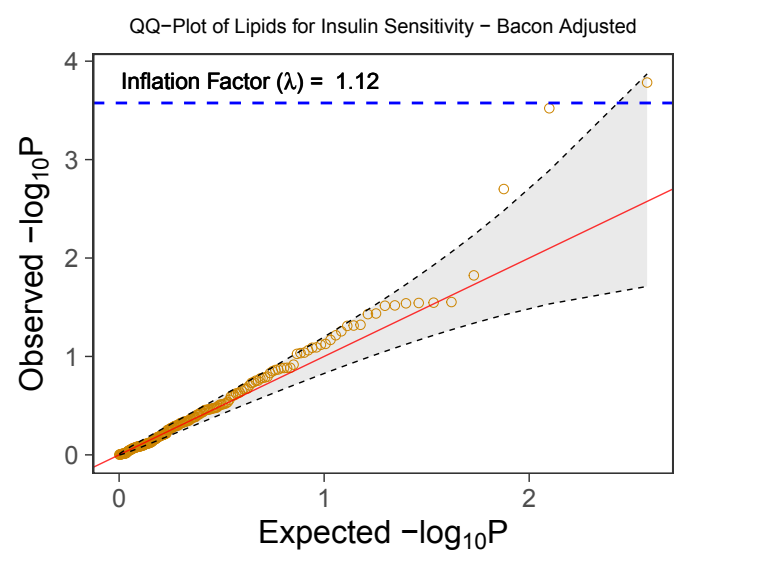

**Supplementary Figure 6. Module composition in multi-modal networks inferred by conventional unsupervised methods**. Module compositions are represented by the proportion of SMs in modules.

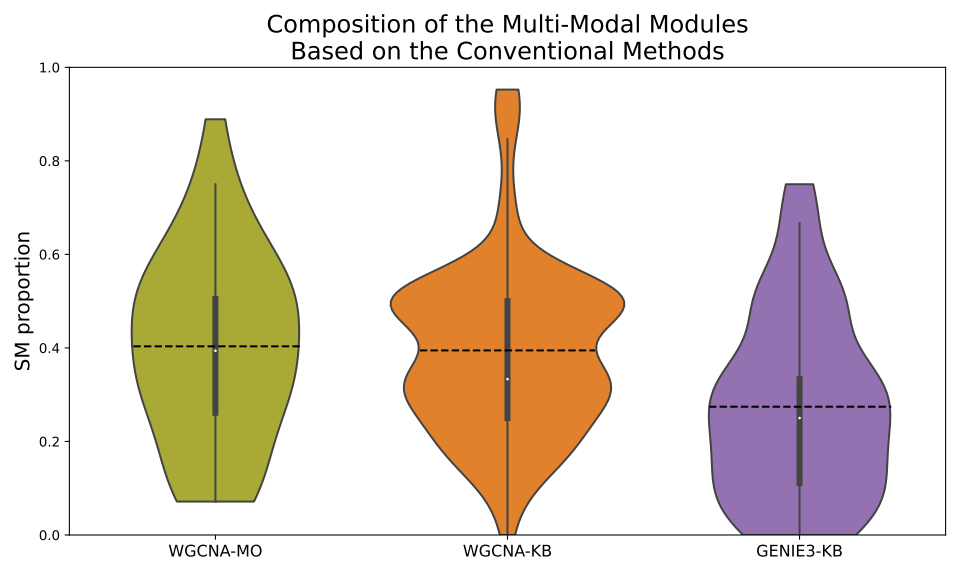

**Supplementary Figure 7**. **Correlation between number of SMs and variance explained by metabolomic principle component 1 (PC1) across GEM-Net modules**.

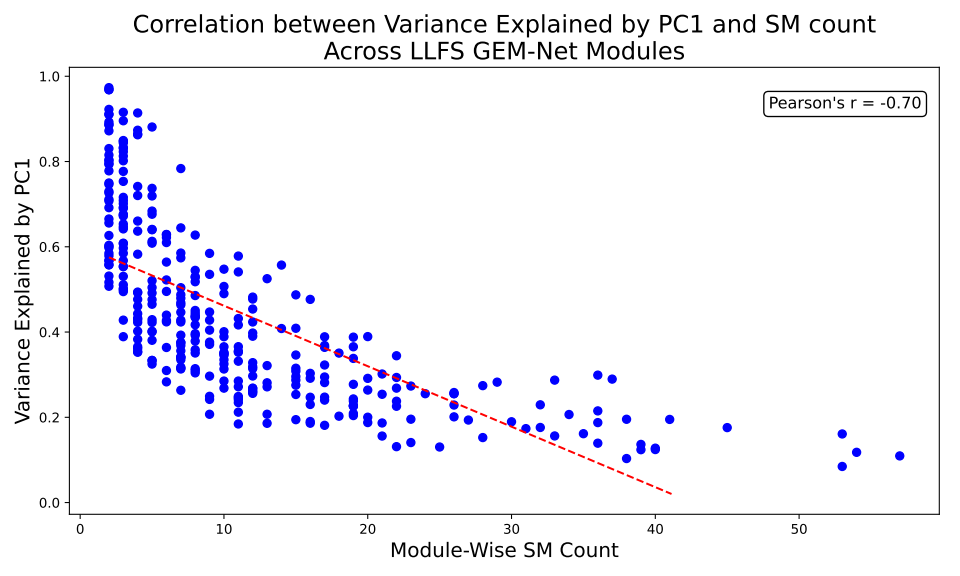

**Supplementary Figure 8. Principle component analysis (PCA) in knowledge-guided GEM-Net modules. a)** Variance explained by metabolomic and transcriptomic PC1 across knowledge-guided GEM-Net modules. **b**) Variance explained by transcriptomic PC1 in GEM-Net modules, grouped by the original source of the baseline gene-level network.

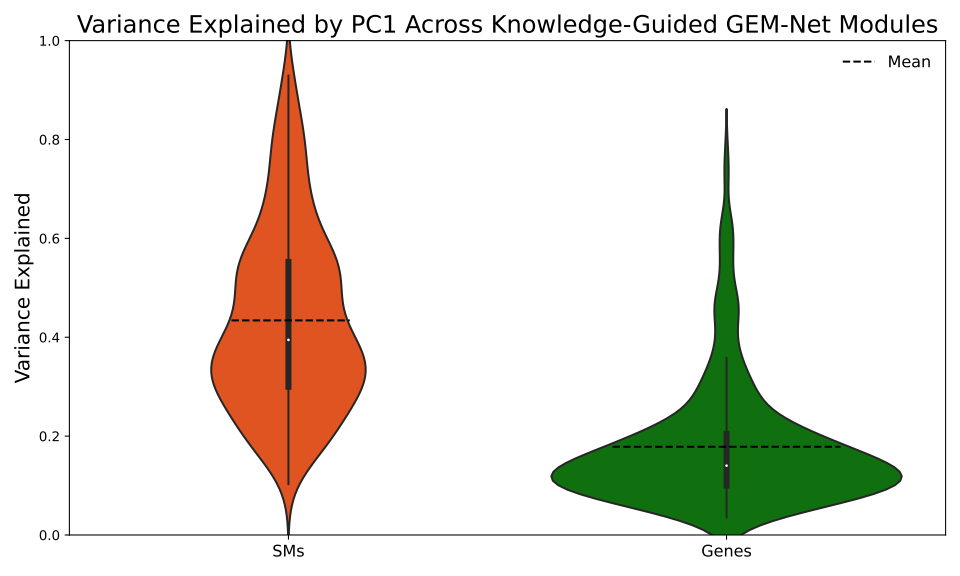

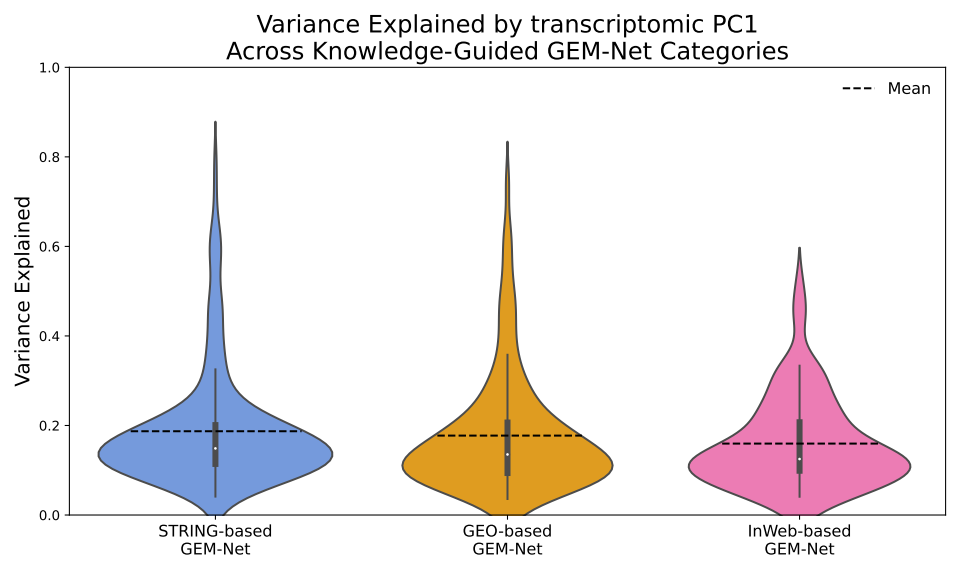

**Supplementary Figure 9. Gene-SM connections in the LLFS GEM-Net.** **a)** Gene-SM connections across modules. **b)** Total number of genes connected to each SM across all modules.

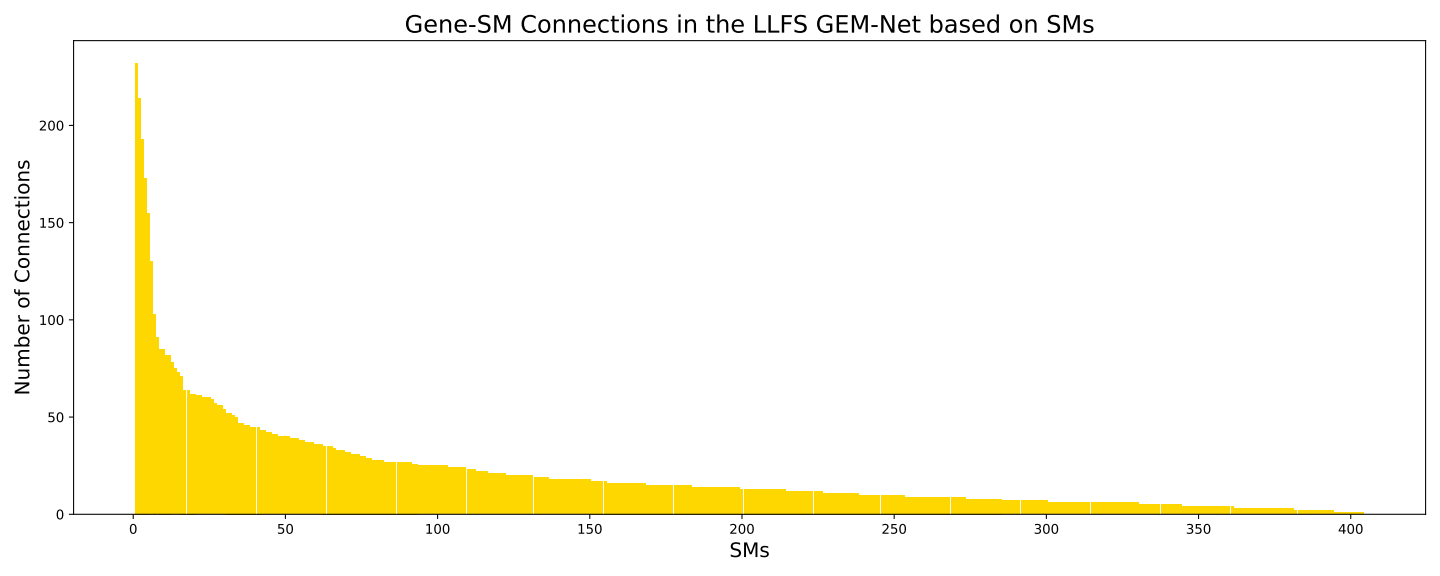

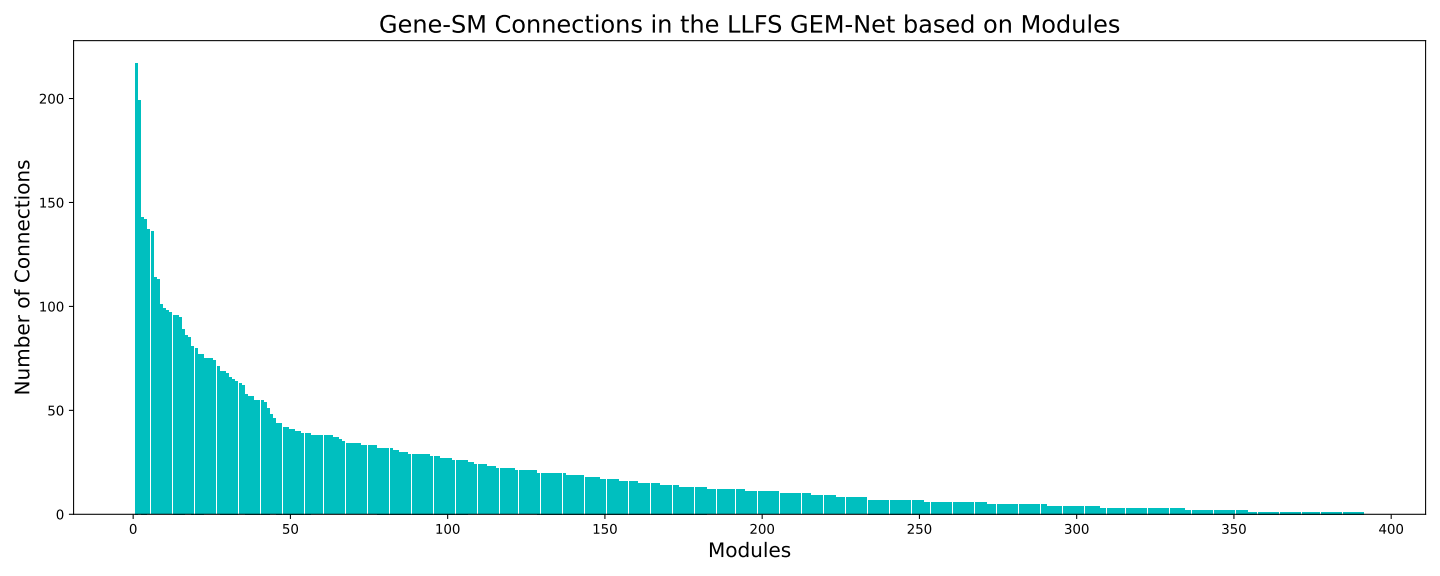

**Supplementary Figure 10. Gene-SM connections in the Knowledge-guided GEM-Net. a)** Gene-SM connections across modules**. b)** Total number of genes connected to each SM across all modules.

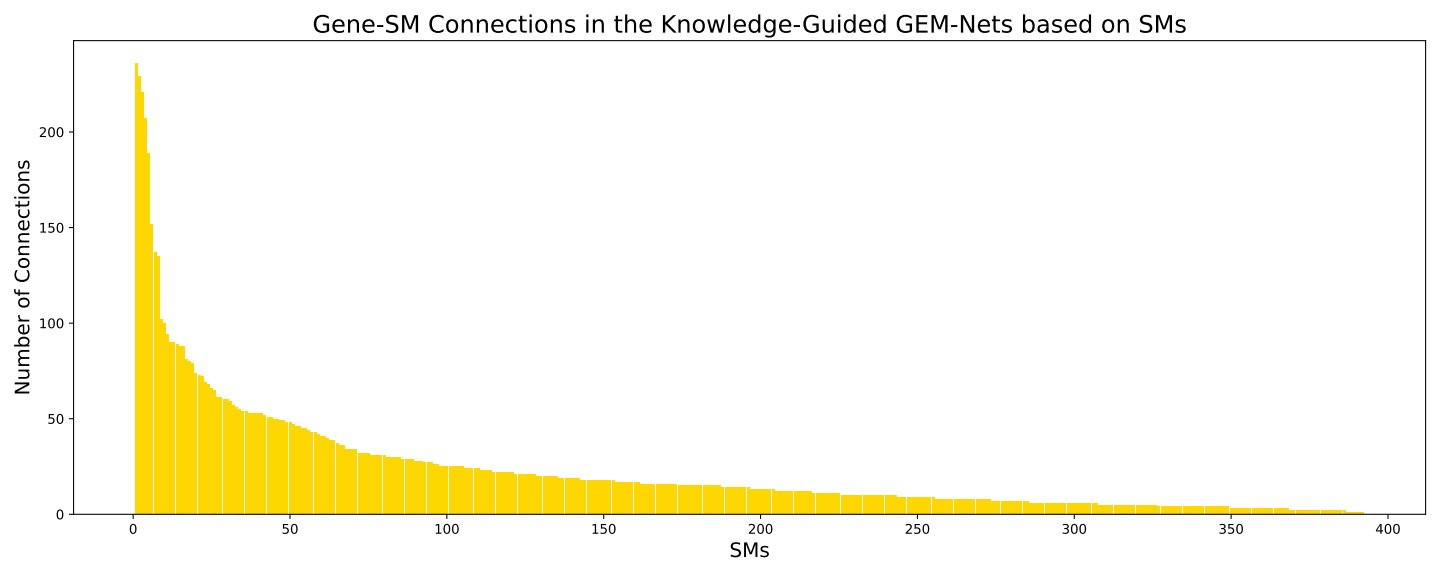

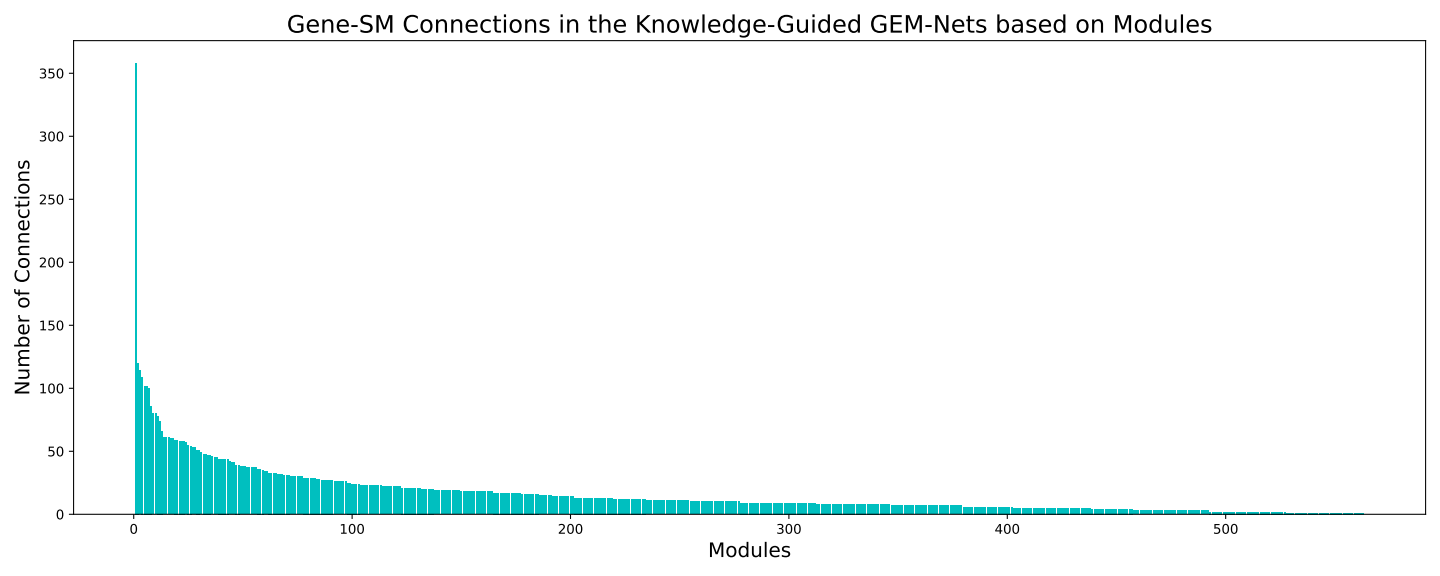

**Supplementary Figure 11. QQ-Plots of the network association tests for IS**. **a**) LLFS GEM-Net modules, **b)** Knowledge-guided GEM-Net modules. Network association tests were performed with Pascal.

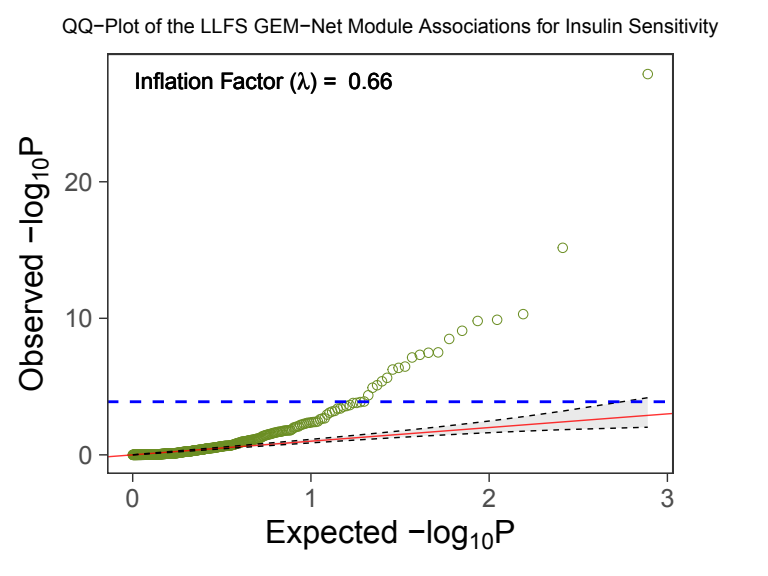

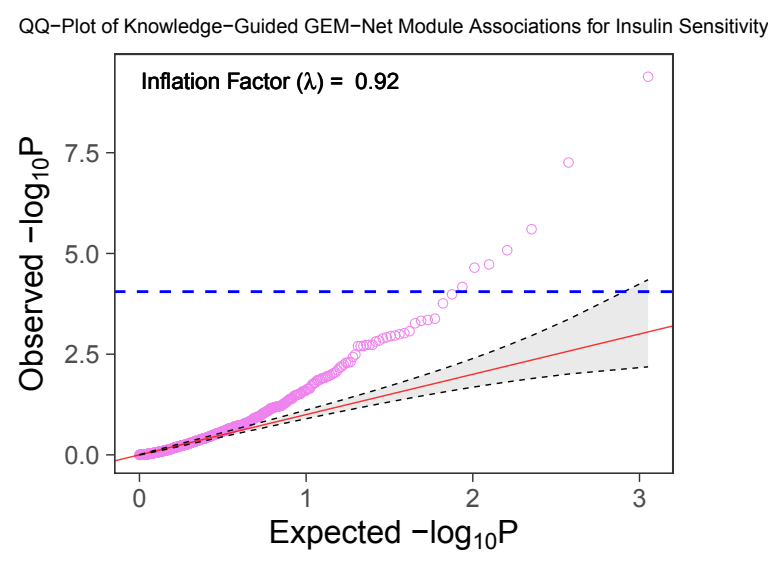

**Supplementary Figure 12.** **P-value distribution of modules in the network association tests for IS**. **a)** LLFS GEM-Net modules, **b)** Knowledge-guided GEM-Net modules. Network association tests were performed with Pascal.

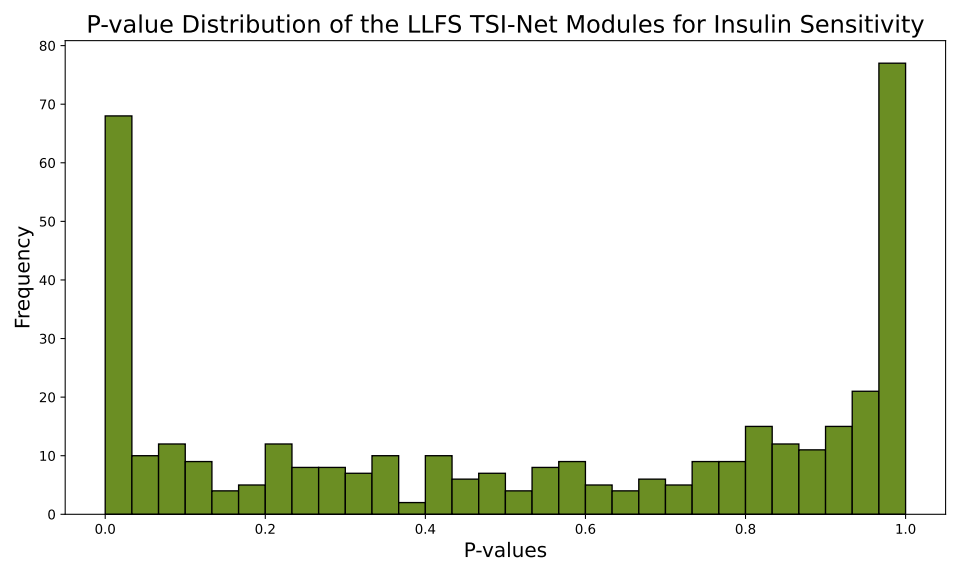

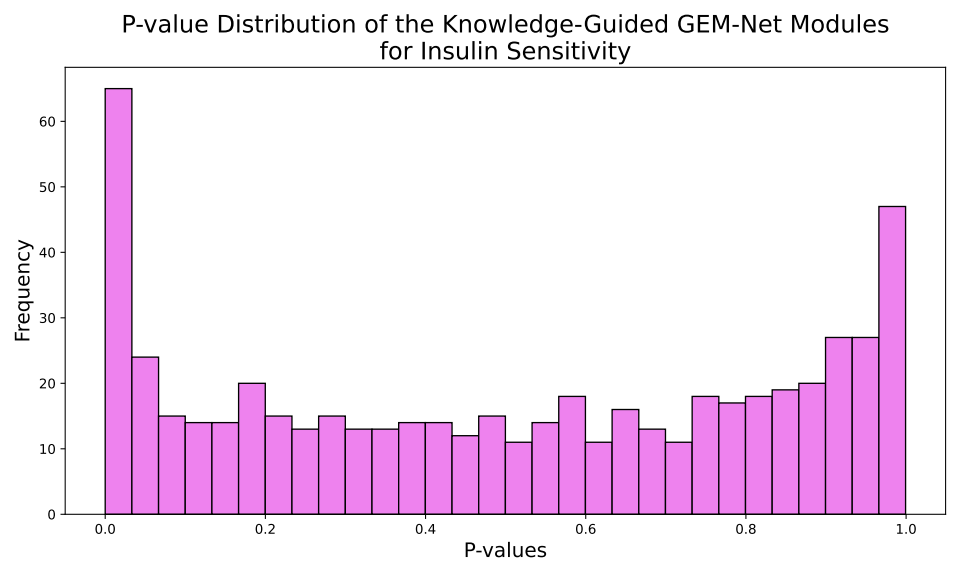
